# The aberrant language network dynamics in autism ages 5–60 years

**DOI:** 10.1101/2024.10.28.620600

**Authors:** Zhe Hu, Xiaolin Guo, Junjie Yang, Zhiheng Qu, Zhongqi Li, Junjing Li, Xiaowei Gao, Jiaxuan Liu, Yaling Wang, Wanchun Li, Wanjing Li, Yien Huang, Jiali Chen, Nan Zhou, Ye Zhang, Xin Wang, Hui Xie, Binke Yuan

## Abstract

Language impairments across both structural components and pragmatic use are frequently observed in individuals with autism spectrum disorder (ASD). These difficulties are thought to stem from atypical brain development and abnormal network interactions, yet an integrative network-level model accounting for such impairments remains lacking. To bridge this gap, we applied the dynamic meta-networking framework of language, a theoretical model capturing domain-segregation dynamics during rest, to examine age-related changes (5-60 years) in cortical language networks in individuals with ASD. To further probe the biological underpinnings of these dynamics, we quantified spatial correspondences between network state hubs and gene co-expression modules as well as neurotransmitter receptor distributions. We identified distinct language meta-states characterized by domain-segregation connectivity patterns, which exhibited spatial alignment with gene co-expression modules and neurotransmitter systems. Individuals with ASD showed state-dependent developmental trajectories marked by age-related hypo- and hyper-connectivity. Critically, these network alterations strongly predicted verbal IQ and communicative difficulties, but were unrelated to social functioning or stereotyped behaviors. Our findings provide novel evidence that language-related network dynamics in ASD are developmentally altered, biologically grounded, and selectively linked to verbal and communicative impairments. These results advance a network-level model of language dysfunction in ASD and highlight potential mechanistic pathways for targeted interventions.

**Lay Summary:** Language challenges are common in autism and can affect both learning language and using it in everyday conversations. In people aged 5-60, we found distinct, age-changing patterns in how the brain’s language areas connect at rest; these patterns align with basic biology (genes and brain chemicals) and strongly relate to verbal ability and daily communication, but not to social difficulties or repetitive behaviors. These results suggest clear, brain-based targets for earlier and more tailored support for language problems in autism.

## Introduction

Autism spectrum disorder (ASD) is a complex neurodevelopmental condition marked by ongoing challenges in social communication and interaction, along with restricted and repetitive behaviors, interests, or activities. Language impairments, including aspects of structural language and pragmatics, frequently occur alongside ASD(Asghari et al., 2021; Brignell, Morgan, et al., 2018; Félix et al., 2024; Gernsbacher et al., 2016; Schaeffer et al., 2023; Tager-Flusberg et al., 2005). The patterns of language impairment in ASD show considerable variability, affecting areas such as phonology, syntax, semantics, lexical strength, and pragmatics(Friedman & Sterling, 2019; Reindal et al., 2023). Three primary profiles of language deficits have been identified in individuals with ASD: (i) minimally verbal autistic individuals, (ii) verbal autistic individuals with structural language impairments(Schaeffer et al., 2023), and (iii) verbal autistic individuals without structural language impairments but with pragmatic impairments. Notably, pragmatic impairment is common in the majority of those with ASD(Andrés-Roqueta & Katsos, 2017; Kalandadze et al., 2018, 2019; Reindal et al., 2023; Schaeffer et al., 2023; Tesink et al., 2009; Williams, 2021) and often presents as difficulties in social communication and interaction(Blume et al., 2021).

A substantial body of evidence shows that language and pragmatic impairments in individuals with ASD are associated with atypical development and abnormal interaction within the language network(Cermak et al., 2022; Dichter, 2012; Duvall et al., 2023; Goodwill et al., 2023; Herringshaw et al., 2016; Kuperberg et al., 2000; Larson et al., 2025; M. Li et al., 2022; Philip et al., 2012; Xiao et al., 2022). The affected brain regions are distributed across bilateral areas of the frontal, temporal, and parietal cortices, which are involved in language. Disruptions within the language network are bidirectional, involving both hyper-connectivity and hypo-connectivity. However, several limitations of current studies should be addressed. First, although language network abnormalities are linked to behavior and clinical features, there is a lack of a theoretical framework to explain these abnormalities. The language system includes domain-specific subsystems(Duffau et al., 2014; Hickok, 2022; Yuan, Xie, Wang, et al., 2023), and different language domains can be affected to varying degrees in individuals with ASD(Schaeffer et al., 2023). Beyond mapping abnormal language regions, a detailed analysis of domain-specific streamline disruptions could provide more behaviorally relevant and translational insights. Second, many studies focus on a single or mixed-age group, overlooking brain and language development(Anderson et al., 2007; Bal et al., 2020; Brignell, Morgan, et al., 2018; Brignell, Williams, et al., 2018; Chan et al., 2023; Y. Chen et al., 2024; Ernst et al., 2015; Georgiou & Spanoudis, 2021; Gernsbacher et al., 2016; Kissine et al., 2023; Pickles et al., 2014; Prescott et al., 2022; Siller & Sigman, 2008; Thurm et al., 2015). Longitudinal research indicates that the brains of individuals with ASD follow complex developmental paths from overgrowth to undergrowth during adolescence(Courchesne et al., 2011, 2019; Ha et al., 2015; Lee et al., 2021; Prigge et al., 2021; The IBIS Network et al., 2017). Nevertheless, how these abnormal developmental trajectories influence the development of language networks across various domains remains largely unclear. Third, most studies use static functional connectivity, neglecting the time-varying patterns of the language network during rest and tasks. The brain is inherently dynamic and reconfigures over a range of timeframes from milliseconds to years(Betzel & Bassett, 2017; Hutchison et al., 2013; Yuan, Xie, Gong, et al., 2023; Yuan, Xie, Wang, et al., 2023). Static functional connectivity may be less sensitive and underestimate the transient disorganization of networks in individuals with ASD(H. Chen et al., 2017; Falahpour et al., 2016; Fu et al., 2019; Gao et al., 2022; Y. Li et al., 2020; Mash et al., 2019; Xie et al., 2022).

To address these limitations, we employed our recently developed dynamic “meta-networking” framework of language to study abnormal language network dynamics in ASD. This framework, observed in healthy adults, included four recurring states with distinct connectivity patterns, hub distributions, structural bases, and cognitive roles(Yuan, Xie, Gong, et al., 2023; Yuan, Xie, Wang, et al., 2023). The first three states relate to language cognition and follow a domain-specific pattern. The first state, characterized by densely connected regions in the superior temporal gyrus (STG), is crucial for perceiving phonemes and processing lexical-phonological information, known as the central stream. The second state, characterized by dense connections in the inferior frontal cortex, is crucial for speech production and is known as the dorsal stream. The third state, characterized by densely connected areas in the inferior parietal cortex, superior and middle temporal gyrus (MTG), is crucial for semantic processing, known as the ventral stream. The fourth state, although it occurs most frequently over time, has weak connectivity and serves as the baseline state. Together, these four states constitute the dynamic “meta-networking” framework of language. The concept of meta-networking refers to a network of networks, aligning with the “meta-networking” theory of cerebral functions(Herbet & Duffau, 2020). According to this theory, complex cognition and behaviors, such as language, emerge from the spatiotemporal integration of distributed yet specialized sub-networks. Our dynamic “meta-networking” framework captures the domain-specific nature of language processing, aligning with the neurobiology of the dual-stream model of speech and language processing(Duffau et al., 2014; Hickok, 2022; Hickok & Poeppel, 2007), and has demonstrated cognitive and clinical relevance (Yuan, Xie, Gong, et al., 2023; Yuan, Xie, Wang, et al., 2023).

In this study, we analyzed resting-state fMRI data from 624 individuals with autism spectrum disorder (ASD) (198 children, 260 adolescents, and 166 adults) and 866 age-matched neurotypical controls (349 children, 293 adolescents, and 224 adults) from the Autism Brain Imaging Data Exchange (ABIDE). Our goal was to address three main questions. First, we examined the developmental trajectory of domain-related dynamics within the cortical language network across healthy children, adolescents, and adults to determine the exact age at which this dynamic pattern appears and stabilizes. Second, we explored the neurochemical and transcriptional bases underlying domain-segregated language dynamics, specifically whether spatial segregation is also reflected in the distribution of gene expression and neurotransmitter receptor density across cortical language regions. Lastly, we investigated how these domain-segregated language dynamics differ in individuals with ASD and evaluated their relationships with both verbal and non-verbal symptom dimensions.

## Materials and Methods

### Participants

The ABIDE initiative is a multi-site open-data resource aimed at speeding up research on the neural bases of autism. It includes two extensive collections: ABIDE I and ABIDE II. These collections feature data from over 24 international brain imaging labs, totaling about 1026 individuals with ASD and 1130 age-matched Healthy Controls (HC). Each dataset provides resting-state fMRI data, along with corresponding structural MRI scans and detailed phenotypic information(Di Martino et al., 2014, 2017).

We conducted our analysis on resting-state fMRI data from all sites. Within each site, we applied the following exclusion criteria to screen resting-state fMRI data: 1) Left-handedness; 2) Subjects with no Intelligence Quotient (IQ) or IQ below 70; 3) excessive head motion (maximum translation > 5 mm or rotation > 5°, or mean framewise displacement [FD] > 0.5 mm); 4) Images that do not cover the entire brain or temporal lobe; 5) Low spatial signal-to-noise ratio (below the third standard deviation of the group), especially in temporal, frontal, and parietal cortices; We also excluded sites with fewer than 10 participants remaining in either the ASD or control group after screening. Following these criteria, a total of 624 individuals with ASD (198 children aged 5-11 years, 260 adolescents aged 12-18 years, and 166 adults aged 19-60 years) and 866 age-matched healthy controls (349 children, 293 adolescents, and 224 adults) were included in the study. Demographic information, IQ scores (full scale IQ [FSIQ], verbal IQ [VIQ], and performance IQ [PIQ]), and Autism Diagnostic Observation Schedule (ADOS) scores are summarized in Table 1 and Supplementary Table 1.

**Table 1.** Demographic information and clinical deficits.

| Characteristics<br>Mean ± SD | Total |  |  | Children (5-11) |  |  | Adolescents (12-18) |  |  | Adults (19-60) |  |  |
| --- | --- | --- | --- | --- | --- | --- | --- | --- | --- | --- | --- | --- |
|  | ASD<br>n = 624 | HC<br>n = 866 | p-value | ASD<br>n = 198 | HC<br>n = 349 | p-value | ASD<br>n = 260 | HC<br>n = 293 | p-value | ASD<br>n = 166 | HC<br>n = 224 | p-value |
| Gender<br>(Female/Male) | 81 / 543 | 214 / 652 | < **0.01<sup>a</sup>** | 33 / 165 | 112 / 237 | < **0.01<sup>a</sup>** | 30 / 230 | 63 / 230 | < **0.01<sup>a</sup>** | 18 / 148 | 39 / 185 | 0.07 |
| Age, Years<br>[MIN, MAX] | 16.59 ± 8.91<br>[5.22, 59] | 15.98 ± 8.58<br>[5.89, 64] | 0.19 | 9.76 ± 1.56<br>[5.22, 11.99] | 9.76 ± 1.29<br>[5.89, 11.99] | 0.96 | 14.73 ± 1.92<br>[12.02, 18.92] | 14.65 ± 1.97<br>[12, 18.90] | 0.64 | 27.64 ± 10.33<br>[19, 59] | 27.40 ± 9.15<br>[19, 64] | 0.82 |
| FSIQ | 106.92 ± 15.08<br>n = 624 | 113.27 ± 12.09<br>n = 866 | < **0.01<sup>b</sup>** | 105.34 ± 15.86<br>n = 198 | 114.93 ± 12.24<br>n = 349 | < **0.01<sup>b</sup>** | 105.06 ± 14.77<br>n = 260 | 110.52 ± 11.97<br>n = 293 | < **0.01<sup>b</sup>** | 110.71 ± 13.57<br>n = 166 | 114.27 ± 11.43<br>n = 224 | 0.05 |
| VIQ | 106.24 ± 16.23<br>n = 474 | 114.08 ± 13.30<br>n = 656 | < **0.01<sup>b</sup>** | 105.53 ± 16.28<br>n = 149 | 116.85 ± 14.03<br>n = 252 | < **0.01<sup>b</sup>** | 104.90 ± 16.38<br>n = 216 | 111.35 ± 13.04<br>n = 268 | < **0.01<sup>b</sup>** | 109.87 ± 15.44<br>n = 109 | 114.35 ± 11.25<br>n = 136 | **0.01<sup>b</sup>** |
| ADOS Social | 7.59 ± 2.70<br>n = 371 | — | — | 7.53 ± 2.58<br>n = 107 | — | — | 7.68 ± 2.88<br>n = 132 | — | — | 7.55 ± 2.63<br>n = 132 | — | — |
| ADOS Comm | 3.60 ± 1.51<br>n = 364 | — | — | 3.27 ± 1.37<br>n = 103 | — | — | 3.64 ± 1.63<br>n = 129 | — | — | 3.83 ± 1.46<br>n = 132 | — | — |
| ADOS RRB | 2.32 ± 1.32<br>n = 265 | — | — | 2.48 ± 1.39<br>n = 85 | — | — | 2.45 ± 1.41<br>n = 82 | — | — | 2.07 ± 1.16<br>n = 98 | — | — |

<sup>a</sup> P-value obtained using Pearson's chi-square test.
<sup>b</sup> P-value obtained using a two-sample t-test.
ASD, Autism spectrum disorder; HC, Healthy control; SD, Standard deviation; FSIQ, full scale intelligence quotient; VIQ, verbal intelligence quotient; PIQ, performance intelligence quotient; ADOS, Autism Diagnostic Observation Schedule; ADOS Comm, ADOS Communication; ADOS
RRB, ADOS Restricted interests and repetitive behavior.

### Data preprocessing

The resting-state fMRI data were preprocessed with the following steps: 1) delete the first 10 volumes; 2) slice timing correction; 3) head motion correction; 4) co-registration; 5) T1 segmentation and normalization into MNI space using DARTEL; 6) functional normalization with the deformation field of T1 images; 7) smoothing with a Gaussian kernel (full-width-at-half-maximum = 6 mm); 8) linear detrending; 9) regression of nuisance signals (24 parameters, including x, y, z translations and rotations (6 parameters), their temporal derivatives (6 parameters), and quadratic terms of 12 parameters); 10) temporal band-pass filtering (0.01–0.1 Hz).

### Language network definition

The putative language network was defined using the Human Brainnetome Atlas(L. Fan et al., 2016). We extracted all parcels that showed significant activation in language-related behavioral domains (i.e., language or speech) or paradigm classes (e.g., semantic, word generation, reading, or comprehension). Thirty-three cortical parcels in the left hemisphere and 15 cortical parcels in the right hemisphere were selected. Their extents are very similar to those identified by other researchers(Fedorenko et al., 2010; Lipkin et al., 2022; Lombardo et al., 2018; Rampinini et al., 2017; Vigneau et al., 2006). Considering bilateral language processing(Chai et al., 2016; Hodgson et al., 2021; Lerner et al., 2011; Muller & Meyer, 2014; Rice et al., 2015; Siegel et al., 2016) and the recruitment of right hemisphere language regions for compensation in clinical populations, 19 homologous parcels in the right hemisphere and one homologous parcel in the left hemisphere were also included. In total, a symmetric language network comprising 68 cortical ROIs (34 in each hemisphere) was defined(Yuan et al., 2022; Yuan, Xie, Wang, et al., 2023), including the superior, middle, and inferior frontal gyrus (SFG, MFG, and IFG, respectively), the ventral parts of the precentral gyrus (PrG) and postcentral gyrus (PoG), the STG, MTG, and inferior temporal gyrus (ITG), the fusiform gyrus (FuG), the parahippocampal gyrus (PhG), and the posterior superior temporal sulcus (pSTS). The coordinates (in Montreal Neurological Institute space) and meta-analysis results for each parcel are summarized in Supplementary Table 2.

### Frame-wise time-varying language network construction using DCC

To identify the recurring temporal states of the language network, the dynamic conditional correlation (DCC) approach was adopted to construct the framewise time-varying language network(Yang et al., 2025) (https://github.com/yuanbinke/Naturalistic-Dynamic-Network-Toolbox). It has been demonstrated that DCC outperformed the standard sliding-window approach in tracking network dynamics(Choe et al., 2017; Lindquist et al., 2014).

DCC is a variation of the multivariate GARCH (generalized autoregressive conditional heteroskedasticity) model(Engle, 1982; Lebo & Box-Steffensmeier, 2008), which has proven to be especially effective for estimating both time-varying variances and correlations. GARCH models describe the conditional variance of a single time series at time t as a linear combination of past values of the conditional variance and the squared process itself. All DCC parameters are estimated using quasi-maximum likelihood methods and do not require any ad hoc parameter settings.

The DCC algorithm consists of two steps. To illustrate, let us assume that there is a pair of time series from two ROIs, *x_t_* and *y_t_*. In the first step, standardized residuals of each time series are estimated using a univariate GARCH (1,1) process. In the second step, an exponentially weighted moving average (EWMA) window is applied to the standardized residuals to compute a non-normalized version of the time-varying correlation matrix between *x_t_* and *y_t_*. The mathematical expressions of the GARCH (1,1) model, DCC model, and EWMA, and the estimations of the model parameters were provided by Lindquist et al.(Lindquist et al., 2014).

K-means clustering was used to decompose the dFC matrices into several recurring connectivity states. The optimal number of clusters *k* was determined using the elbow criterion, which is the ratio of within-cluster distance to between-cluster distances(Allen et al., 2014). The L1 distance function (‘Manhattan distance’) was employed to measure the distance from each point to the centroid. Each time window was ultimately assigned to one of these connectivity states.

### Topological properties of dFC states

For each subject, a state-specific connectivity matrix was estimated by taking the median of all connectivity matrices assigned to the same state label(Damaraju et al., 2014). Before calculation, a correlation threshold (*r* > 0.2) was applied to remove weak connections that could have originated from noise. Because negative connections are debatable(Murphy & Fox, 2017), only positive connections were included in this work. In this study, we used a weighted network instead of a binarized one to preserve all connectivity information. Given the heterogeneous lesion locations, we focused only on changes in nodal and global topological properties, not on individual connections.

*<u>Global topological metrics</u>*: for each state-specific matrix, three network topological properties were calculated: 1) Total connectivity strength (FC strength). The FC strength of a network was calculated by summing functional connectivity strengths of all suprathreshold connections into one value(Nelson et al., 2017); 2) Global network efficiency (gE). The gE reflects the capability for parallel information transfer and functional integration. It is defined as the average of the inverses for all weighted shortest path lengths (the minimal number of edges that one node must traverse to reach another) in the thresholded matrix(Rubinov & Sporns, 2010); 3) Local network efficiency (lE). The lE reflects the relative functional segregation and is defined as the average of the global efficiency of each node’s neighborhood sub-graph; 4) Rich-club organization. Rich-club organization in a network refers to a phenomenon that the hub nodes are more densely connected among themselves than non-hub nodes. The presence of a rich-club organization provides important information on the higher-order structure of a network and indicates the functional specialization(Van Den Heuvel & Sporns, 2011).

#### Nodal topological metrics

We calculated nodal strength, gE, and lE for each node. We also calculated betweenness centrality (BC), an index indicating whether a particular node lies on the shortest paths between all pairs of nodes in the network.

As a supplementary analysis, we also computed the static functional connectivity (sFC) matrix using the Pearson correlation coefficient. Then, we calculated the three topological properties of sFC and examined the changes in the two patient groups.

### Functional relevance and biological basis of meta states

To evaluate the functional importance of hubs in each state, we conducted term-based meta-analyses using the platform NeuroQuery (https://neuroquery.org/)(Dockès et al., 2020). NeuroQuery predicts the spatial distribution of activity based on 418,772 activation points linked to 7647 terms from the full texts of 13,459 neuroimaging articles. To confirm functional specificity, we analyzed language-related terms such as speech perception, inner speech, phonological processing, speech production, and semantic processing. The functional relevance was measured by calculating the dice coefficients between binary images of hub nodes and meta-analysis results (dice = 2* (hub * meta)/(hub + meta)). Since the meta-analysis showed left-lateralized activations, dice coefficients were computed for both the entire brain and the left hemisphere.

To further elucidate the molecular underpinnings of these network dynamics, we examined the ROI-wise gene expression profiles of cortical language networks. Gene expression data were obtained from the Allen Human Brain Atlas (AHBA), comprising 3,702 regional samples from six postmortem human brains (http://human.brain-map.org). We then preprocessed the AHBA data using the Abagen toolbox (https://github.com/rmarkello/abagen) according to the published pipeline(ArnatkevicCiūtė et al., 2019). Because the AHBA provides right-hemisphere data for only two donors, analyses were restricted to the left hemisphere. This yielded a 34 × 15,633 matrix of regional gene expression levels, which was used to characterize transcriptional activity.

To elucidate the modular architecture, we first applied z-score normalization to the gene-expression profiles of 34 left-hemisphere ROIs. We then computed a 34 × 34 Pearson correlation matrix, thresholded at |*r*| < 0.1, and removed self-connections to keep only strong co-expression links. Next, we applied the Louvain community-detection algorithm from the Brain Connectivity Toolbox(Rubinov & Sporns, 2010) (http://www.brain-connectivity-toolbox.net) to this weighted network, using the default resolution parameter (*γ* = 1.0) and symmetric handling of negative weights. The Louvain method initially assigns each node to its own community. It then iteratively evaluates each node, moving it to the neighboring community that maximizes the modularity gain (Q) until no further improvement occurs. Once local optimization stalls, communities are combined into “super-nodes” to form a reduced network. The process of relocating nodes and aggregating networks repeats until the overall modularity score (Q) converges. This procedure yields a final modularity score (Q) and assigns a community label to each ROI, which is then exported for downstream analyses.

To examine the neurochemical underpinnings of meta-states, we correlated nodal strength in each meta-state with neurotransmitter distribution maps from the JuSpace toolbox (https://github.com/juryxy/JuSpace). The JuSpace toolbox provides distribution maps of multiple neurotransmitter transporters and receptors derived from healthy volunteers, including 5-HT1a (5-hydroxytryptamine receptor subtype 1a), 5-HT1b, 5-HT2a, 5-HT4, CB1 (cannabinoid receptor 1), CBF(cerebral blood flow), D1 (dopamine receptor 1), D2, DAT (dopamine transporter), F-DOPA (Fluorodopa), GABAa (gamma-aminobutric acid receptor a), KappaOp (kappa opioid receptor), MU (μ-opioid receptors), NAT (noradrenaline transporter), NMDA (N-methyl-D-aspartate receptor), SERT (serotonin transporter), VAChT (Vesicular acetylcholine transporter), mGluR5 (Metabotropic glutamate receptor 5). Spatial Spearman correlations were computed between nodal strength in each meta-state and each selected neurotransmitter map. Statistical significance was assessed via 5,000 permutation tests. For each permutation, we generated a null distribution of correlation coefficients and calculated the empirical p-value as: *P* = (number of permutation tests<actual accuracy+1) / (number of permutation tests+1).

### Statistical analysis

Age, FIQ, VIQ, and PIQ were analyzed using two-sample t-tests. Sex data were analyzed with Pearson’s chi-square test.

We applied ComBat harmonization(Fortin et al., 2017, 2018; Xia et al., 2019) to the dynamic functional connectivity matrices for each meta-state. This approach attenuates unwanted inter-site variability arising from acquisition differences while preserving biologically meaningful between-subject variance. Age and sex were included as covariates in the ComBat model to retain biologically relevant variability.

For each meta-state that had undergone ComBat harmonization, edge strength, nodal, and global topological properties were compared using two-sample t-tests. Edge strength results were corrected using a network-based statistic (NBS)(Zalesky et al., 2010), with an edge *p*-value of 0.01 and a component *p*-value of 0.05. Nodal and global topological property results were corrected using the False Discovery Rate (FDR), with a corrected *p*-value of 0.05.

The associations between edge strength and language deficits in individuals with ASD (i.e., VIQ and ADOS communication scores) were evaluated by calculating the partial Pearson correlation coefficient after controlling for sex, age, and FSIQ. The results were adjusted using NBS with a component *p*-value of 0.05.

### Machine learning-based dFC-behavior prediction model

To investigate whether the reorganized dFCs were linked to autistic language deficits, we built multivariate machine learning-based dFC-deficit prediction models. The relevance vector regression (RVR) algorithm and linear kernel function were used(Cui & Gong, 2018; Yuan et al., 2019). RVR has no algorithm-specific parameters and therefore does not require extra computational resources to estimate the optimal parameters(Tipping, 1999).

#### Linear relevance vector regression (RVR)

RVR is a Bayesian framework for learning sparse regression models. In RVR, only some samples (smaller than the training sample size), termed the ‘relevance vectors’, are used to fit the model:

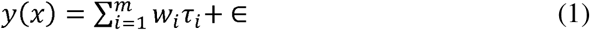

where τ*_i_* are basis functions, ∊ is normally distributed with mean 0 and variance *β*. RVR uses training data to build a regression model:

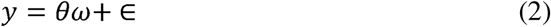

where *y = [y*_1_, *y*_2_*,… y_n_]^T^*, *θ* = [*θ*_1_, *θ*_2_,… *θ_n_*], *θ_i_* = [*τ*_i_*(x*_1_), *τ_i_*(*x*_1_), … *τ_i_*(*x_n_*)]*^T^*. Each vector *θ_i_*, consisting of the values of the basis function τ*_i_* for the input vectors, is a relevance vector.

The model parameters *β* were found by using the maximum likelihood estimates from the conditional distribution: *p(y | α, β)* = 𝒩(*y* | 0, 𝒞), where the 𝒞 = *βI_n_* + Φ*A*^-1^Φ*^T^*. To make the RVM favor sparse regression models, prior distributions were assumed for both *w_i_* and *β*^-1^, i.e., *p*(*w_i_* | *α_i_*) = 𝒩(0, *α_i_*^-1^). The ways of treating priors, however, lead to the same relevance vector machine construction.

#### Prediction accuracy and significance

Leave-one-out-cross-validation (LOOCV) was used to calculate the prediction accuracy (the Pearson correlation coefficient between the predicted and actual labels). In each turn of the LOOCV, one patient was designated as the test sample, and the remaining patients were used to train the lesion model. The predicted score was then obtained by the feature matrix of the tested sample.

The significance level was computed based on 1000 permutation tests. For each permutation test, the prediction labels (e.g., verbal IQ) were randomized, and the same RVR prediction process as used in the actual data was carried out. After 1000 permutations, a random distribution of accuracies was obtained and the *P* value was correspondingly calculated: 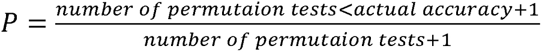.

Features of each prediction model were based on the subject’s median metric across all states. As a complementary analysis, we also constructed an sFC-behavior prediction model.

### Validation analysis

In our validation analysis, we incorporated both internal and external validation strategies. For internal validation, we used data from the NYU site, which contributed the largest sample to the ABIDE cohort. For external validation, we selected imaging data from two independent sites to assess the robustness of our findings.

Specifically, our external validation study included a total of 490 participants (280 females; mean age = 16.84, SD = 13.91) from two independent sites. The first validation dataset was the Chinese Color Nest Project (CCNP: http://deepneuro.bnu.edu.cn/?p=163). This dataset primarily comprised Chinese children and adolescents recruited from the Chongqing and Beijing regions. A total of 479 participants aged 6 to 17.9 years were initially included. After quality control using the same exclusion criteria, 297 participants were retained for analysis (167 females; mean age = 10.67, SD = 3.16). Participants had no neurological or psychiatric disorders and were not taking psychotropic medication. During the resting-state fMRI scan, participants were instructed to fixate on a light crosshair or a cartoon image presented on a dark screen, stay still, and refrain from engaging in any specific thoughts(X.-R. Fan et al., 2023). This project was approved by the Institutional Review Board of the Institute of Psychology, Chinese Academy of Sciences (ethical approval number: H18017). Detailed scanning parameters are provided in the Supplementary Table 3. The second validation site recruited 233 participants. After quality control, 192 participants were retained for analysis (113 females; mean age = 26.35, SD = 18.10). All participants were required to be in good health, with no history of psychiatric disorders, substance abuse (including illicit drug or alcohol abuse), or contraindications to MRI. Resting-state fMRI data were acquired at Southwest University Hospital, Chongqing. During the scan, participants were instructed to keep their eyes closed, remain still, and stay awake(Wei et al., 2018). Detailed scanning parameters are provided in the Supplementary Table 4.

For both external and internal validation, we adopted the same preprocessing pipeline and ‘meta-networking’ framework as that used in the ABIDE cohort. Furthermore, because the external validation dataset involved multiple sites, we applied ComBat harmonization to the dynamic functional connectivity matrices while preserving group, age, and sex effects as covariates.

## Results

### Demographic and clinical characteristics

Table 1 summarizes the demographic, language, and clinical information of the participants. Males with ASD were dominant in all three groups. The FSIQ and VIQ were significantly lower in the ASD group across all three groups.

### The temporally recurring states of the language network in resting state

In the six groups of HCs and ASDs, four recurring states were identified (Fig. 1). Consistent with the dynamic “meta-networking” framework of language, these four states exhibited distinct functional connectivity patterns and hub distributions. State 1 was characterized by moderate to high positive connectivity between nodes in the bilateral STG, PrG, and PoG, but weak or moderate negative connectivity between prefrontal and temporal nodes. In state 2, strong positive connectivity was observed among prefrontal nodes. State 3 stood out from States 1 and 2 due to its strong connectivity among temporal nodes. State 4 showed an overall weak connectivity pattern.

**Figure 1.**
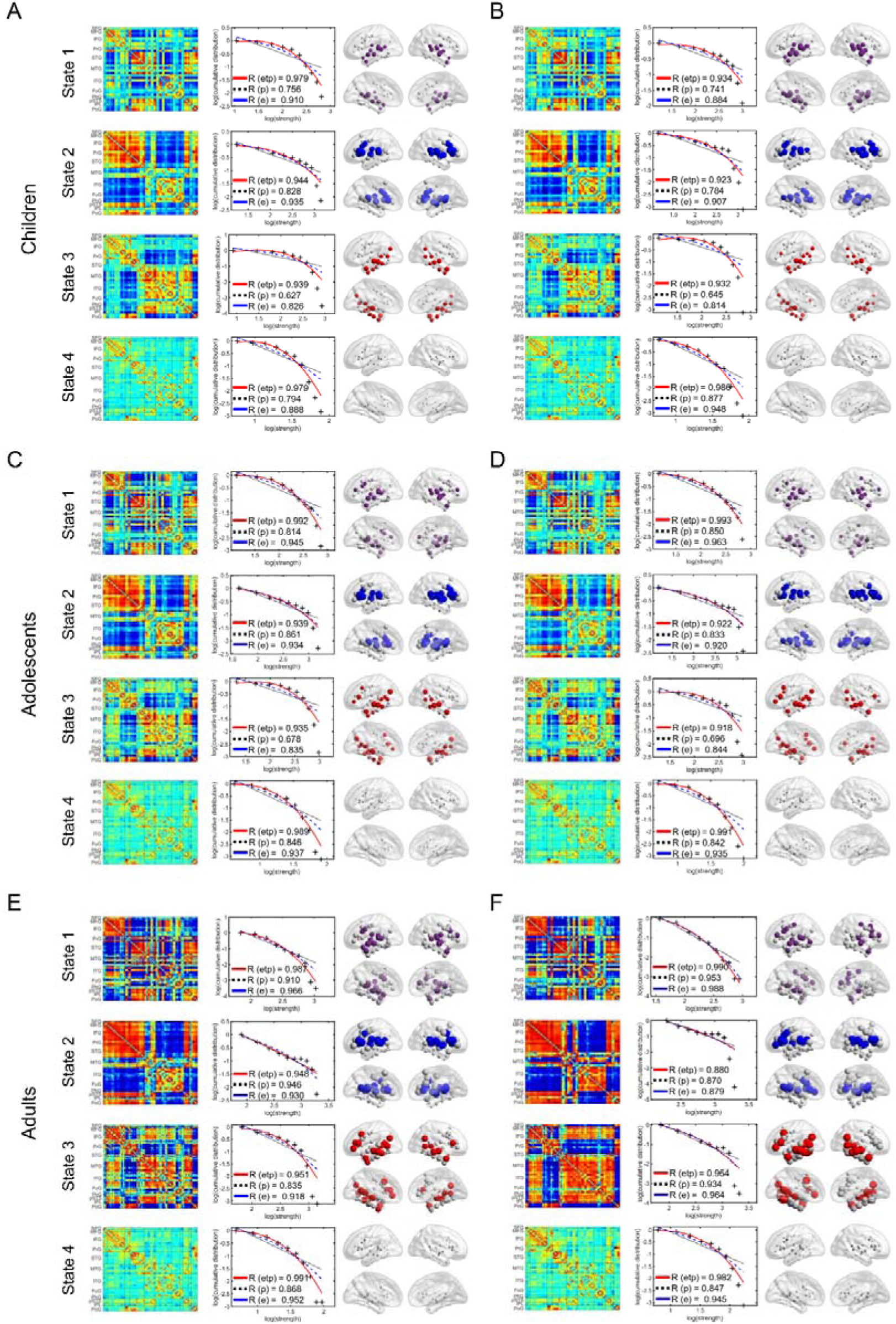
Cortical language network dynamics in HCs and ASDs. The connectivity matrices of the four states in ASDs (A, C, and E) and HCs (B, D, and F), log-log plots of the cumulative nodal strength distributions, and nodal strength distributions were presented. The plus sign (black) represents observed data, the solid line (red) is the fit of the exponentially truncated power-law distribution, 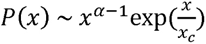, the dashed line (blue) is an exponential distribution, 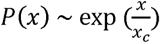, and the dotted line (black) is a power-law, *P(x) ∼ x^α-^*^1^. *R*^2^ was calculated to assess the goodness of fit, with higher values indicating better model fitting. The exponentially truncated power-law is the best fit for all four states. In the third column, the first 20 nodes with the highest nodal strength were defined as hubs.

### All states exhibited rich-club organizations

The normalized rich-club curves were calculated respectively on the weighted network thresholded with a correlation coefficient of 0.2. The normalized rich-club coefficients *φ_norm_(k)* were larger than 1 over a range of values of *K* for all states (Supplementary Fig. 1).

### State-dependent hub distributions followed a domain-segregation pattern

We found that the nodal strength distributions across all states were best fitted by the exponentially truncated power-law form (Fig. 1), indicating the presence of a small set of highly connected hub nodes. In each state, the top 20 nodes with the highest nodal strength were identified as hubs. In the first three states, non-random hub distributions were observed (Fig. 1). In state 1, hubs were mainly located in the STG. In state 2, hubs were primarily in the prefrontal cortex and posterior temporal cortex. In state 3, hubs were primarily in the temporal cortex, posterior parietal cortex, and IFG. We did not consider the hubs in state 4 due to the weak connectivity strength.

To evaluate the functional importance of hubs in the first three states, we conducted term-based meta-analyses using NeuroQuery. Based on the hub distributions, five terms, speech perception, inner speech, phonological processing, speech production, and semantic processing, were analyzed. The dice coefficients between hubs and meta-results varied by term and state (Supplementary Fig. 2). For example, in healthy adolescents, the dice coefficients were: 1) state 1: speech perception, 0.44; inner speech, 0.47; phonological processing, 0.46; speech production, 0.44; semantic, 0.22. State 2: speech perception, 0.36; inner speech, 0.38; phonological processing, 0.46; speech production, 0.37; semantic, 0.21. State 3: speech perception, 0.45; inner speech, 0.47; phonological processing, 0.53; speech production, 0.44; semantic, 0.26.

Given the critical role of STG in speech perception and phonological processing, we theorized that hubs in state 1 were mainly involved in speech perception and phonological processing (central stream). Hubs in state 2 showed significant overlap with regions linked to speech articulation and phonological processing. Considering the importance of IFG and the ventral part of PrG in speech production(Lu et al., 2021), we theorized that hubs in state 2 were primarily engaged in speech production (dorsal stream). Hubs in state 3 showed substantial overlap with regions associated with all five terms. Given the vital role of IPL, ATL, and orbitofrontal cortex in semantic processing(G. Zhang et al., 2023), we hypothesized that hubs in state 3 were primarily involved in semantic processing (Ventral stream). Similar patterns were observed in the other groups, indicating a domain-segregation of language network dynamics in resting state across all three groups.

### Domain-specific gene expression modules of language networks

Gene-based modular clustering identified three modules (Fig. 2): module 1 was mainly located in the frontal lobe and precentral gyrus; module 2 in the anterior temporal lobe; and module 3 in the posterior temporal lobe, postcentral gyrus, and nearby regions. We then calculated Dice coefficients between the first three meta-states and the gene modules to measure the spatial correspondence between gene expression patterns and network states. Gene module 2 showed the highest overlap with State 2 (Dice = 0.49), gene module 3 had the most overlap with State 1 (Dice = 0.40), and gene module 1 had moderate overlaps with State 1 (Dice = 0.25) and State 3 (Dice = 0.27). These results suggest that different gene expression modules underlie the network organization of language meta-states, offering a genetic basis for the domain specificity of language processing.

**Figure 2.**
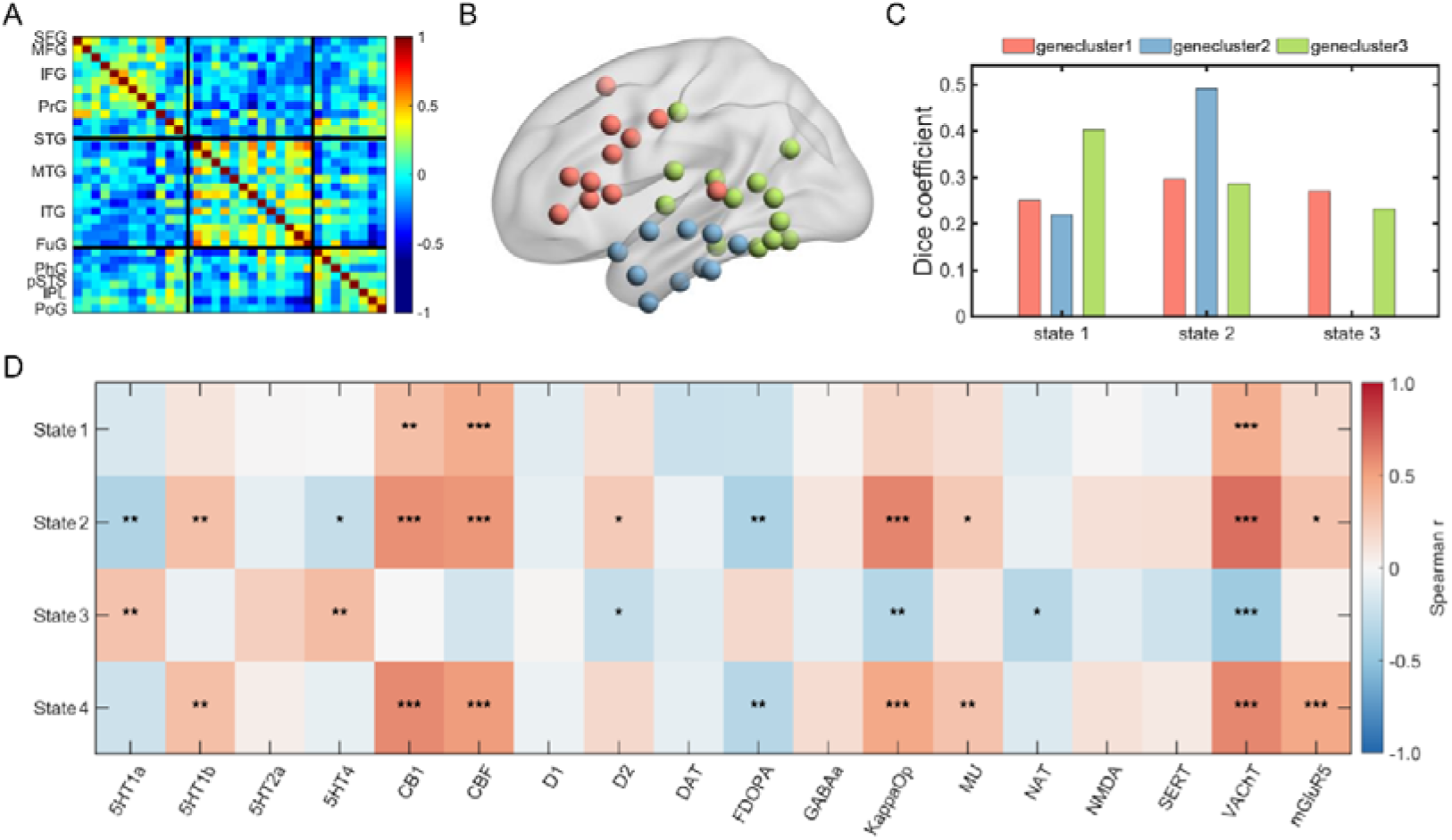
Molecular architecture underlying the language network dynamics. (A) Gene co-expression matrix among left-hemisphere ROIs. Three distinct modules were identified via modularity analysis. (B) Spatial distribution of the modules mapped onto the cortical surface. (C) Dice coefficients quantifying spatial overlap between gene modules and dynamic network states. (D) Spatial correlations between each meta-state’s nodal strength and neurotransmitter distribution maps.

### Neurochemistry architecture related to domain-specific language network patterns

We assessed Spearman spatial correlations between each meta-state’s nodal strength and 18 neurotransmitter distribution maps (Fig. 2). In StateL1, nodal strength correlated positively with CB1, CBF, and VAChT. In StateL2, nodal strength showed strong positive correlations with CB1, CBF, KappaOp, and VAChT; moderate positive correlations with 5-HT1B, D2, MU, and mGluR5; and negative correlations with 5-HT1A, 5-HT4, and FDOPA. In StateL3, nodal strength correlated positively with 5-HT1A and 5-HT4, and negatively with D2, KappaOp, NAT, and VAChT. In StateL4, nodal strength correlated positively with 5-HT1B, CB1, CBF, KappaOp, MU, VAChT, and mGluR5, and negatively with FDOPA. Overall, these patterns indicate that the language network’s neurochemical coupling varies across meta-states, reflecting its dynamic reliance on diverse neurotransmitter systems.

### The hypo- and hyper-connectivity of language network in individuals with ASD

Language network disruptions in individuals with ASD were found in the first three states (Fig. 3). The hypo- and hyper-connectivity of language network in ASD were state-specific and age-dependent. Inter-hemispheric hypo- and hyper-connectivity were dominant. The brain regions exhibiting hypo- and hyper-connectivity include not only the hub regions in each state but also the connections between hub regions and non-hub regions in each state.

**Figure 3.**
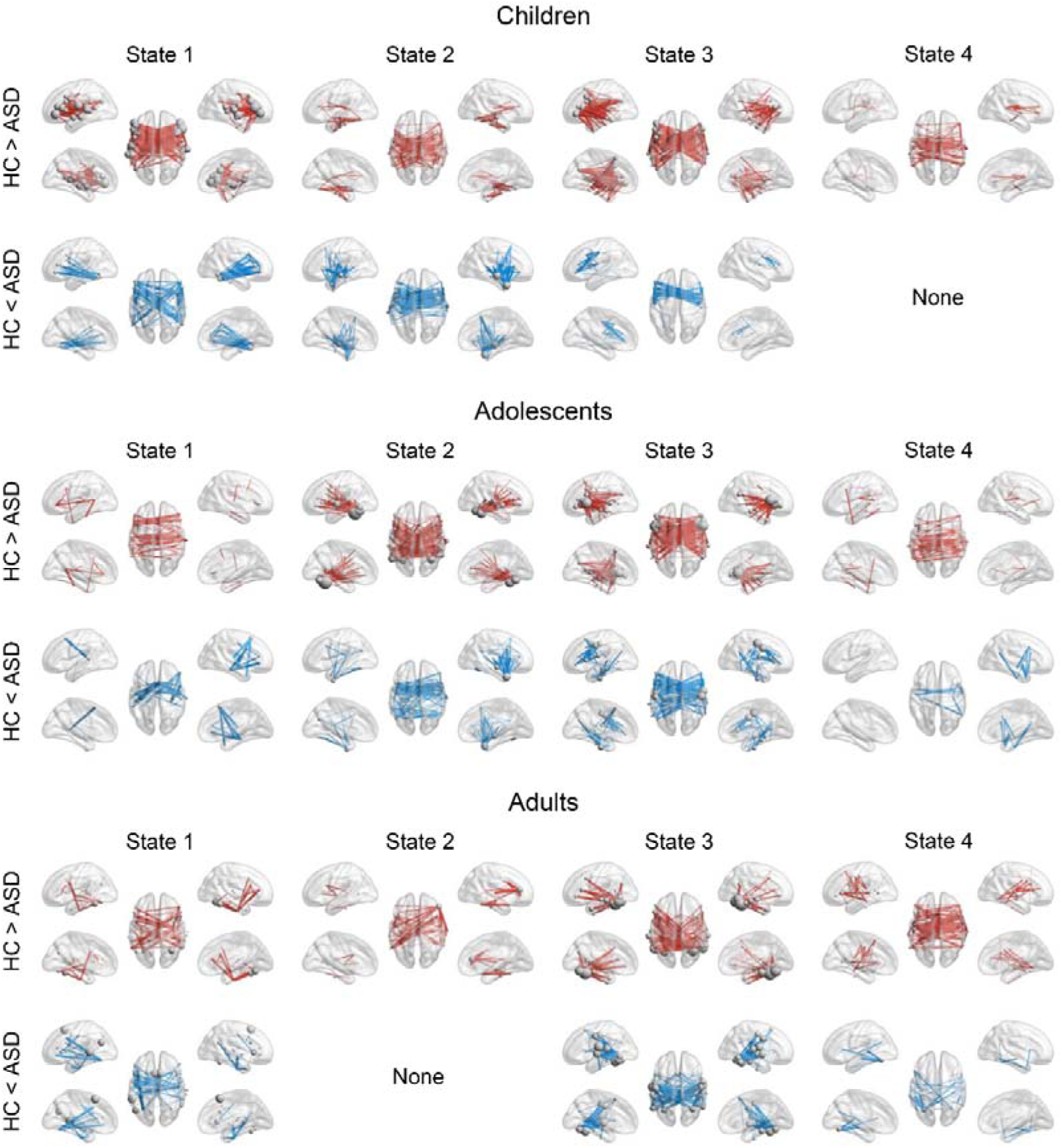
Domain-specific and age-dependent hyper- and hypo-connectivity in individuals with ASD. For each state, differences in edges between groups were corrected using NBS with an edge *P* value of 0.01 and a component *P* value of 0.05.

Those hypo- and hyper-connectivities were significantly correlated with VIQ, ADOS communication, and social scores (Supplementary Figs 3, 4, 5). These significantly correlated brain regions are spread across the frontal, temporal, and parietal lobes. For VIQ, hypo-connectivity shows a negative correlation, while hyper-connectivity shows a positive correlation. Conversely, for ADOS communication and social scores, hypo-connectivity and hyper-connectivity are positively and negatively correlated, respectively. Additionally, we found that hypo- and hyper-connectivity in specific brain regions are significantly linked to ADOS stereotypical scores (Supplementary Fig 6). The regions and the directions of these correlations are also related to age.

**Figure 4.**
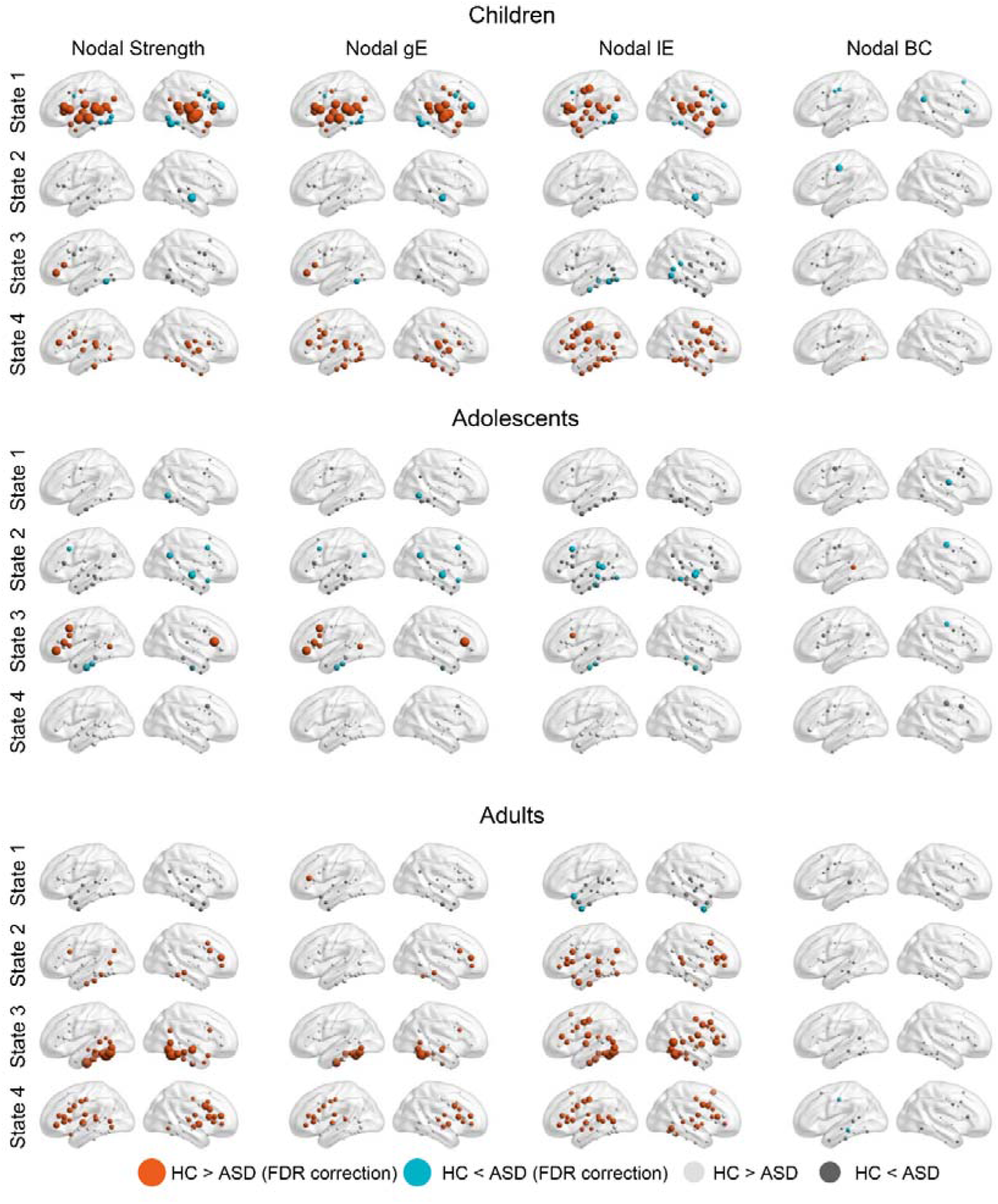
Changes in nodal topological characteristics. Four nodal topological properties—nodal strength, global efficiency (gE), local efficiency (lE), and betweenness centrality (BC)—were calculated for each subject’s median of state dFCs. Between-group statistical comparisons were conducted using two-sample t-tests with FDR correction (*p* < 0.05).

### The global and nodal topological characteristics of language network dynamics in individuals with ASD

The global and nodal topological characteristics are both state- and age-dependent (Figs. 5 and 6). In children, State 1 and State 4 exhibit robust differences for all three metrics, State 3 shows a difference for lE only. Similarly, case-control contrasts in adults are distinct and widespread. Specifically, lE differs between groups in State 1, while State 2 shows broad effects spanning Str, gE, and lE; State 3 also reveals widespread, highly significant differences across all metrics; and State 4 presents consistent differences for Str, gE, and lE. However, in adolescents with ASDs, we observed a focused pattern: significant differences concentrate primarily in State 2, with no reliable effects in States 1, 3, or 4. Overall, case-control contrasts are minimal in adolescence but become widespread in childhood and adulthood, and those topological characteristics were significantly correlated with VIQ, ADOS communication, social and stereotypical scores (Supplementary Tables 5, 6, 7, 8).

**Figure 5.**
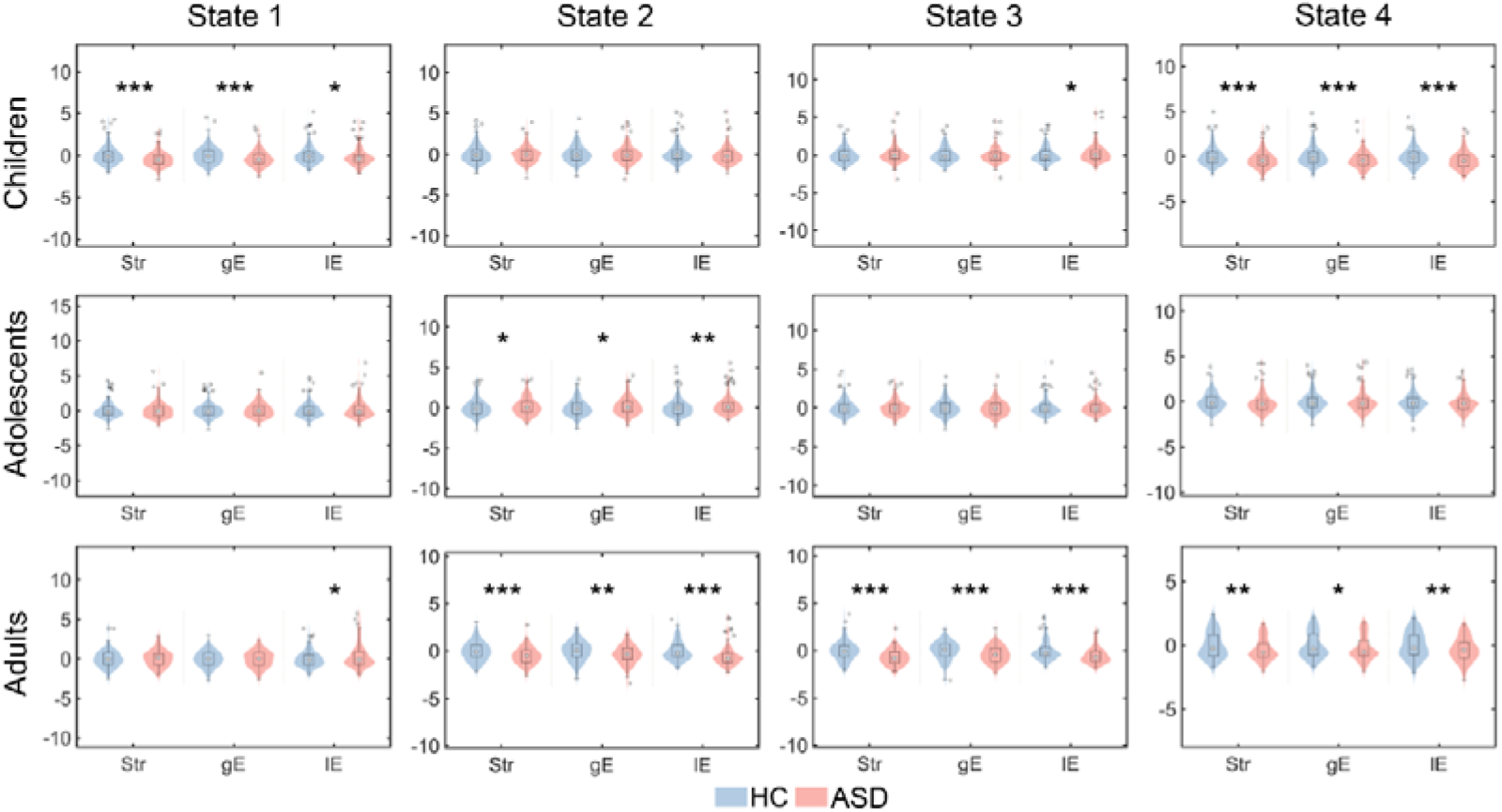
Global topological characteristics of each state in individuals with ASD. Three network topological properties—total functional connectivity strength (Str), network global efficiency (gE), and local efficiencies (lE)—were calculated using each subject’s median of state dFCs. For clarity, the topological values were z-scored. The “*” symbol indicates statistical comparisons between HC and ASD with an uncorrected p-value of 0.05. *, uncorrected *p* < 0.05; **, uncorrected *p* < 0.01; ***, uncorrected *p* < 0.001.

**Figure 6.**
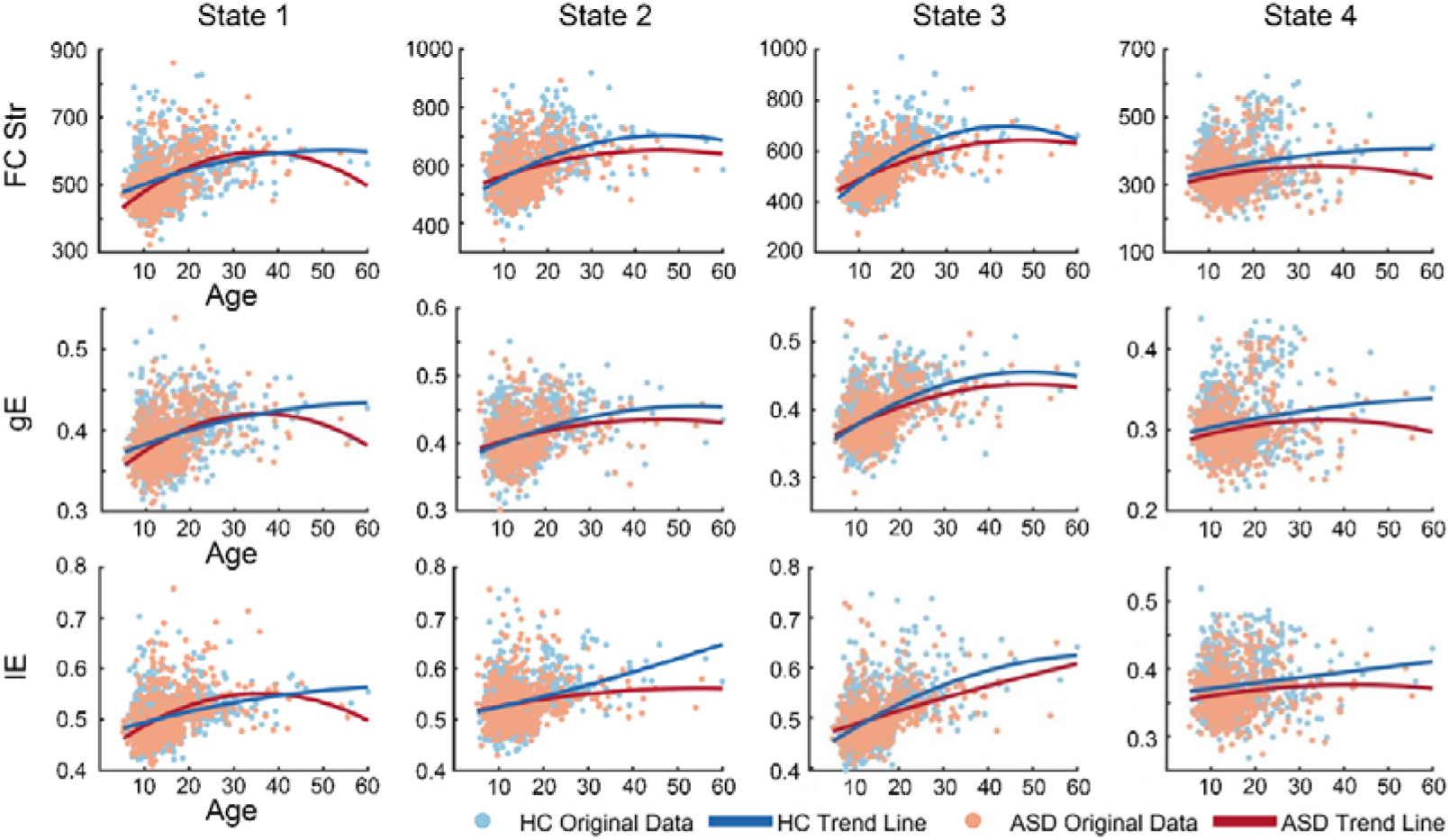
Domain-specific development trajectories of language networks fitted using polynomial functions. Graphs show the original data (light blue for HC, light red for ASD), with the fitted trajectories superimposed (solid blue for HC, light red for ASD).

### Domain-specific development trajectories of language network in ASD

The developmental trajectories of the three topological properties in State 1 show that the ASD trajectory increases through childhood and adolescence, briefly exceeding the HC trajectory during the teenage years (around 15-18 years), then plateaus and declines in adulthood, culminating in a second crossover at approximately age 40 after which HC exceeds ASD. In States 2 and 3, ASD lies slightly above HC in early adolescence but is subsequently overtaken by HC, which maintains the advantage from mid-adolescence into adulthood. In State 4, HC remains higher across all metrics, with gradually widening gaps into adulthood and no intersections, reflecting a modest increase in HC contrasted with an ASD trajectory that plateaus or declines at later ages. Overall, HC exhibits more sustained age-related gains, whereas ASD shows earlier peaks followed by plateauing or decline. (Fig. 6; Supplementary Fig. 7).

### The dFCs significantly predicted VIQ and communication scores in individuals with ASD

RVR-based dFC-deficits prediction analyses showed that dFCs of the four states significantly predicted VIQ in individuals with ASD across all three groups (predicted vs. actual, permutation tests, children, *r* = 0.333, *p* = 0.019; adolescents, *r* = 0.307, *p* = 0.005; adults, *r* = 0.348, *p* = 0.036) (Figure 7). The dFCs of the four states also significantly predicted ADOS communication scores in adolescents (*r* = 0.333, *p* = 0.006) and adults with ASD (*r* = 0.377, *p* = 0.013).

**Figure 7.**
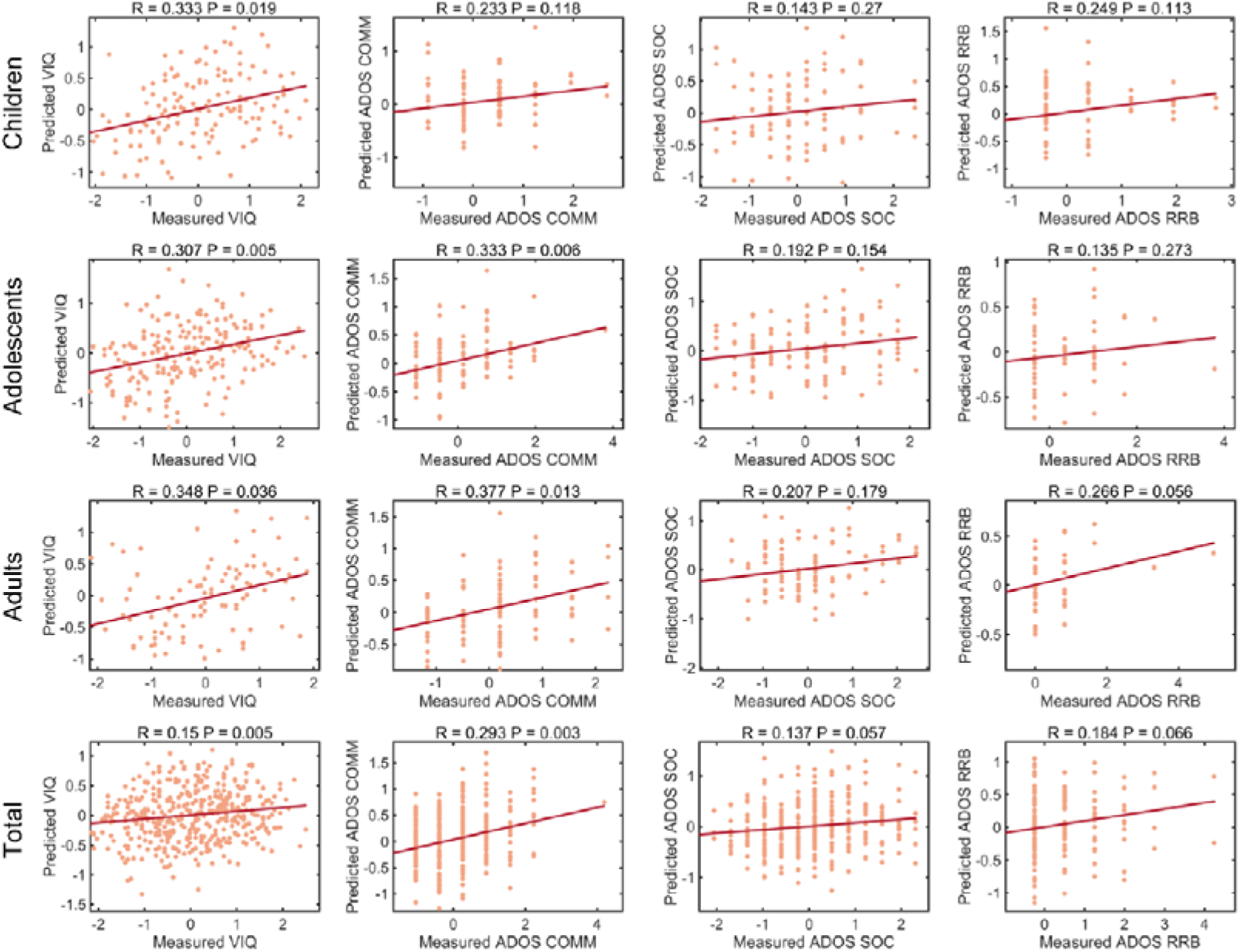
dFC-deficits model accuracies and significance. The scatter plots displayed both actual (normalized within each group) and predicted scores, along with the corresponding linear fit lines. The *R* value represents the Pearson correlation coefficient between the predicted and actual scores, while the model significance and *P* values were calculated based on 1000 permutation tests.

We then constructed prediction models for all ASD samples and found that dFCs of the four states still significantly predicted verbal IQ (*r=* 0.15, *p* = 0.005) and ADOS communication scores (*r* = 0.293, *p* = 0.003).

### Between-group comparisons of sFC

For sFC, inter-hemisphere hypo-connectivity was observed in all three ASD groups (Supplementary Fig. 8). Hyper-connectivity was only seen in children with ASD. Several instances of hypo- and hyper-connectivity correlated with VIQ scores (Supplementary Fig. 9). There was no significant group difference in either nodal or global topological properties (Supplementary Figs. 10 and 11).

Additionally, sFC-deficits prediction analyses showed that sFC significantly predicted ASD’s verbal IQ (children, *r* = 0.46, *p* = 0.002; adolescents, *r* = 0.35, *p* = 0.009; adults, *r* = 0.33, *p* = 0.04; total, *r* = 0.34, *p* < 0.001) and ADOS communication scores (adolescents, *r* = 0.32, *p* = 0.018; adults, *r* = 0.45, *p* = 0.001; total, *r* = 0.27, *p* = 0.046) (Supplementary Fig. 12).

### Validation analysis

Consistent with the ABIDE cohort, we observed comparable language network connectivity patterns in the validation cohort (Supplementary Fig. 13). Spatial similarity analyses showed that state similarity between the two cohorts ranged from 0.48 to 0.97 (Supplementary Fig. 14). Furthermore, the dynamic patterns of the language network were replicated in the NYU dataset (Supplementary Figs. 15 and 16). In HC, similarity between the NYU and ABIDE cohorts ranged from 0.73 to 0.96 (Supplementary Fig. 17), whereas in the ASD group it ranged from 0.75 to 0.96 (Supplementary Fig. 17). In the validation cohort, the age-related trajectories observed in the HC group closely resembled those in the ABIDE cohort (Supplementary Fig. 18). By combining the four dynamic connectivity patterns of the language network with developmental trajectories, we found that the four language network dynamic patterns consistent with the full sample could already be reliably identified during childhood, indicating that the domain-segregation pattern of the language network is largely established by approximately five years of age.

## Discussion

In this study, we analyzed resting-state fMRI data from the ABIDE datasets to examine the aberrant dynamics of cortical language network in children, adolescents, and adults with ASD. Unlike previous studies that focused solely on network connectivity, we employed the dynamic “meta-networking” model of language during resting state, as proposed by Yuan et al.(Yuan, Xie, Wang, et al., 2023). This framework enabled us to scrutinize domain-specific alterations in individuals with ASD. We demonstrated that: (1) the language network dynamics in healthy children, adolescents, and adults exhibited domain-segregation patterns. The observation of domain-separation language network dynamics in healthy children suggests that the functional segregation and specialization of domain-related language processing mature during preschool-age(Brauer et al., 2013; Ernst et al., 2015; Olulade et al., 2020; O’Muircheartaigh et al., 2013; Reynolds et al., 2019); (2) the domain-segregation dynamics have a genetic and neurochemical foundation. Different gene expression modules and neurochemical coupling underlie the network organization of language meta-states; (3) hypo- and hyper-connectivity in the language network in ASD encompasses not only hub regions in each state but also hub–non hub connections within each state. Changes in global and nodal graph-theoretic characteristics indicated that children exhibited broad alterations in States 1 (central stream) and 4 (baseline), with additional effects on local efficiency in State 3 (ventral stream). Adolescents showed a more focal effect confined to State 2 (dorsal stream). Adults displayed widespread alterations across States 2 - 4, with local efficiency also affected in State 1; (4) connections across the four states significantly predicted individuals’ verbal IQ and communication scores based on the machine learning-based models. Together, this study revealed domain-specific and age-dependent language-network disruptions in ASD. These results advance a network-level model of language dysfunction in ASD and highlight potential mechanistic pathways for targeted interventions.

A recent literature review noted that individuals with ASD exhibit under-connectivity of inter-hemispheric language regions(Larson et al., 2025). However, we found that the language network changes in individuals with ASDs were complex. We found that the domain-segregation language network dynamics of individuals with ASD were sub-optimal, and domain- and state-specific hyper- and hypo-connectivity were observed for various critical language areas. Specifically, in children with ASD, we observed hypo- and hyper-connectivity among IFG, PrG, PoG and temporal cortex in states 2 and 3. In adolescents with ASD, hypo- and hyper-connectivity among IFG, STG, and MTG were observed in states 1 and 2. Similarly, in adults with ASD, hypo-and hyper-connectivity among IFG, STG, and ITG were observed in state 1. These language network changes were also associated with and predictable for VIQ and ADOS communication scores, but not for social and stereotyped behavior scores, suggesting their language specificity(Anderson et al., 2007; Bedford et al., 2025; Buch, 2023; Lombardo et al., 2018; Pickles et al., 2014; Schaeffer et al., 2023; Seol et al., 2014; Xiao et al., 2022).

The most notable feature of brain development in individuals with ASD is overgrowth from infancy to early adolescence, followed by undergrowth from late adolescence into adulthood(Courchesne, 2003; Courchesne et al., 2011; Hardan et al., 2006; Lee et al., 2021; Schumann et al., 2010; The IBIS Network et al., 2017). A more detailed analysis of brain regions and networks showed that the developmental paths of network properties vary across different brain regions(Kozhemiako et al., 2019) and networks(Ball et al., 2017). In our cross-sectional samples, we observed that the developmental paths of the topological properties in the first three states in the ASD group match the aforementioned pattern compared to the control group. Interestingly, however, the crossover points between the developmental paths of state 2 and state 3 with state 1 differed between the ASD and control groups. In State 1 the trajectories cross twice, first in the teenage years when ASD briefly exceeds HC and again around 40 years when HC becomes higher, while in States 2 and 3 a single crossover occurs in mid adolescence, approximately 15 years. The hub regions of state 1 are mainly located in the STG and the ventral part of the sensorimotor network. These regions are involved in speech sound perception and representation, serving as the key interface between auditory and language systems(Bhaya-Grossman & Chang, 2022). A significant body of research has documented overgrowth of the bilateral STG in individuals with ASD(Bedford et al., 2025; Ecker et al., 2022; Jou et al., 2010; Khundrakpam et al., 2017; Wang et al., 2022), with this anomaly persisting from infancy(Courchesne, 2003; Duan et al., 2024; Lombardo et al., 2015) into young adulthood. Hypoactivation of the STG has been shown to be a neural marker of poorer language outcomes later on in individuals with ASD(Duan et al., 2024; Lombardo et al., 2015; Xiao et al., 2022). Besides the crucial role of the STG in language development, some studies suggest that disruptions in functional connectivity in the STG and superior temporal sulcus are also linked to emotional disturbances in individuals with ASD(Alaerts et al., 2014). These findings suggest that ongoing anomalies in speech perception and representation could underlie the atypical language development associated with ASD, as well as its more complex emotional and social dysfunctions.

Furthermore, by integrating gene expression and neurotransmitter profiles, we found that each meta-state is characterized by a distinct neuromodulatory signature. StateL1 showed strong positive correlations with CB1, CBF, and VAChT. Notably, CBF influences speech perception and processing(Paus et al., 1996; J. Zhang et al., 2022), and CB1 indirectly influences sensory processing and cognitive activity(De La Salle et al., 2019; Hajós et al., 2008; Smith & Zheng, 2016; Whitney et al., 2003), implying that cholinergic and endocannabinoid systems support early auditory and phonological processes. StateL2 involves multiple neurochemical systems—including endocannabinoid, cholinergic, opioid, serotonergic, dopaminergic, glutamatergic systems, and CBF. Glutamate, a primary excitatory neurotransmitter, is vital for normal language processing and the understanding language disorders(W. Li et al., 2020). Newer evidence emphasizes a close interaction between glutamatergic and dopaminergic systems, where increased subcortical dopamine can disrupt cortical glutamate balance(Jauhar et al., 2018). Opioid receptors play a critical role in initiating and executing behaviors essential for speech production. StateL3 was marked by positive correlations with 5-HT1a and 5-HT4, and negative correlations with D2, KappaOp, NAT, and VAChT. This neurochemical profile is supported by evidence that dopaminergic signaling via D2 receptors enhances semantic activation by speeding up access to lexical and conceptual representations(Andreou et al., 2014), and that the cholinergic system is crucial for semantic fluency and quick word retrieval(Kehagia et al., 2013).

Taken together with the observed spatial associations between dynamic states and gene expression modules, these findings provide converging molecular evidence supporting the validity of the meta-networking model of language. They further imply that the functional specificity and segregation of language-related brain states rely on shared gene expression patterns and coordinated neuromodulatory systems. Moreover, the presence of such spatial correspondences alongside the suboptimal dynamic network patterns identified in individuals with ASD suggests that abnormalities may also exist at the genetic and neurotransmitter levels(Buch, 2023; Shan et al., 2025). Further studies are needed to substantiate and refine this interpretation.

## Limitations

This study has several limitations. First, this study did not include structured linguistic assessments for ASD, such as those related to phonology, semantics, and syntax, nor did it cover lexicon and pragmatics. Although we identified state-specific differences, we were unable to explore further whether these differences are linked to domain-specific deficits. In fact, both VIQ and ADOS assessments are highly complex, covering various aspects of both linguistic and non-linguistic abilities. This complexity may explain why the functional connectivity observed in this study, as well as in other related studies, shows only weak correlations with these measures. Second, while our research contributes to a theoretical framework of abnormal language network dynamics, the clinical relevance regarding diagnosis and intervention remains unclear. Based on our findings, a reasonable hypothesis is that the dynamic “meta-networking” framework of language could potentially serve as a reference to evaluate the degree of abnormality in the language network of individuals with ASD or the extent of reorganization after intervention. Third, due to the age range limitations of the ABIDE dataset (participants aged five and above), we were unable to examine language network abnormalities before the age of five. The period before the age of five is critical for language development. Understanding the dynamic interactions and separation between language-related brain areas, particularly in the frontal and temporal lobes during this time, is not only a significant scientific question but also essential for understanding language development abnormalities in ASD.

## Conclusions

In this study, we refined the characterization of hypo- and hyper-connectivity patterns within the language network of children, adolescents, and adults with ASD by applying the dynamic “meta-networking” framework of language. We further demonstrated that domain-segregation cortical language dynamics are biologically grounded, showing spatial correspondences with gene expression modules and neuromodulatory systems. Together, these findings advance a network-level model of language dysfunction in ASD, providing mechanistic insights into how atypical developmental trajectories of language networks may give rise to verbal and communicative impairments.

## Data/code availability statement

Behavioral and fMRI data are available at https://fcon_1000.projects.nitrc.org/indi/abide/. The computation of DCC and the visualization of results were performed using th e NaDyNet toolbox(Yang et al., 2025), available at [https://github.com/yuanbinke/Naturalistic-Dynamic-Network-Toolbox]. Other software used in the study was based on DPABI (http://rfmri.org/dpabi), SPM (https://www.fil.ion.ucl.ac.uk/spm/), GRETNA (https://www.nitrc.org/projects/gretna/), and BrainNetViewer (https://www.nitrc.org/projects/bnv/).

## Contributions

Zhe Hu contributed to fMRI data analysis, results visualization and interpretation, the revision of the manuscript, and the quality control of phenotypic data.

Xiaolin Guo contributed to fMRI data analysis, interpretation of results, and the quality control of phenotypic and imaging data.

Junjie Yang contributed to data analysis, computer programming.

Zhiheng Qu and Zhongqi Li contributed to the quality control of phenotypic and imaging data.

Junjing Li contributed to fMRI data analysis.

Xiaowei Gao, Jiaxuan Liu, Yaling Wang, Wanchun Li, Wanjing Li, Yien Huang and Jiali Chen contributed to the interpretation of results.

Xin Wang gained funding, interpreted the results, critically reviewed, and revised the manuscript.

Hui Xie gained funding, interpreted the results, critically reviewed, and revised the manuscript.

Binke Yuan conceptualized the study design, secured funding, supervised data analysis, interpreted the results, wrote the draft, critically reviewed, and revised the manuscript.

All authors reviewed the manuscript, participated in revision, and approved the final version.

## References

Alaerts, K., Woolley, D. G., Steyaert, J., Di Martino, A., Swinnen, S. P., & Wenderoth, N. (2014). Underconnectivity of the superior temporal sulcus predicts emotion recognition deficits in autism. Social Cognitive and Affective Neuroscience, 9(10), 1589–1600. 10.1093/scan/nst156

Allen, E. A., Damaraju, E., Plis, S. M., Erhardt, E. B., Eichele, T., & Calhoun, V. D. (2014). Tracking Whole-Brain Connectivity Dynamics in the Resting State. Cerebral Cortex, 24(3), 663–676. 10.1093/cercor/bhs352

Anderson, D. K., Lord, C., Risi, S., DiLavore, P. S., Shulman, C., Thurm, A., Welch, K., & Pickles, A. (2007). Patterns of growth in verbal abilities among children with autism spectrum disorder. Journal of Consulting and Clinical Psychology, 75(4), 594–604. 10.1037/0022-006X.75.4.594

Andreou, C., Veith, K., Bozikas, V., Lincoln, T., & Moritz, S. (2014). Effects of dopaminergic modulation on automatic semantic priming: A double-blind study. Journal of Psychiatry & Neuroscience, 39(2). 10.1503/jpn.130035

Andrés-Roqueta, C., & Katsos, N. (2017). The Contribution of Grammar, Vocabulary and Theory of Mind in Pragmatic Language Competence in Children with Autistic Spectrum Disorders. Frontiers in Psychology, 8, 996. 10.3389/fpsyg.2017.00996

Arnatkevic□iūtė, A., Fulcher, B. D., & Fornito, A. (2019). A practical guide to linking brain-wide gene expression and neuroimaging data. NeuroImage, 189, 353–367. 10.1016/j.neuroimage.2019.01.011

Asghari, S. Z., Farashi, S., Bashirian, S., & Jenabi, E. (2021). Distinctive prosodic features of people with autism spectrum disorder: A systematic review and meta-analysis study. Scientific Reports, 11(1), 23093. 10.1038/s41598-021-02487-6

Bal, V. H., Fok, M., Lord, C., Smith, I. M., Mirenda, P., Szatmari, P., Vaillancourt, T., Volden, J., Waddell, C., Zwaigenbaum, L., Bennett, T., Duku, E., Elsabbagh, M., Georgiades, S., Ungar, W. J., & Zaidman□Zait, A. (2020). Predictors of longer□term development of expressive language in two independent longitudinal cohorts of language□delayed preschoolers with Autism Spectrum Disorder. Journal of Child Psychology and Psychiatry, 61(7), 826–835. 10.1111/jcpp.13117

Ball, G., Beare, R., & Seal, M. L. (2017). Network component analysis reveals developmental trajectories of structural connectivity and specific alterations in autism spectrum disorder. Human Brain Mapping, 38(8), 4169–4184. 10.1002/hbm.23656

Bedford, S. A., Lai, M.-C., Lombardo, M. V., Chakrabarti, B., Ruigrok, A., Suckling, J., Anagnostou, E., Lerch, J. P., Taylor, M., Nicolson, R., Stelios, G., Crosbie, J., Schachar, R., Kelley, E., Jones, J., Arnold, P. D., Courchesne, E., Pierce, K., Eyler, L. T., … Williams, S. C. (2025). Brain-Charting Autism and Attention-Deficit/Hyperactivity Disorder Reveals Distinct and Overlapping Neurobiology. Biological Psychiatry, 97(5), 517–530. 10.1016/j.biopsych.2024.07.024

Betzel, R. F., & Bassett, D. S. (2017). Multi-scale brain networks. NeuroImage, 160, 73–83. 10.1016/j.neuroimage.2016.11.006

Bhaya-Grossman, I., & Chang, E. F. (2022). Speech Computations of the Human Superior Temporal Gyrus. Annual Review of Psychology, 73(1), 79–102. 10.1146/annurev-psych-022321-035256

Blume, J., Wittke, K., Naigles, L., & Mastergeorge, A. M. (2021). Language Growth in Young Children with Autism: Interactions Between Language Production and Social Communication. Journal of Autism and Developmental Disorders, 51(2), 644–665. 10.1007/s10803-020-04576-3

Brauer, J., Anwander, A., Perani, D., & Friederici, A. D. (2013). Dorsal and ventral pathways in language development. Brain and Language, 127(2), 289–295. 10.1016/j.bandl.2013.03.001

Brignell, A., Morgan, A. T., Woolfenden, S., Klopper, F., May, T., Sarkozy, V., & Williams, K. (2018). A systematic review and meta-analysis of the prognosis of language outcomes for individuals with autism spectrum disorder. Autism & Developmental Language Impairments, 3, 2396941518767610. 10.1177/2396941518767610

Brignell, A., Williams, K., Jachno, K., Prior, M., Reilly, S., & Morgan, A. T. (2018). Patterns and Predictors of Language Development from 4 to 7 Years in Verbal Children With and Without Autism Spectrum Disorder. Journal of Autism and Developmental Disorders, 48(10), 3282–3295. 10.1007/s10803-018-3565-2

Buch, A. M. (2023). Molecular and network-level mechanisms explaining individual differences in autism spectrum disorder. Nature Neuroscience, 26, 650–663.

Cermak, C. A., Arshinoff, S., Ribeiro De Oliveira, L., Tendera, A., Beal, D. S., Brian, J., Anagnostou, E., & Sanjeevan, T. (2022). Brain and Language Associations in Autism Spectrum Disorder: A Scoping Review. Journal of Autism and Developmental Disorders, 52(2), 725–737. 10.1007/s10803-021-04975-0

Chai, L. R., Mattar, M. G., Blank, I. A., Fedorenko, E., & Bassett, D. S. (2016). Functional Network Dynamics of the Language System. Cerebral Cortex, 26(11), 4148–4159. 10.1093/cercor/bhw238

Chan, C. Y. Z., Williams, K., May, T., Wan, W. H., & Brignell, A. (2023). Is language ability associated with behaviors of concern in autism? A systematic review. Autism Research, 16(2), 250–270. 10.1002/aur.2855

Chen, H., Nomi, J. S., Uddin, L. Q., Duan, X., & Chen, H. (2017). Intrinsic functional connectivity variance and state□specific under□connectivity in autism. Human Brain Mapping, 38(11), 5740–5755. 10.1002/hbm.23764

Chen, Y., Siles, B., & Tager□Flusberg, H. (2024). Receptive language and receptive□expressive discrepancy in minimally verbal autistic children and adolescents. Autism Research, 17(2), 381–394. 10.1002/aur.3079

Choe, A. S., Nebel, M. B., Barber, A. D., Cohen, J. R., Xu, Y., Pekar, J. J., Caffo, B., & Lindquist, M. A. (2017). Comparing test-retest reliability of dynamic functional connectivity methods. NeuroImage, 158, 155–175. 10.1016/j.neuroimage.2017.07.005

Courchesne, E. (2003). Evidence of Brain Overgrowth in the First Year of Life in Autism. JAMA, 290(3), 337. 10.1001/jama.290.3.337

Courchesne, E., Karns, C., Davis, H., Ziccardi, R., Carper, R., Tigue, Z., Chisum, H. J., Moses, P., Pierce, K., Lord, C., Lincoln, A., Pizzo, S., Schreibman, L., Haas, R., Akshoomoff, N., & Courchesne, R. (2011). Unusual brain growth patterns in early life in patients with autistic disorder: An MRI study. Neurology, 76(24), 2111–2111. 10.1212/01.wnl.0000399191.79091.28

Courchesne, E., Pramparo, T., Gazestani, V. H., Lombardo, M. V., Pierce, K., & Lewis, N. E. (2019). The ASD Living Biology: From cell proliferation to clinical phenotype. Molecular Psychiatry, 24(1), 88–107. 10.1038/s41380-018-0056-y

Cui, Z., & Gong, G. (2018). The effect of machine learning regression algorithms and sample size on individualized behavioral prediction with functional connectivity features. NeuroImage, 178, 622–637. 10.1016/j.neuroimage.2018.06.001

Damaraju, E., Allen, E. A., Belger, A., Ford, J. M., McEwen, S., Mathalon, D. H., Mueller, B. A., Pearlson, G. D., Potkin, S. G., Preda, A., Turner, J. A., Vaidya, J. G., Van Erp, T. G., & Calhoun, V. D. (2014). Dynamic functional connectivity analysis reveals transient states of dysconnectivity in schizophrenia. NeuroImage: Clinical, 5, 298–308. 10.1016/j.nicl.2014.07.003

De La Salle, S., Inyang, L., Impey, D., Smith, D., Choueiry, J., Nelson, R., Heera, J., Baddeley, A., Ilivitsky, V., & Knott, V. (2019). Acute separate and combined effects of cannabinoid and nicotinic receptor agonists on MMN-indexed auditory deviance detection in healthy humans. Pharmacology Biochemistry and Behavior, 184, 172739. 10.1016/j.pbb.2019.172739

Di Martino, A., O’Connor, D., Chen, B., Alaerts, K., Anderson, J. S., Assaf, M., Balsters, J. H., Baxter, L., Beggiato, A., Bernaerts, S., Blanken, L. M. E., Bookheimer, S. Y., Braden, B. B., Byrge, L., Castellanos, F. X., Dapretto, M., Delorme, R., Fair, D. A., Fishman, I., … Milham, M. P. (2017). Enhancing studies of the connectome in autism using the autism brain imaging data exchange II. Scientific Data, 4(1), 170010. 10.1038/sdata.2017.10

Di Martino, A., Yan, C.-G., Li, Q., Denio, E., Castellanos, F. X., Alaerts, K., Anderson, J. S., Assaf, M., Bookheimer, S. Y., Dapretto, M., Deen, B., Delmonte, S., Dinstein, I., Ertl-Wagner, B., Fair, D. A., Gallagher, L., Kennedy, D. P., Keown, C. L., Keysers, C., … Milham, M. P. (2014). The autism brain imaging data exchange: Towards a large-scale evaluation of the intrinsic brain architecture in autism. Molecular Psychiatry, 19(6), 659–667. 10.1038/mp.2013.78

Dichter, G. S. (2012). Functional magnetic resonance imaging of autism spectrum disorders. Dialogues in Clinical Neuroscience, 14(3), 319–351. 10.31887/DCNS.2012.14.3/gdichter

Dockès, J., Poldrack, R. A., Primet, R., Gözükan, H., Yarkoni, T., Suchanek, F., Thirion, B., & Varoquaux, G. (2020). NeuroQuery, comprehensive meta-analysis of human brain mapping. eLife, 9, e53385. 10.7554/eLife.53385

Duan, K., Eyler, L., Pierce, K., Lombardo, M. V., Datko, M., Hagler, D. J., Taluja, V., Zahiri, J., Campbell, K., Barnes, C. C., Arias, S., Nalabolu, S., Troxel, J., Ji, P., & Courchesne, E. (2024). Differences in regional brain structure in toddlers with autism are related to future language outcomes. Nature Communications, 15(1), 5075. 10.1038/s41467-024-48952-4

Duffau, H., Moritz-Gasser, S., & Mandonnet, E. (2014). A re-examination of neural basis of language processing: Proposal of a dynamic hodotopical model from data provided by brain stimulation mapping during picture naming. Brain and Language, 131, 1–10. 10.1016/j.bandl.2013.05.011

Duvall, L., May, K. E., Waltz, A., & Kana, R. K. (2023). The neurobiological map of theory of mind and pragmatic communication in autism. Social Neuroscience, 18(4), 191–204. 10.1080/17470919.2023.2242095

Ecker, C., Pretzsch, C. M., Bletsch, A., Mann, C., Schaefer, T., Ambrosino, S., Tillmann, J., Yousaf, A., Chiocchetti, A., Lombardo, M. V., Warrier, V., Bast, N., Moessnang, C., Baumeister, S., Dell’Acqua, F., Floris, D. L., Zabihi, M., Marquand, A., Cliquet, F., … Murphy, D. G. M. (2022). Interindividual Differences in Cortical Thickness and Their Genomic Underpinnings in Autism Spectrum Disorder. American Journal of Psychiatry, 179(3), 242–254. 10.1176/appi.ajp.2021.20050630

Engle, R. F. (1982). Autoregressive Conditional Heteroscedasticity with Estimates of the Variance of United Kingdom Inflation. Econometrica, 50(4), 987. 10.2307/1912773

Ernst, M., Torrisi, S., Balderston, N., Grillon, C., & Hale, E. A. (2015). fMRI Functional Connectivity Applied to Adolescent Neurodevelopment. Annual Review of Clinical Psychology, 11(1), 361–377. 10.1146/annurev-clinpsy-032814-112753

Falahpour, M., Thompson, W. K., Abbott, A. E., Jahedi, A., Mulvey, M. E., Datko, M., Liu, T. T., & Müller, R.-A. (2016). Underconnected, But Not Broken? Dynamic Functional Connectivity MRI Shows Underconnectivity in Autism Is Linked to Increased Intra-Individual Variability Across Time. Brain Connectivity, 6(5), 403–414. 10.1089/brain.2015.0389

Fan, L., Li, H., Zhuo, J., Zhang, Y., Wang, J., Chen, L., Yang, Z., Chu, C., Xie, S., Laird, A. R., Fox, P. T., Eickhoff, S. B., Yu, C., & Jiang, T. (2016). The Human Brainnetome Atlas: A New Brain Atlas Based on Connectional Architecture. Cerebral Cortex, 26(8), 3508–3526. 10.1093/cercor/bhw157

Fan, X.-R., Wang, Y.-S., Chang, D., Yang, N., Rong, M.-J., Zhang, Z., He, Y., Hou, X., Zhou, Q., Gong, Z.-Q., Cao, L.-Z., Dong, H.-M., Nie, J.-J., Chen, L.-Z., Zhang, Q., Zhang, J.-X., Zhang, L., Li, H.-J., Bao, M., … Zuo, X.-N. (2023). A longitudinal resource for population neuroscience of school-age children and adolescents in China. Scientific Data, 10(1), 545. 10.1038/s41597-023-02377-8

Fedorenko, E., Hsieh, P.-J., Nieto-Castañón, A., Whitfield-Gabrieli, S., & Kanwisher, N. (2010). New Method for fMRI Investigations of Language: Defining ROIs Functionally in Individual Subjects. Journal of Neurophysiology, 104(2), 1177–1194. 10.1152/jn.00032.2010

Félix, J., Santos, M. E., & Benitez-Burraco, A. (2024). Specific Language Impairment, Autism Spectrum Disorders and Social (Pragmatic) Communication Disorders: Is There Overlap in Language Deficits? A Review. Review Journal of Autism and Developmental Disorders, 11(1), 86–106. 10.1007/s40489-022-00327-5

Fortin, J.-P., Cullen, N., Sheline, Y. I., Taylor, W. D., Aselcioglu, I., Cook, P. A., Adams, P., Cooper, C., Fava, M., McGrath, P. J., McInnis, M., Phillips, M. L., Trivedi, M. H., Weissman, M. M., & Shinohara, R. T. (2018). Harmonization of cortical thickness measurements across scanners and sites. NeuroImage, 167, 104–120. 10.1016/j.neuroimage.2017.11.024

Fortin, J.-P., Parker, D., Tunç, B., Watanabe, T., Elliott, M. A., Ruparel, K., Roalf, D. R., Satterthwaite, T. D., Gur, R. C., Gur, R. E., Schultz, R. T., Verma, R., & Shinohara, R. T. (2017). Harmonization of multi-site diffusion tensor imaging data. NeuroImage, 161, 149–170. 10.1016/j.neuroimage.2017.08.047

Friedman, L., & Sterling, A. (2019). A Review of Language, Executive Function, and Intervention in Autism Spectrum Disorder. Seminars in Speech and Language, 40(04), 291–304. 10.1055/s-0039-1692964

Fu, Z., Tu, Y., Di, X., Du, Y., Sui, J., Biswal, B. B., Zhang, Z., De Lacy, N., & Calhoun, V. D. (2019). Transient increased thalamic-sensory connectivity and decreased whole-brain dynamism in autism. NeuroImage, 190, 191–204. 10.1016/j.neuroimage.2018.06.003

Gao, Y., Sun, J., Cheng, L., Yang, Q., Li, J., Hao, Z., Zhan, L., Shi, Y., Li, M., Jia, X., & Li, H. (2022). Altered resting state dynamic functional connectivity of amygdala subregions in patients with autism spectrum disorder: A multi-site fMRI study. Journal of Affective Disorders, 312, 69–77. 10.1016/j.jad.2022.06.011

Georgiou, N., & Spanoudis, G. (2021). Developmental Language Disorder and Autism: Commonalities and Differences on Language. Brain Sciences, 11(5), 589. 10.3390/brainsci11050589

Gernsbacher, M. A., Morson, E. M., & Grace, E. J. (2016). Language and Speech in Autism. Annual Review of Linguistics, 2(1), 413–425. 10.1146/annurev-linguistics-030514-124824

Goodwill, A. M., Low, L. T., Fox, P. T., Fox, P. M., Poon, K. K., Bhowmick, S. S., & Chen, S. H. A. (2023). Meta-analytic connectivity modelling of functional magnetic resonance imaging studies in autism spectrum disorders. Brain Imaging and Behavior, 17(2), 257–269. 10.1007/s11682-022-00754-2

Ha, S., Sohn, I.-J., Kim, N., Sim, H. J., & Cheon, K.-A. (2015). Characteristics of Brains in Autism Spectrum Disorder: Structure, Function and Connectivity across the Lifespan. Experimental Neurobiology, 24(4), 273–284. 10.5607/en.2015.24.4.273

Hajós, M., Hoffmann, W. E., & Kocsis, B. (2008). Activation of Cannabinoid-1 Receptors Disrupts Sensory Gating and Neuronal Oscillation: Relevance to Schizophrenia. Biological Psychiatry, 63(11), 1075–1083. 10.1016/j.biopsych.2007.12.005

Hardan, A. Y., Muddasani, S., Vemulapalli, M., Keshavan, M. S., & Minshew, N. J. (2006). An MRI Study of Increased Cortical Thickness in Autism. Am J Psychiatry.

Herbet, G., & Duffau, H. (2020). Revisiting the Functional Anatomy of the Human Brain: Toward a Meta-Networking Theory of Cerebral Functions. Physiological Reviews, 100(3), 1181–1228. 10.1152/physrev.00033.2019

Herringshaw, A. J., Ammons, C. J., DeRamus, T. P., & Kana, R. K. (2016). Hemispheric differences in language processing in autism spectrum disorders: A metaDanalysis of neuroimaging studies. Autism Research, 9(10), 1046–1057. 10.1002/aur.1599

Hickok, G. (2022). The dual stream model of speech and language processing. In Handbook of Clinical Neurology (Vol. 185, pp. 57–69). Elsevier. 10.1016/B978-0-12-823384-9.00003-7

Hickok, G., & Poeppel, D. (2007). The cortical organization of speech processing. Nature Reviews Neuroscience, 8(5), 393–402. 10.1038/nrn2113

Hodgson, V. J., Lambon Ralph, M. A., & Jackson, R. L. (2021). Multiple dimensions underlying the functional organization of the language network. NeuroImage, 241, 118444. 10.1016/j.neuroimage.2021.118444

Hutchison, R. M., Womelsdorf, T., Allen, E. A., Bandettini, P. A., Calhoun, V. D., Corbetta, M., Della Penna, S., Duyn, J. H., Glover, G. H., Gonzalez-Castillo, J., Handwerker, D. A., Keilholz, S., Kiviniemi, V., Leopold, D. A., De Pasquale, F., Sporns, O., Walter, M., & Chang, C. (2013). Dynamic functional connectivity: Promise, issues, and interpretations. NeuroImage, 80, 360–378. 10.1016/j.neuroimage.2013.05.079

Jauhar, S., McCutcheon, R., Borgan, F., Veronese, M., Nour, M., Pepper, F., Rogdaki, M., Stone, J., Egerton, A., Turkheimer, F., McGuire, P., & Howes, O. D. (2018). The relationship between cortical glutamate and striatal dopamine in first-episode psychosis: A cross-sectional multimodal PET and magnetic resonance spectroscopy imaging study. The Lancet Psychiatry, 5(10), 816–823. 10.1016/S2215-0366(18)30268-2

Jou, R. J., Minshew, N. J., Keshavan, M. S., Vitale, M. P., & Hardan, A. Y. (2010). Enlarged right superior temporal gyrus in children and adolescents with autism. Brain Research, 1360, 205–212. 10.1016/j.brainres.2010.09.005

Kalandadze, T., Bambini, V., & Næss, K.-A. B. (2019). A systematic review and meta-analysis of studies on metaphor comprehension in individuals with autism spectrum disorder: Do task properties matter? Applied Psycholinguistics, 40(6), 1421–1454. 10.1017/S0142716419000328

Kalandadze, T., Norbury, C., Nærland, T., & Næss, K.-A. B. (2018). Figurative language comprehension in individuals with autism spectrum disorder: A meta-analytic review. Autism, 22(2), 99–117. 10.1177/1362361316668652

Kehagia, A. A., Barker, R. A., & Robbins, T. W. (2013). Cognitive Impairment in Parkinson’s Disease: The Dual Syndrome Hypothesis. Neurodegenerative Diseases, 11(2), 79–92. 10.1159/000341998

Khundrakpam, B. S., Lewis, J. D., Kostopoulos, P., Carbonell, F., & Evans, A. C. (2017). Cortical Thickness Abnormalities in Autism Spectrum Disorders Through Late Childhood, Adolescence, and Adulthood: A Large-Scale MRI Study. Cerebral Cortex, 27(3), 1721–1731. 10.1093/cercor/bhx038

Kissine, M., Saint-Denis, A., & Mottron, L. (2023). Language acquisition can be truly atypical in autism: Beyond joint attention. Neuroscience & Biobehavioral Reviews, 153, 105384. 10.1016/j.neubiorev.2023.105384

Kozhemiako, N., Vakorin, V., Nunes, A. S., Iarocci, G., Ribary, U., & Doesburg, S. M. (2019). Extreme male developmental trajectories of homotopic brain connectivity in autism. Human Brain Mapping, 40(3), 987–1000. 10.1002/hbm.24427

Kuperberg, G. R., McGuire, P. K., Bullmore, E. T., Brammer, M. J., Rabe-Hesketh, S., Wright, I. C., Lythgoe, D. J., Williams, S. C. R., & David, A. S. (2000). Common and Distinct Neural Substrates for Pragmatic, Semantic, and Syntactic Processing of Spoken Sentences: An fMRI Study. Journal of Cognitive Neuroscience, 12(2), 321–341. 10.1162/089892900562138

Larson, C., Thomas, H. R., Crutcher, J., Stevens, M. C., & Eigsti, I.-M. (2025). Language Networks in Autism Spectrum Disorder: A systematic review of connectivity-based fMRI studies. Review Journal of Autism and Developmental Disorders, 12(1), 110–137. 10.1007/s40489-023-00382-6

Lebo, M. J., & Box-Steffensmeier, J. M. (2008). Dynamic Conditional Correlations in Political Science. American Journal of Political Science, 52(3), 688–704.

Lee, J. K., Andrews, D. S., Ozonoff, S., Solomon, M., Rogers, S., Amaral, D. G., & Nordahl, C. W. (2021). Longitudinal Evaluation of Cerebral Growth Across Childhood in Boys and Girls With Autism Spectrum Disorder. Biological Psychiatry, 90(5), 286–294. 10.1016/j.biopsych.2020.10.014

Lerner, Y., Honey, C. J., Silbert, L. J., & Hasson, U. (2011). Topographic Mapping of a Hierarchy of Temporal Receptive Windows Using a Narrated Story. The Journal of Neuroscience, 31(8), 2906–2915. 10.1523/JNEUROSCI.3684-10.2011

Li, M., Wang, Y., Tachibana, M., Rahman, S., & Kagitani□Shimono, K. (2022). Atypical structural connectivity of language networks in autism spectrum disorder: A meta□analysis of diffusion tensor imaging studies. Autism Research, 15(9), 1585–1602. 10.1002/aur.2789

Li, W., Kutas, M., Gray, J. A., Hagerman, R. H., & Olichney, J. M. (2020). The Role of Glutamate in Language and Language Disorders—Evidence from ERP and Pharmacologic Studies. Neuroscience & Biobehavioral Reviews, 119, 217–241. 10.1016/j.neubiorev.2020.09.023

Li, Y., Zhu, Y., Nguchu, B. A., Wang, Y., Wang, H., Qiu, B., & Wang, X. (2020). Dynamic Functional Connectivity Reveals Abnormal Variability and Hyper□connected Pattern in Autism Spectrum Disorder. Autism Research, 13(2), 230–243. 10.1002/aur.2212

Lindquist, M. A., Xu, Y., Nebel, M. B., & Caffo, B. S. (2014). Evaluating dynamic bivariate correlations in resting-state fMRI: A comparison study and a new approach. NeuroImage, 101, 531–546. 10.1016/j.neuroimage.2014.06.052

Lipkin, B., Tuckute, G., Affourtit, J., Small, H., Mineroff, Z., Kean, H., Jouravlev, O., Rakocevic, L., Pritchett, B., Siegelman, M., Hoeflin, C., Pongos, A., Blank, I. A., Struhl, M. K., Ivanova, A., Shannon, S., Sathe, A., Hoffmann, M., Nieto-Castañón, A., & Fedorenko, E. (2022). Probabilistic atlas for the language network based on precision fMRI data from >800 individuals. Scientific Data, 9(1), 529. 10.1038/s41597-022-01645-3

Lombardo, M. V., Pierce, K., Eyler, L. T., Carter Barnes, C., Ahrens-Barbeau, C., Solso, S., Campbell, K., & Courchesne, E. (2015). Different Functional Neural Substrates for Good and Poor Language Outcome in Autism. Neuron, 86(2), 567–577. 10.1016/j.neuron.2015.03.023

Lombardo, M. V., Pramparo, T., Gazestani, V., Warrier, V., Bethlehem, R. A. I., Carter Barnes, C., Lopez, L., Lewis, N. E., Eyler, L., Pierce, K., & Courchesne, E. (2018). Large-scale associations between the leukocyte transcriptome and BOLD responses to speech differ in autism early language outcome subtypes. Nature Neuroscience, 21(12), 1680–1688. 10.1038/s41593-018-0281-3

Lu, J., Zhao, Z., Zhang, J., Wu, B., Zhu, Y., Chang, E. F., Wu, J., Duffau, H., & Berger, M. S. (2021). Functional maps of direct electrical stimulation-induced speech arrest and anomia: A multicentre retrospective study. Brain, 144(8), 2541–2553. 10.1093/brain/awab125

Mash, L. E., Linke, A. C., Olson, L. A., Fishman, I., Liu, T. T., & Müller, R. (2019). Transient states of network connectivity are atypical in autism: A dynamic functional connectivity study. Human Brain Mapping, 40(8), 2377–2389. 10.1002/hbm.24529

Muller, A. M., & Meyer, M. (2014). Language in the brain at rest: New insights from resting state data and graph theoretical analysis. Frontiers in Human Neuroscience, 8. 10.3389/fnhum.2014.00228

Murphy, K., & Fox, M. D. (2017). Towards a consensus regarding global signal regression for resting state functional connectivity MRI. NeuroImage, 154, 169–173. 10.1016/j.neuroimage.2016.11.052

Nelson, B. G., Bassett, D. S., Camchong, J., Bullmore, E. T., & Lim, K. O. (2017). Comparison of large-scale human brain functional and anatomical networks in schizophrenia. NeuroImage: Clinical, 15, 439–448. 10.1016/j.nicl.2017.05.007

Olulade, O. A., Seydell-Greenwald, A., Chambers, C. E., Turkeltaub, P. E., Dromerick, A. W., Berl, M. M., Gaillard, W. D., & Newport, E. L. (2020). The neural basis of language development: Changes in lateralization over age. Proceedings of the National Academy of Sciences, 117(38), 23477–23483. 10.1073/pnas.1905590117

O’Muircheartaigh, J., Dean, D. C., Dirks, H., Waskiewicz, N., Lehman, K., Jerskey, B. A., & Deoni, S. C. L. (2013). Interactions between White Matter Asymmetry and Language during Neurodevelopment. The Journal of Neuroscience, 33(41), 16170–16177. 10.1523/JNEUROSCI.1463-13.2013

Paus, T., Perry, D. W., Zatorre, R. J., Worsley, K. J., & Evans, A. C. (1996). Modulation of Cerebral Blood Flow in the Human Auditory Cortex During Speech: Role of Motor□to□sensory Discharges. European Journal of Neuroscience, 8(11), 2236–2246. 10.1111/j.1460-9568.1996.tb01187.x

Philip, R. C. M., Dauvermann, M. R., Whalley, H. C., Baynham, K., Lawrie, S. M., & Stanfield, A. C. (2012). A systematic review and meta-analysis of the fMRI investigation of autism spectrum disorders. Neuroscience & Biobehavioral Reviews, 36(2), 901–942. 10.1016/j.neubiorev.2011.10.008

Pickles, A., Anderson, D. K., & Lord, C. (2014). Heterogeneity and plasticity in the development of language: A 17□year follow□up of children referred early for possible autism. Journal of Child Psychology and Psychiatry, 55(12), 1354–1362. 10.1111/jcpp.12269

Prescott, K. E., Mathée□Scott, J., Reuter, T., Edwards, J., Saffran, J., & Ellis Weismer, S. (2022). Predictive language processing in young autistic children. Autism Research, 15(5), 892–903. 10.1002/aur.2684

Prigge, M. B. D., Lange, N., Bigler, E. D., King, J. B., Dean, D. C., Adluru, N., Alexander, A. L., Lainhart, J. E., & Zielinski, B. A. (2021). A 16-year study of longitudinal volumetric brain development in males with autism. NeuroImage, 236, 118067. 10.1016/j.neuroimage.2021.118067

Rampinini, A. C., Handjaras, G., Leo, A., Cecchetti, L., Ricciardi, E., Marotta, G., & Pietrini, P. (2017). Functional and spatial segregation within the inferior frontal and superior temporal cortices during listening, articulation imagery, and production of vowels. Scientific Reports, 7(1), 17029. 10.1038/s41598-017-17314-0

Reindal, L., Nærland, T., Weidle, B., Lydersen, S., Andreassen, O. A., & Sund, A. M. (2023). Structural and Pragmatic Language Impairments in Children Evaluated for Autism Spectrum Disorder (ASD). Journal of Autism and Developmental Disorders, 53(2), 701–719. 10.1007/s10803-020-04853-1

Reynolds, J. E., Long, X., Grohs, M. N., Dewey, D., & Lebel, C. (2019). Structural and functional asymmetry of the language network emerge in early childhood. Developmental Cognitive Neuroscience, 39, 100682. 10.1016/j.dcn.2019.100682

Rice, G. E., Lambon Ralph, M. A., & Hoffman, P. (2015). The Roles of Left Versus Right Anterior Temporal Lobes in Conceptual Knowledge: An ALE Meta-analysis of 97 Functional Neuroimaging Studies. Cerebral Cortex, 25(11), 4374–4391. 10.1093/cercor/bhv024

Rubinov, M., & Sporns, O. (2010). Complex network measures of brain connectivity: Uses and interpretations. NeuroImage, 52(3), 1059–1069. 10.1016/j.neuroimage.2009.10.003

Schaeffer, J., Abd El-Raziq, M., Castroviejo, E., Durrleman, S., Ferré, S., Grama, I., Hendriks, P., Kissine, M., Manenti, M., Marinis, T., Meir, N., Novogrodsky, R., Perovic, A., Panzeri, F., Silleresi, S., Sukenik, N., Vicente, A., Zebib, R., Prévost, P., & Tuller, L. (2023). Language in autism: Domains, profiles and co-occurring conditions. Journal of Neural Transmission, 130(3), 433–457. 10.1007/s00702-023-02592-y

Schumann, C. M., Bloss, C. S., Barnes, C. C., Wideman, G. M., Carper, R. A., Akshoomoff, N., Pierce, K., Hagler, D., Schork, N., Lord, C., & Courchesne, E. (2010). Longitudinal Magnetic Resonance Imaging Study of Cortical Development through Early Childhood in Autism. The Journal of Neuroscience, 30(12), 4419–4427. 10.1523/JNEUROSCI.5714-09.2010

Seol, K. I., Song, S. H., Kim, K. L., Oh, S. T., Kim, Y. T., Im, W. Y., Song, D. H., & Cheon, K.-A. (2014). A Comparison of Receptive-Expressive Language Profiles between Toddlers with Autism Spectrum Disorder and Developmental Language Delay. Yonsei Medical Journal, 55(6), 1721. 10.3349/ymj.2014.55.6.1721

Shan, X., Wang, P., Yin, Q., Li, Y., Wang, X., Feng, Y., Xiao, J., Li, L., Huang, X., Chen, H., & Duan, X. (2025). Atypical dynamic neural configuration in autism spectrum disorder and its relationship to gene expression profiles. European Child & Adolescent Psychiatry, 34(1), 169–179. 10.1007/s00787-024-02476-w

Siegel, J. S., Ramsey, L. E., Snyder, A. Z., Metcalf, N. V., Chacko, R. V., Weinberger, K., Baldassarre, A., Hacker, C. D., Shulman, G. L., & Corbetta, M. (2016). Disruptions of network connectivity predict impairment in multiple behavioral domains after stroke. Proceedings of the National Academy of Sciences, 113(30). 10.1073/pnas.1521083113

Siller, M., & Sigman, M. (2008). Modeling longitudinal change in the language abilities of children with autism: Parent behaviors and child characteristics as predictors of change. Developmental Psychology, 44(6), 1691–1704. 10.1037/a0013771

Smith, P. F., & Zheng, Y. (2016). Cannabinoids, cannabinoid receptors and tinnitus. Hearing Research, 332, 210–216. 10.1016/j.heares.2015.09.014

Tager-Flusberg, H., Paul, R., & Lord, C. (2005). Language and Communication in Autism. In Handbook of autism and pervasive developmental disorders: Diagnosis, development, neurobiology, and behavior, Vol. 1, 3rd *ed.* (pp. 335–364). John Wiley & Sons, Inc.

Tesink, C. M. J. Y., Buitelaar, J. K., Petersson, K. M., Van Der Gaag, R. J., Kan, C. C., Tendolkar, I., & Hagoort, P. (2009). Neural correlates of pragmatic language comprehension in autism spectrum disorders. Brain, 132(7), 1941–1952. 10.1093/brain/awp103

The IBIS Network, Hazlett, H. C., Gu, H., Munsell, B. C., Kim, S. H., Styner, M., Wolff, J. J., Elison, J. T., Swanson, M. R., Zhu, H., Botteron, K. N., Collins, D. L., Constantino, J. N., Dager, S. R., Estes, A. M., Evans, A. C., Fonov, V. S., Gerig, G., Kostopoulos, P., … Piven, J. (2017). Early brain development in infants at high risk for autism spectrum disorder. Nature, 542(7641), 348–351. 10.1038/nature21369

Thurm, A., Manwaring, S. S., Swineford, L., & Farmer, C. (2015). Longitudinal study of symptom severity and language in minimally verbal children with autism. Journal of Child Psychology and Psychiatry, 56(1), 97–104. 10.1111/jcpp.12285

Tipping, M. E. (1999). The Relevance Vector Machine. Advances in Neural Information Processing Systems, 12, 652–658.

Van Den Heuvel, M. P., & Sporns, O. (2011). Rich-Club Organization of the Human Connectome. The Journal of Neuroscience, 31(44), 15775–15786. 10.1523/JNEUROSCI.3539-11.2011

Vigneau, M., Beaucousin, V., Hervé, P. Y., Duffau, H., Crivello, F., Houdé, O., Mazoyer, B., & Tzourio-Mazoyer, N. (2006). Meta-analyzing left hemisphere language areas: Phonology, semantics, and sentence processing. NeuroImage, 30(4), 1414–1432. 10.1016/j.neuroimage.2005.11.002

Wang, H., Ma, Z.-H., Xu, L.-Z., Yang, L., Ji, Z.-Z., Tang, X.-Z., Liu, J.-R., Li, X., Cao, Q.-J., & Liu, J. (2022). Developmental brain structural atypicalities in autism: A voxel-based morphometry analysis. Child and Adolescent Psychiatry and Mental Health, 16(1), 7. 10.1186/s13034-022-00443-4

Wei, D., Zhuang, K., Ai, L., Chen, Q., Yang, W., Liu, W., Wang, K., Sun, J., & Qiu, J. (2018). Structural and functional brain scans from the cross-sectional Southwest University adult lifespan dataset. Scientific Data, 5(1), 180134. 10.1038/sdata.2018.134

Whitney, O., Soderstrom, K., & Johnson, F. (2003). CB1 cannabinoid receptor activation inhibits a neural correlate of song recognition in an auditory/perceptual region of the zebra finch telencephalon. Journal of Neurobiology, 56(3), 266–274. 10.1002/neu.10233

Williams, G. L. (2021). Theory of autistic mind: A renewed relevance theoretic perspective on so-called autistic pragmatic ‘impairment’. Journal of Pragmatics, 180, 121–130. 10.1016/j.pragma.2021.04.032

Xia, M., Si, T., Sun, X., Ma, Q., Liu, B., Wang, L., Meng, J., Chang, M., Huang, X., Chen, Z., Tang, Y., Xu, K., Gong, Q., Wang, F., Qiu, J., Xie, P., Li, L., & He, Y. (2019). Reproducibility of functional brain alterations in major depressive disorder: Evidence from a multisite resting-state functional MRI study with 1,434 individuals. NeuroImage, 189, 700–714. 10.1016/j.neuroimage.2019.01.074

Xiao, Y., Wen, T. H., Kupis, L., Eyler, L. T., Goel, D., Vaux, K., Lombardo, M. V., Lewis, N. E., Pierce, K., & Courchesne, E. (2022). Neural responses to affective speech, including motherese, map onto clinical and social eye tracking profiles in toddlers with ASD. Nature Human Behaviour, 6(3), 443–454. 10.1038/s41562-021-01237-y

Xie, Y., Xu, Z., Xia, M., Liu, J., Shou, X., Cui, Z., Liao, X., & He, Y. (2022). Alterations in Connectome Dynamics in Autism Spectrum Disorder: A Harmonized Mega- and Meta-analysis Study Using the Autism Brain Imaging Data Exchange Dataset. Biological Psychiatry, 91(11), 945–955. 10.1016/j.biopsych.2021.12.004

Yang, J., Hu, Z., Li, J., Guo, X., Gao, X., Liu, J., Wang, Y., Qu, Z., Li, W., Li, Z., Li, W., Huang, Y., Chen, J., Wen, H., & Yuan, B. (2025). NaDyNet: A toolbox for dynamic network analysis of naturalistic stimuli. NeuroImage, 311, 121203. 10.1016/j.neuroimage.2025.121203

Yuan, B., Xie, H., Gong, F., Zhang, N., Xu, Y., Zhang, H., Liu, J., Chen, L., Li, C., Tan, S., Lin, Z., Hu, X., Gu, T., Cheng, J., Lu, J., Liu, D., Wu, J., & Yan, J. (2023). Dynamic network reorganization underlying neuroplasticity: The deficits-severity-related language network dynamics in patients with left hemispheric gliomas involving language network. Cerebral Cortex, 33(13), 8273–8285. 10.1093/cercor/bhad113

Yuan, B., Xie, H., Wang, Z., Xu, Y., Zhang, H., Liu, J., Chen, L., Li, C., Tan, S., Lin, Z., Hu, X., Gu, T., Lu, J., Liu, D., & Wu, J. (2023). The domain-separation language network dynamics in resting state support its flexible functional segregation and integration during language and speech processing. NeuroImage, 274, 120132. 10.1016/j.neuroimage.2023.120132

Yuan, B., Zhang, N., Gong, F., Wang, X., Yan, J., Lu, J., & Wu, J. (2022). Longitudinal assessment of network reorganizations and language recovery in postoperative patients with glioma. Brain Communications, 4(2), fcac046. 10.1093/braincomms/fcac046

Yuan, B., Zhang, N., Yan, J., Cheng, J., Lu, J., & Wu, J. (2019). Resting-state functional connectivity predicts individual language impairment of patients with left hemispheric gliomas involving language network. NeuroImage: Clinical, 24, 102023. 10.1016/j.nicl.2019.102023

Zalesky, A., Fornito, A., & Bullmore, E. T. (2010). Network-based statistic: Identifying differences in brain networks. NeuroImage, 53(4), 1197–1207. 10.1016/j.neuroimage.2010.06.041

Zhang, G., Xu, Y., Wang, X., Li, J., Shi, W., Bi, Y., & Lin, N. (2023). A social-semantic working-memory account for two canonical language areas. Nature Human Behaviour, 7(11), 1980–1997. 10.1038/s41562-023-01704-8

Zhang, J., Shang, D., Ye, J., Ling, Y., Zhong, S., Zhang, S., Zhang, W., Zhang, L., Yu, Y., He, F., Ye, X., & Luo, B. (2022). Altered Coupling Between Cerebral Blood Flow and Voxel-Mirrored Homotopic Connectivity Affects Stroke-Induced Speech Comprehension Deficits. Frontiers in Aging Neuroscience, 14, 922154. 10.3389/fnagi.2022.922154

