## Supplementary Figure 1, Supplementary Figure 2, Supplementary Figure 3, Supplementary Figure 4, Supplementary Figure 5, Supplementary Figure 6 for "The aberrant language network dynamics in autism ages 5–60 years"

**Supplementary Methods**

Nodal strength and distribution. The nodal strength for a given node was calculated by summing up the correlation coefficients of all the suprathreshold connections that connected with this node. To determine whether there were hubs in each state, we calculated the nodal strength distribution $P (k)$*.* Three possible forms of distribution were fitted to the probability of nodal strength: a power-law, $P \left( x \right) \sim x^{\alpha-1}$; and exponential, $P \left( x \right) \sim exp(\frac{x}{x_{c}})$; and an exponentially truncated power-law, $P \left( x \right) \sim x^{\alpha-1}exp(\frac{x}{x_{c}})$. An exponentially truncated power-law distribution is expected for human brain networks, which denotes a long-tailed broad-scale topologies and a large proportion of network connectivity will be concentrated on a subset node (i.e., hubs) (Achard, Salvador, Whitcher, Suckling, & Bullmore, 2006; Gong et al., 2009; He et al., 2009).

**Supplementary Figures**

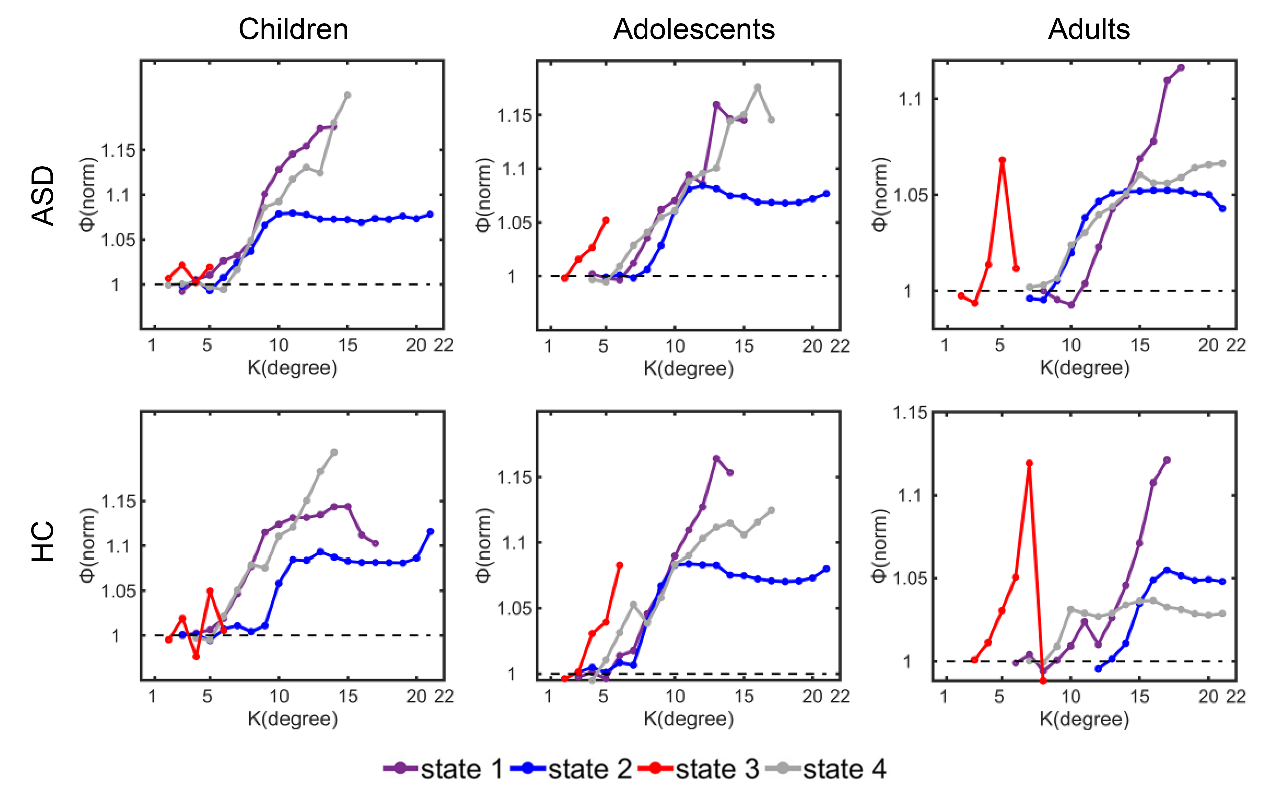

**Supplementary Figure 1. The rich-club organizations.** The normalized rich-club curves were calculated on the weighted network thresholded with a correlation strength of 0.2. The rich-club organizations, i.e., the ranges of K with the normalized rich-club coefficients larger than 1 were state-dependent.

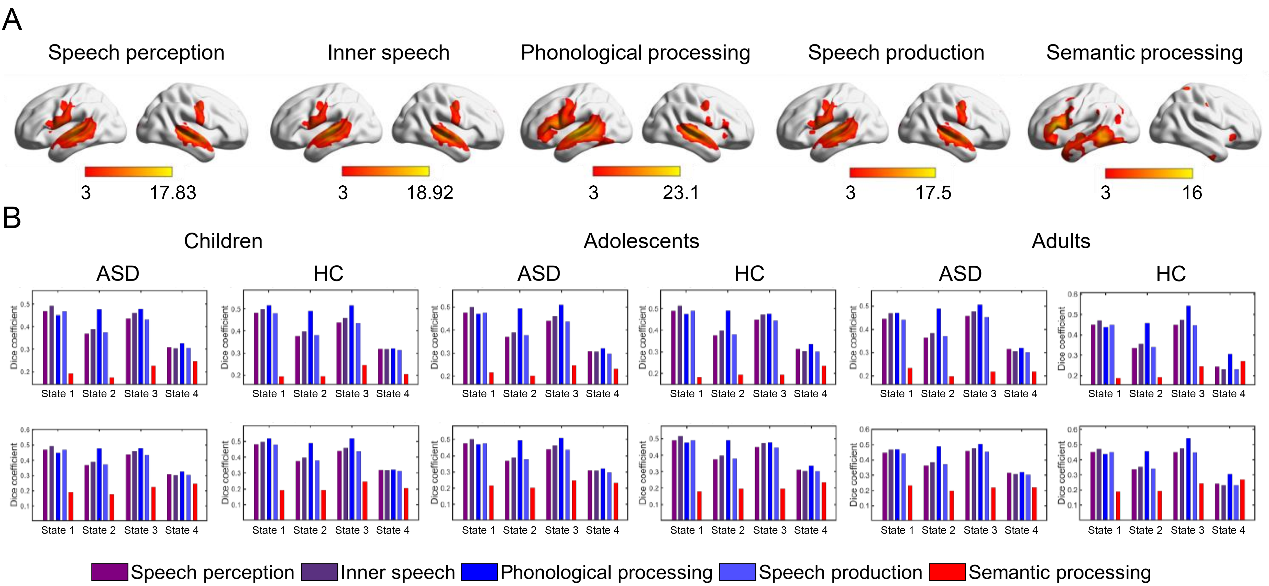

**Supplementary Figure 2. Functional relevance of hubs in the first three states.** A. The meta results of speech perception, inner speech, phonological processing, speech production, and semantic processing, were from‘NeuroQuery’ ([*https://neuroquery.org/*](https://neuroquery.org/)) (Dockes et al., 2020). Each of these maps was thresholded at *Z* = 3 (a typical value used by NeuroQuery) for illustrative purposes and only positive results were shown. B–D: the dice coefficients between binary images of nodes and meta results. Considering the left-lateralized activations of meta results, the dice coefficients were calculated both at the whole-brain level and in the left hemisphere.

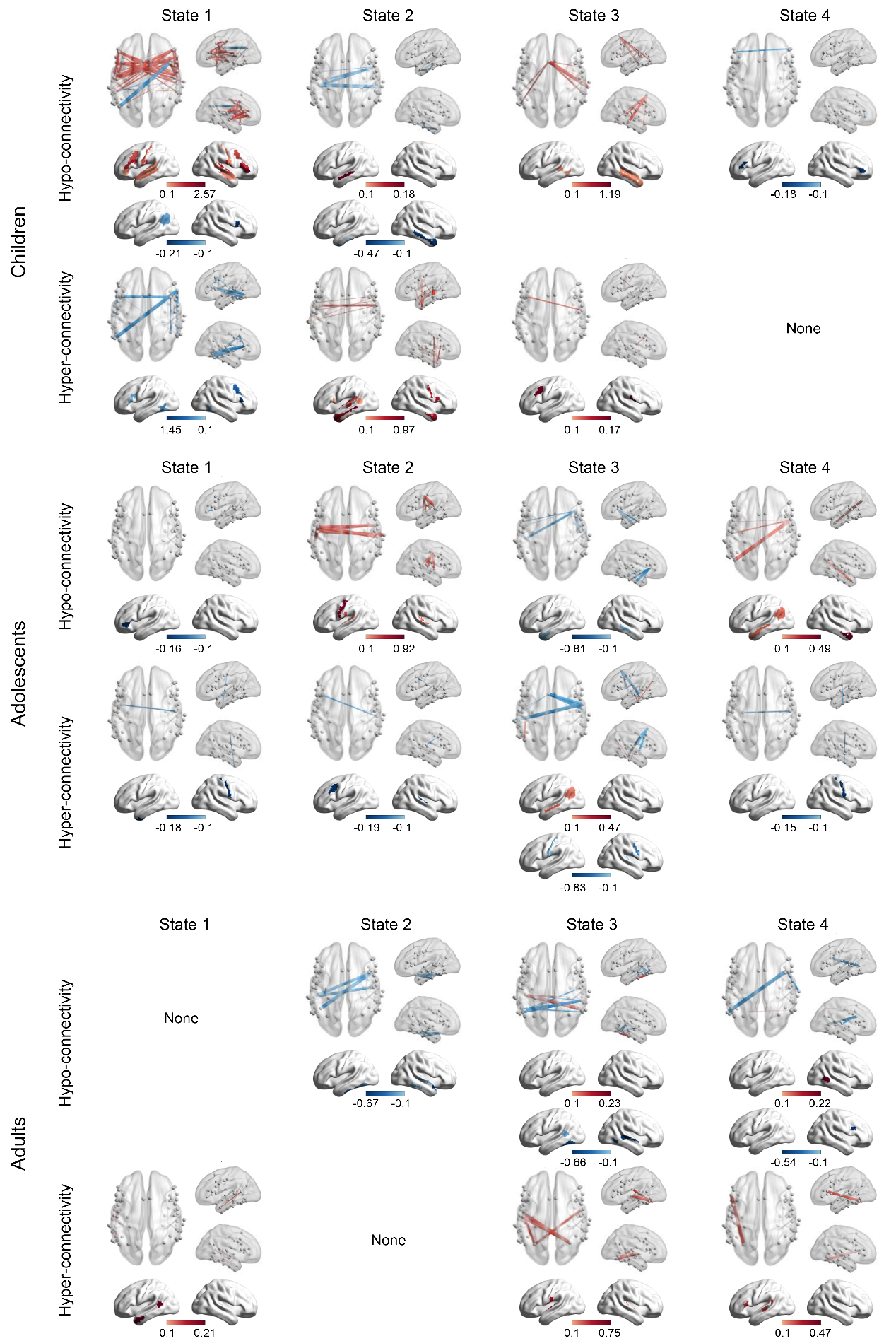

**Supplementary Figure 3. Partial correlations between VIQ and functional connectivity differed between HCs and ASDs.** The blue lines indicate a negative correlation with VIQ scores, while the red lines indicate a positive correlation. Pearson correlation coefficients were calculated using sex, age, and FIQ as covariates, and only edges with a significant correlation (*P* < 0.05 with NBS correction) with behavioral data were retained.

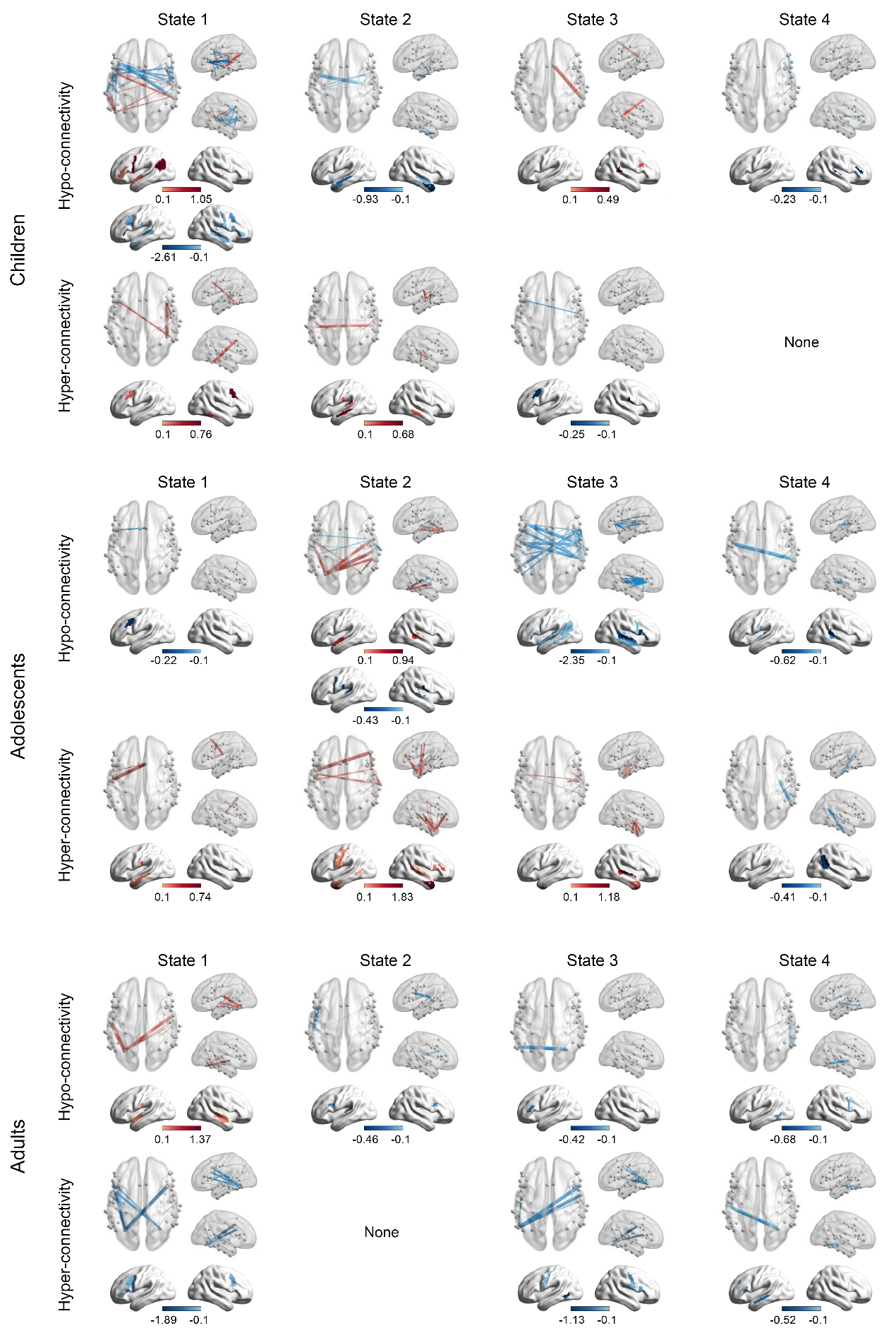

**Supplementary Figure 4. Partial correlation between ADOS communication scores and functional connectivity differed between HCs and ASDs.**

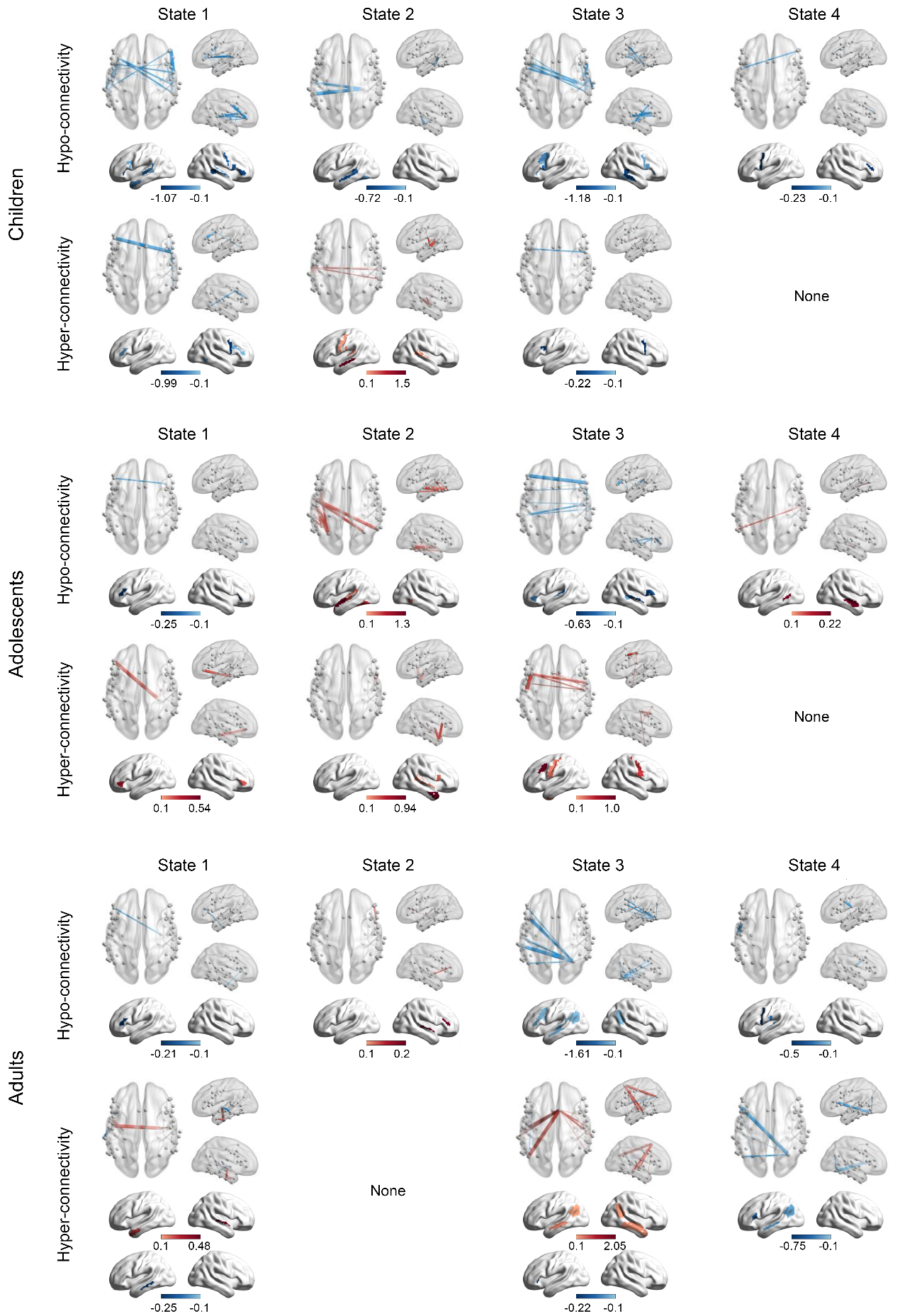

**Supplementary Figure 5. Partial correlation between ADOS social scores and functional connectivity differed between HCs and ASDs.**

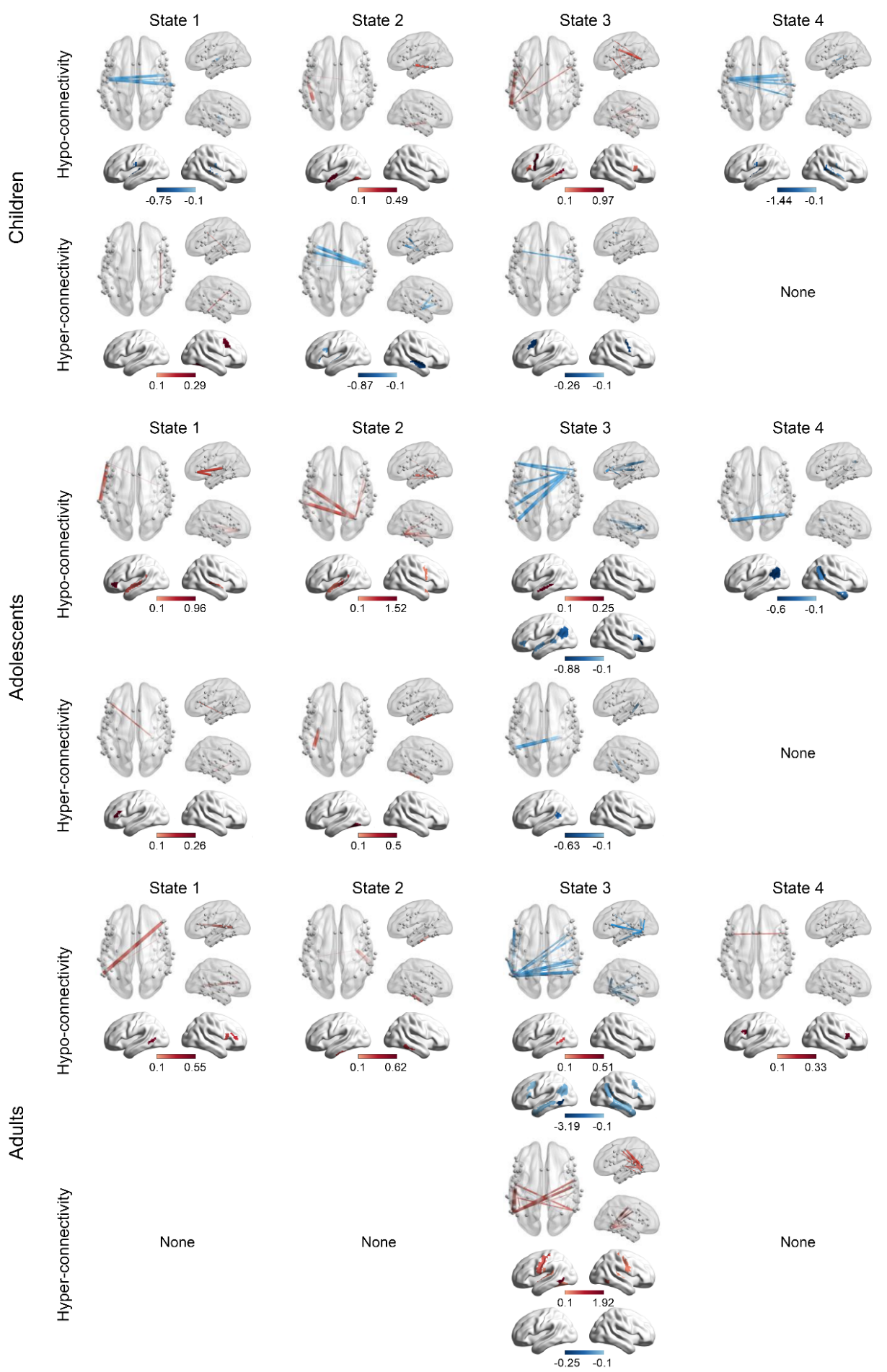

**Supplementary Figure 6. Partial correlation between ADOS RRB scores and functional connectivity differed between HCs and ASDs.**

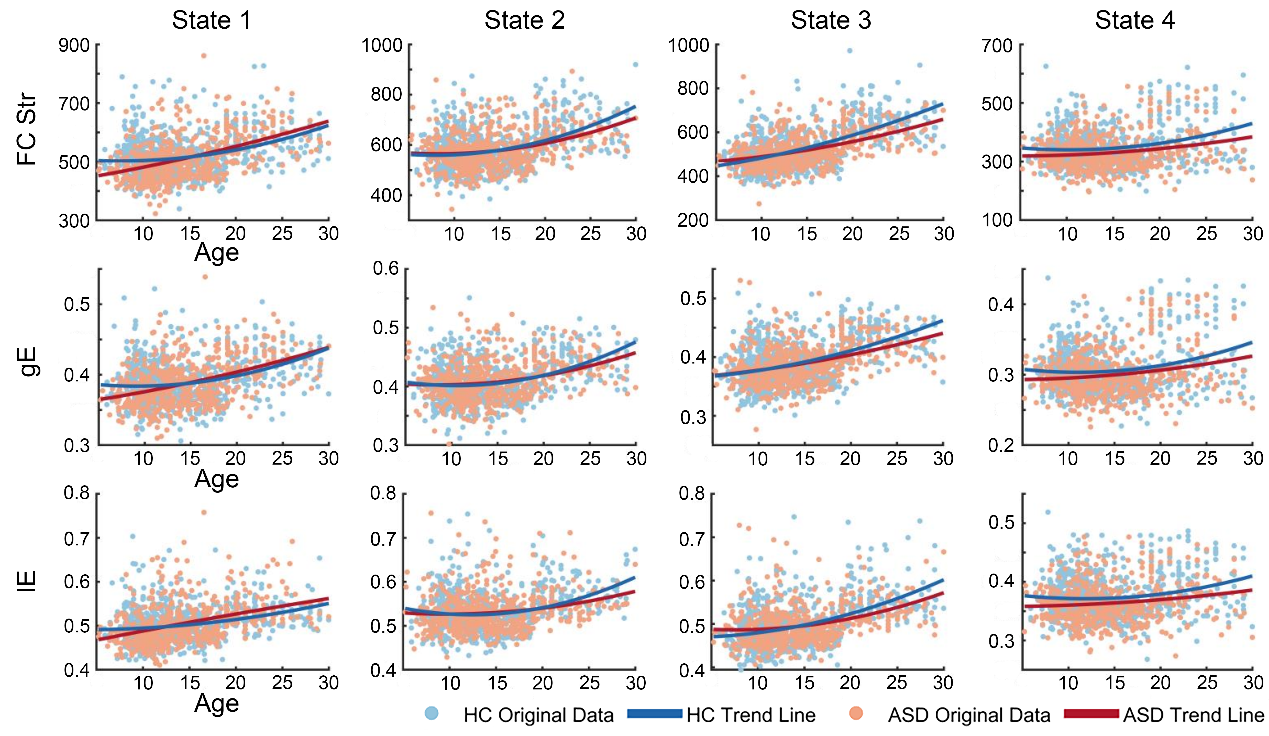

**Supplementary Figure 7. Domain-specific development trajectories of language networks fitted using polynomial functions.** Graphs show the original data (light blue for HC, light red for ASD), with the fitted trajectories superimposed (solid blue for HC, light red for ASD) in individuals under 30 years old.

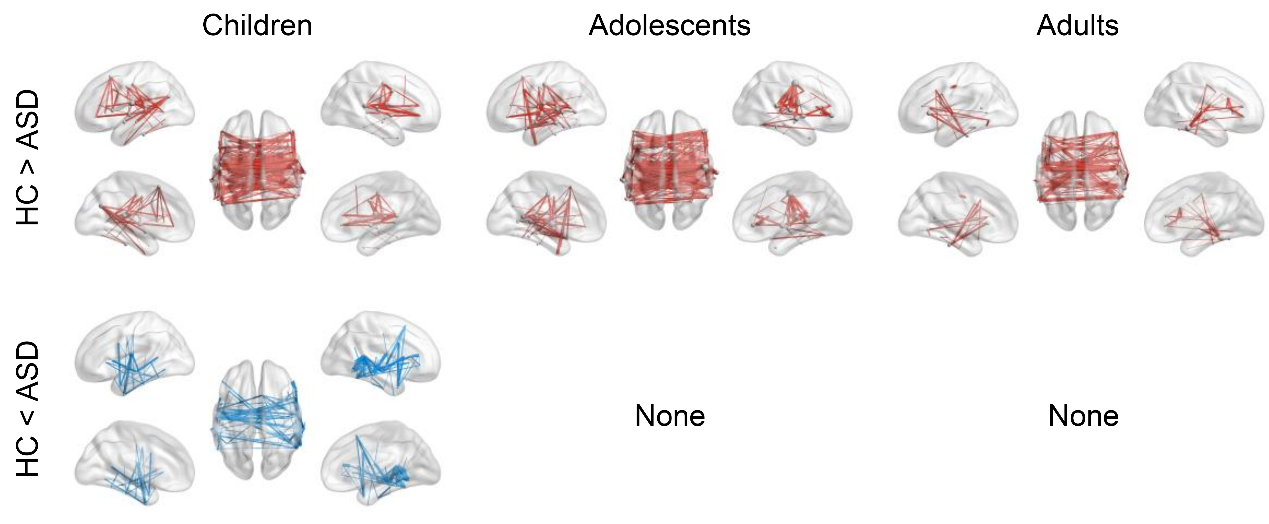

**Supplementary Figure 8. Group differences of sFC.** The results have been corrected using NBS (Edge *P* < 0.01, Component *P* < 0.05). Red connections indicate HC > ASD, while blue connections indicate HC < ASD.

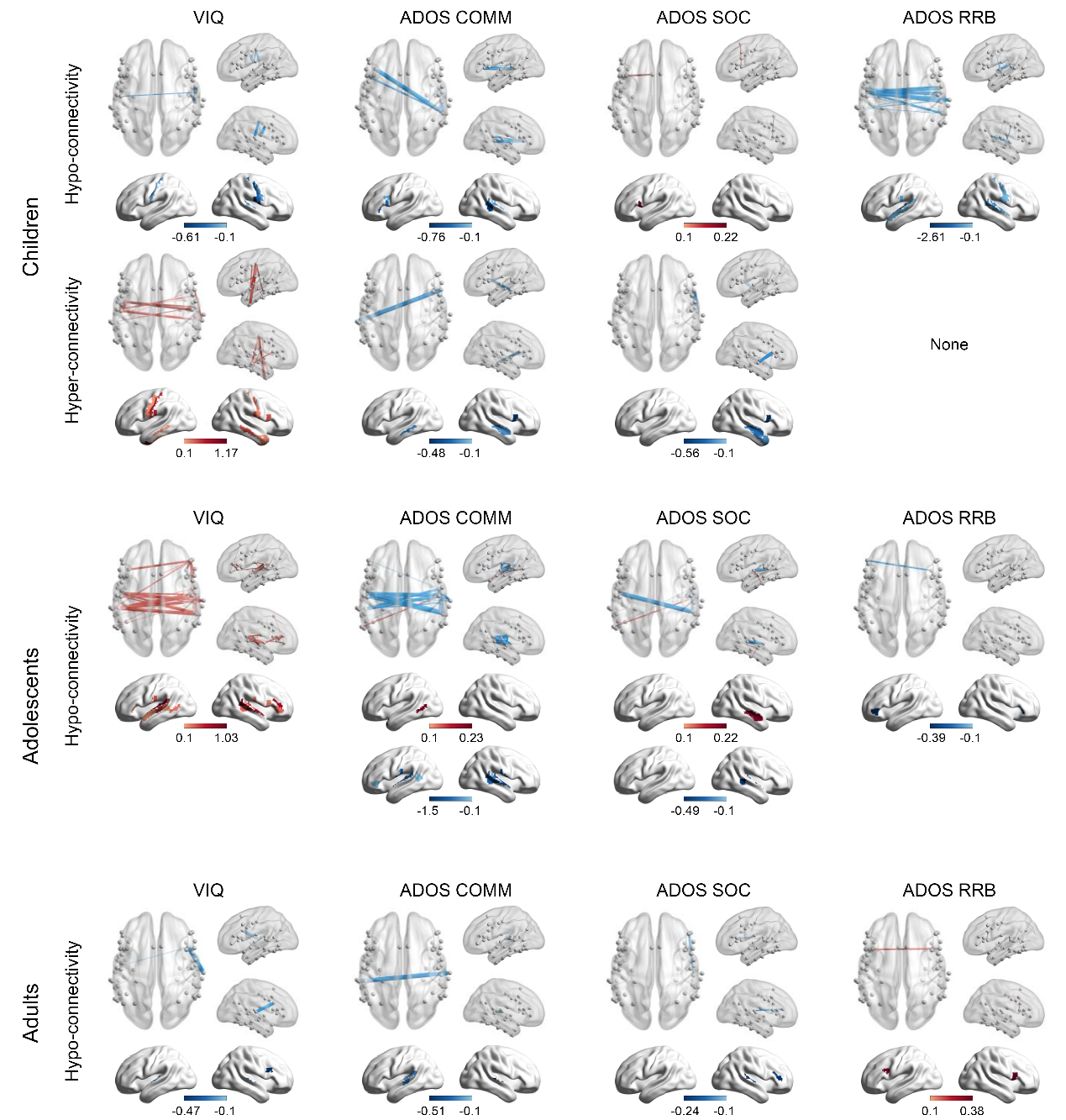

**Supplementary Figure 9.** **Partial correlation between behavioral scores and sFC differed between HCs and ASDs**.

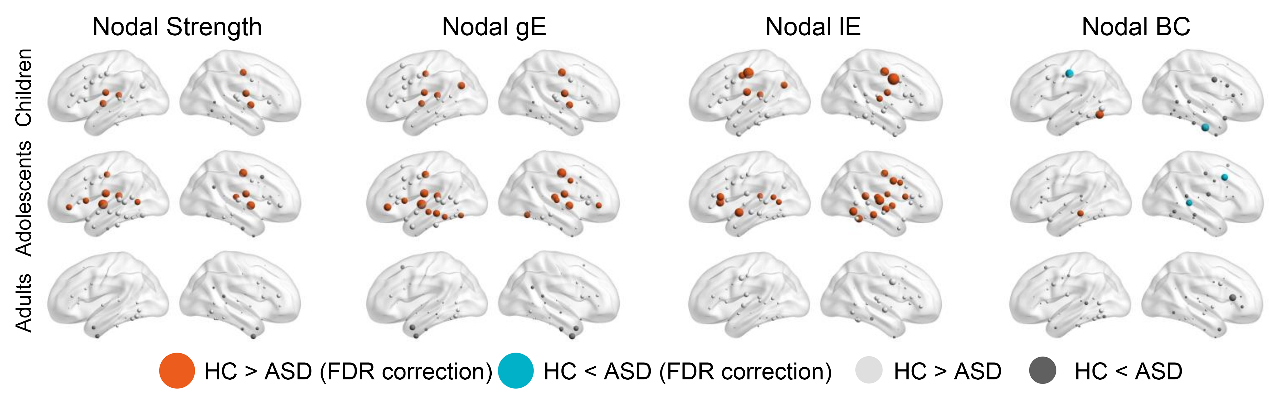

**Supplementary Figure 10.** **Changes of** **nodal** **topological characteristics of sFC**. Four nodal topological properties, the nodal Degree Centrality(DC), the nodal Between Centrality(BC), the nodal global efficiencies (gE), and local efficiencies (lE) were calculated for each age of sFC. Between-group statistical comparisons were performed by using two-sample t-tests (FDR correction, *P* < 0.05).

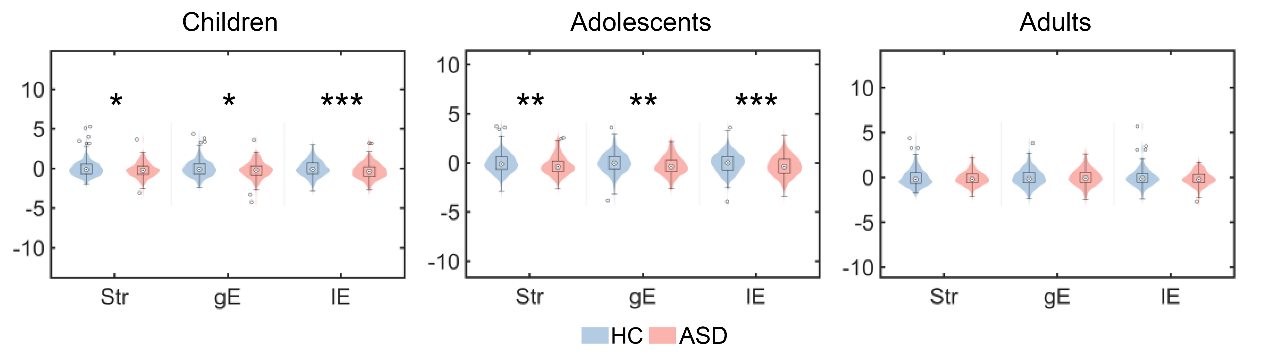

**Supplementary Figure 11.** **Changes of global topological characteristics of sFC**. Three network topological properties, the total functional connectivity strength (Str), the network global (gE), and local efficiencies (lE) were calculated. For illustration purposes, the topological values were z-scaled. Between-group statistical comparisons were performed by using Mann-Whitney-Wilcoxon nonparametric test paired t-tests. *, uncorrected *P* < 0.05.

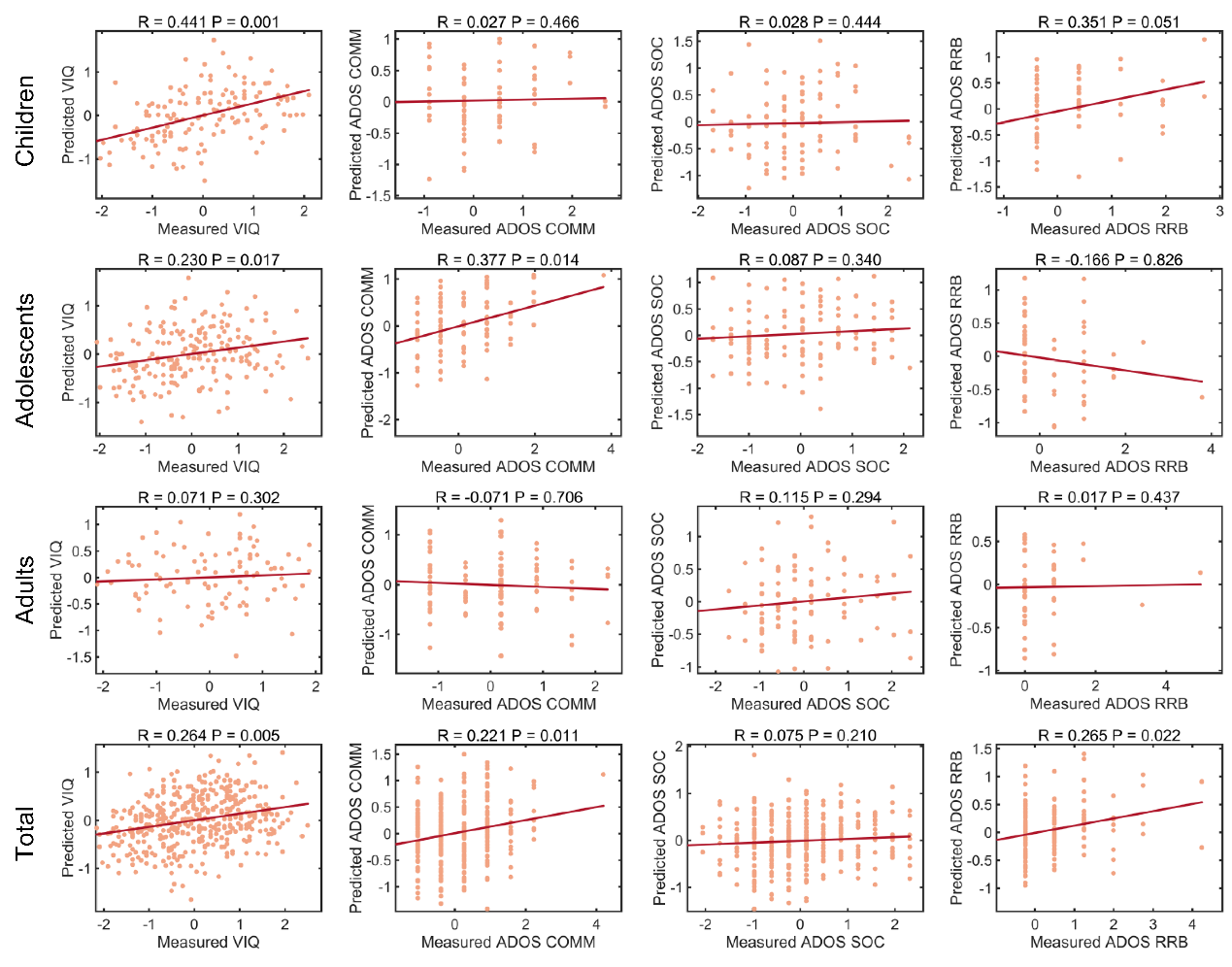

**Supplementary Figure 12.** **The accuracies and significance of sFC-based prediction models.** The scatter plots showed real (normalized within each group) and predicted scores and the corresponding linear fitted lines. The *R* value was the Pearson correlation coefficient between the predicted values and the actual values, and the model significance and the *P* values were based on 1000 permutation tests.

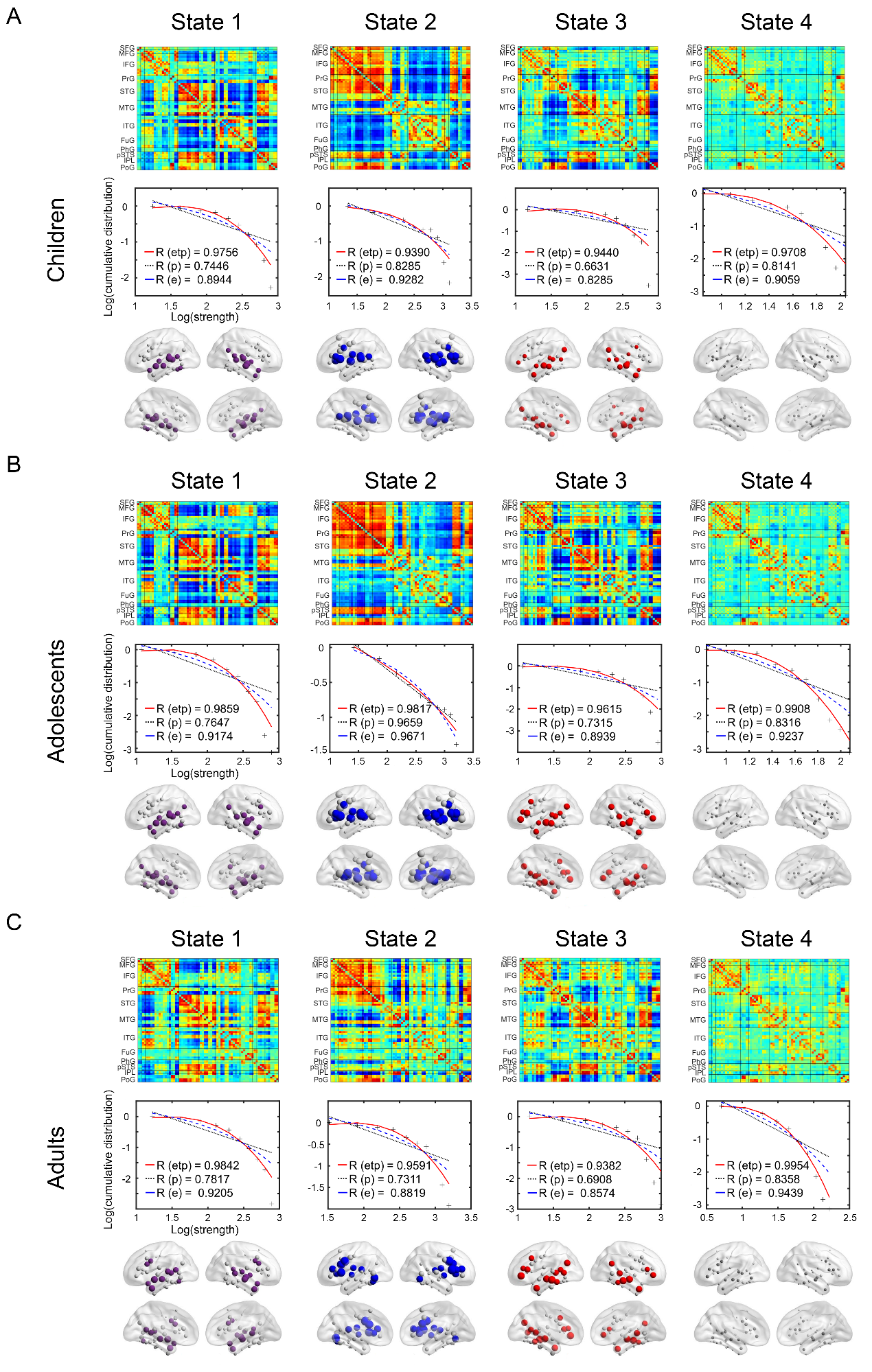

**Supplementary Figure 13. Cortical language network dynamics in the external validation cohort.** Figures A–C illustrate the four language network states in HCs in the external validation cohort: (A) children, (B) adolescents, and (C) adults.

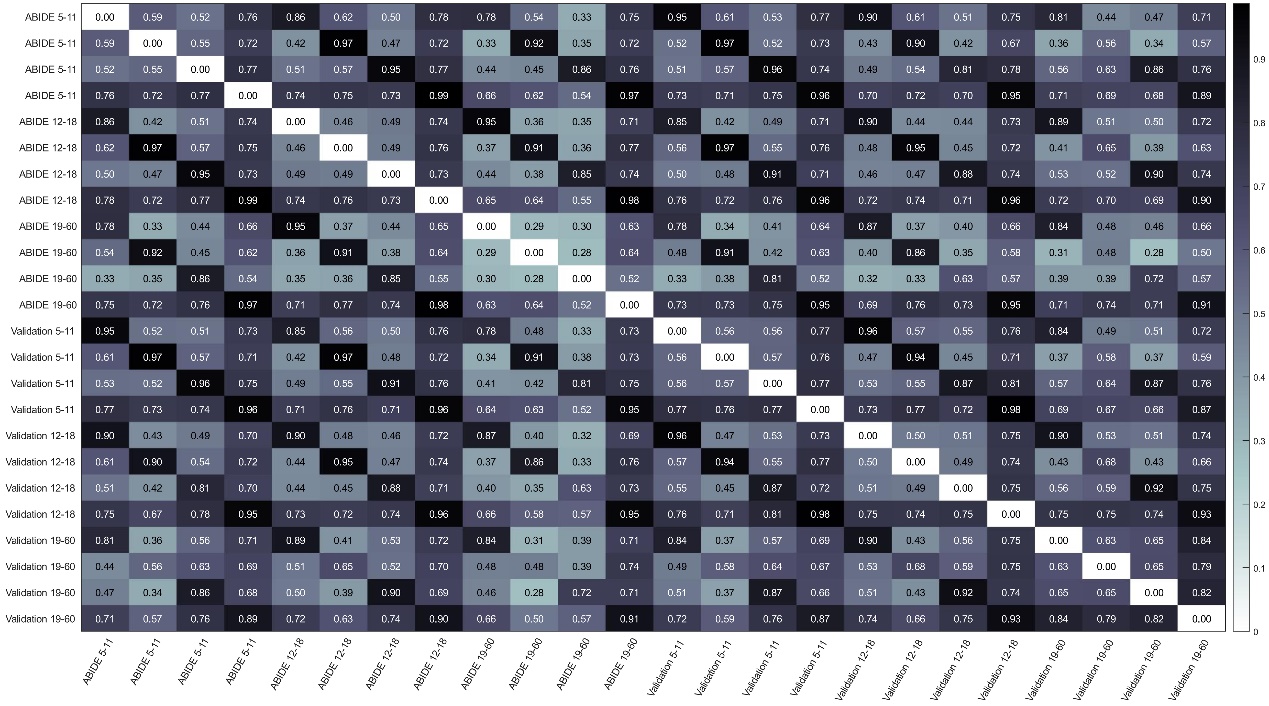

**Supplementary Figure 14. Spatial similarity of language network state connectivity patterns between ABIDE and the external validation cohort.** The figure shows pairwise similarity between language network state connectivity patterns derived from the ABIDE cohort and the validation cohort. For each state, the functional connectivity matrix was vectorized and compared using correlation coefficients; higher values indicate greater similarity in connectivity organization. Cell values denote the corresponding similarity coefficients, and the color bar indicates the magnitude of similarity. Axis labels specify cohort, age range, and state.

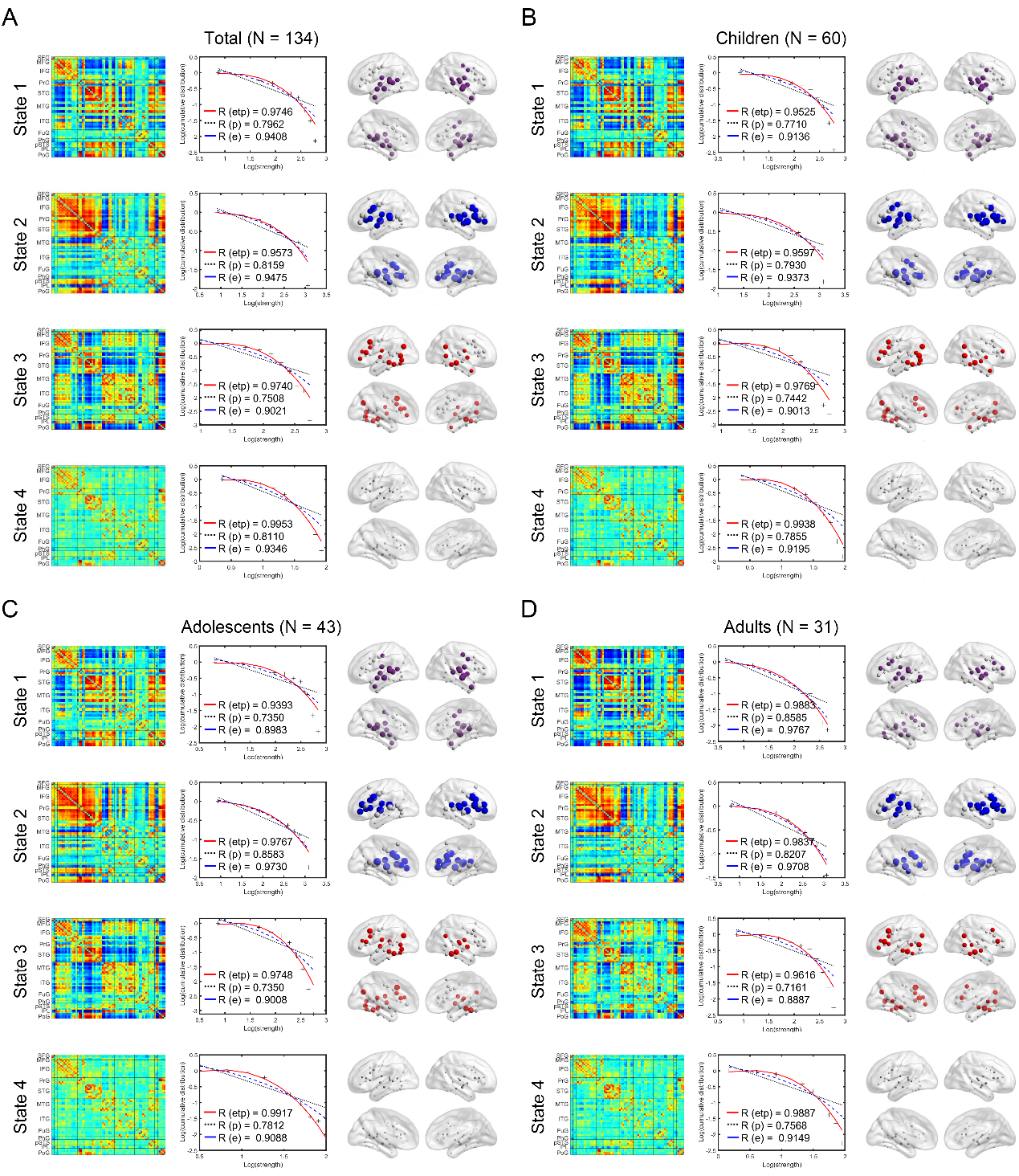

**Supplementary Figure 15. Cortical language network dynamics in HCs at the NYU site.** Figures A–D illustrate the four language network states in HCs at the NYU site: (A) all participants, (B) children, (C) adolescents, and (D) adults.

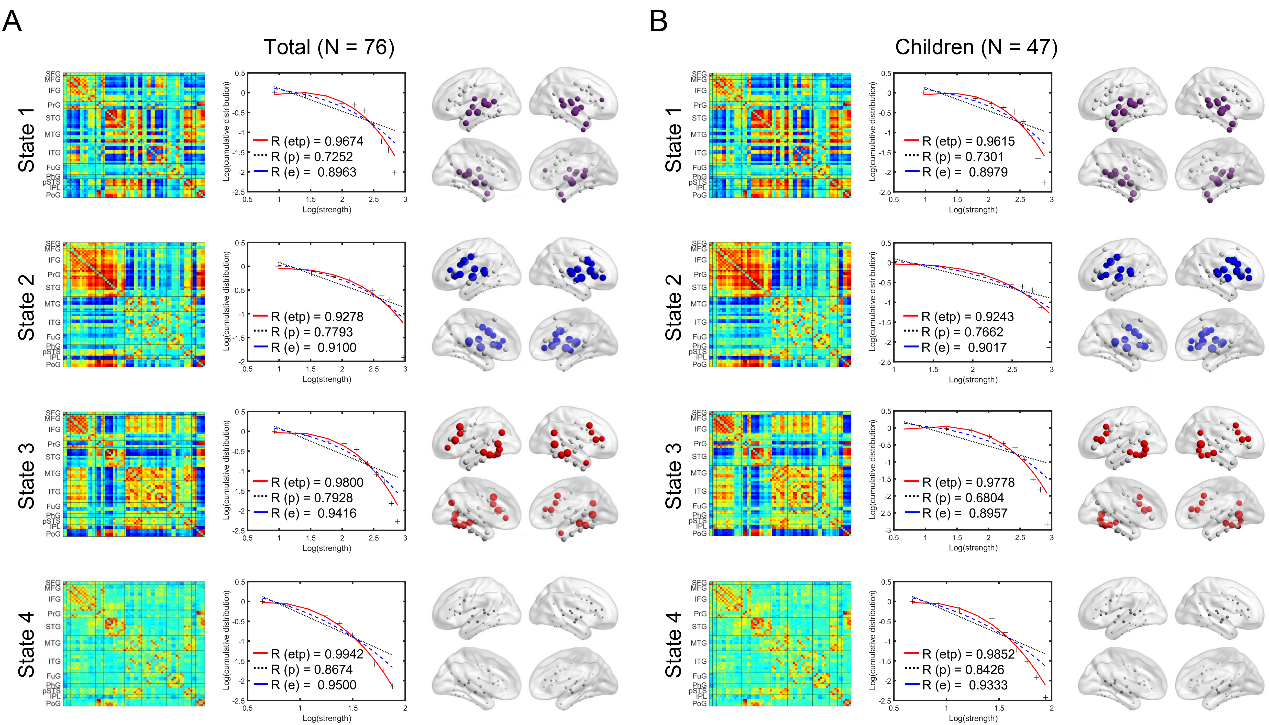

**Supplementary Figure 16. Cortical language network dynamics in ASDs at the NYU site.** Figures A–B illustrate the four language network states in ASDs at the NYU site: (A) all participants, (B) children.

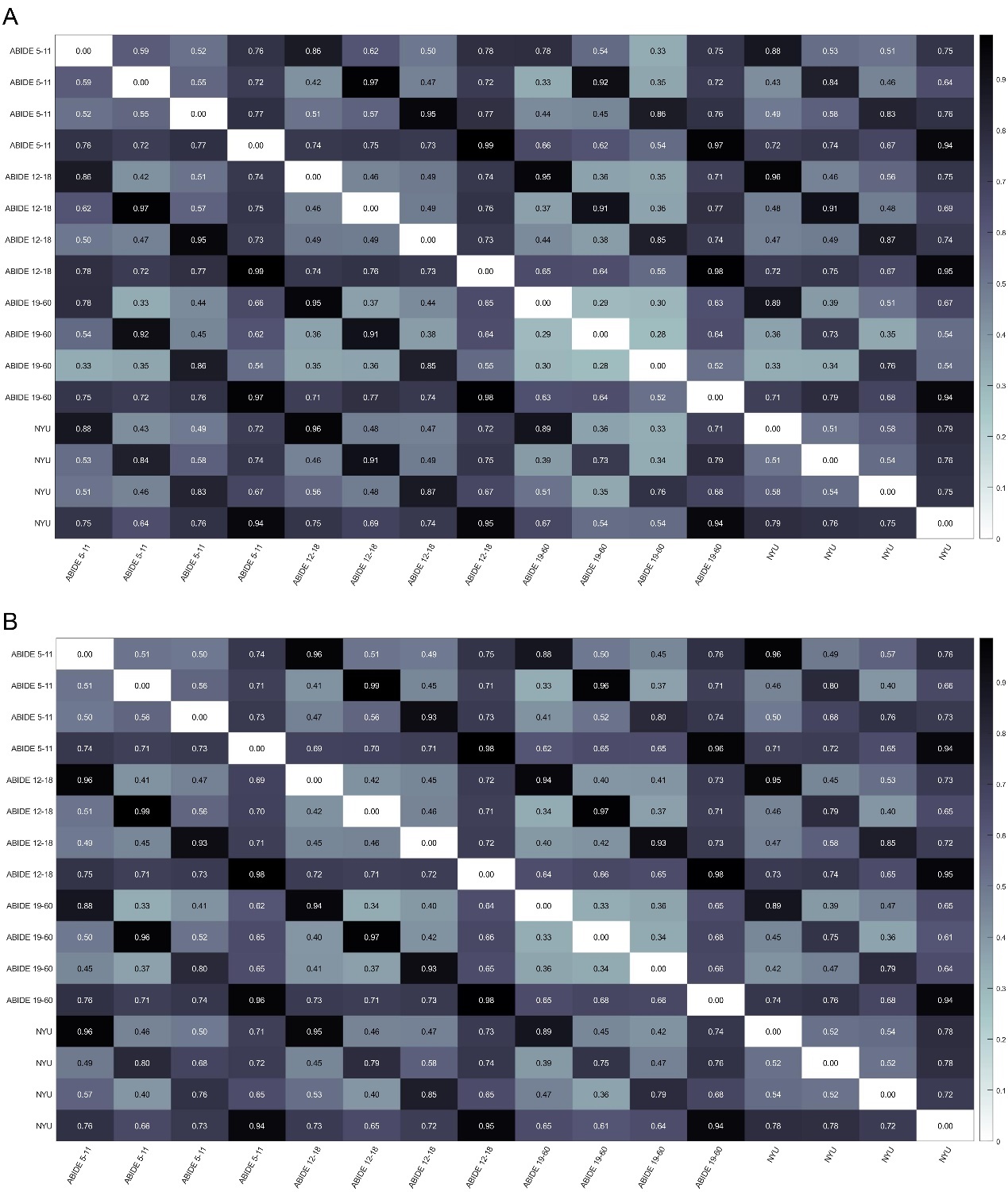

**Supplementary Figure 17. Spatial similarity of language network state connectivity patterns between ABIDE and the NYU site.** Figures A–B show pairwise similarity between language network state connectivity patterns derived from the ABIDE cohort and the NYU cohort. (A) HC, (B) ASD.

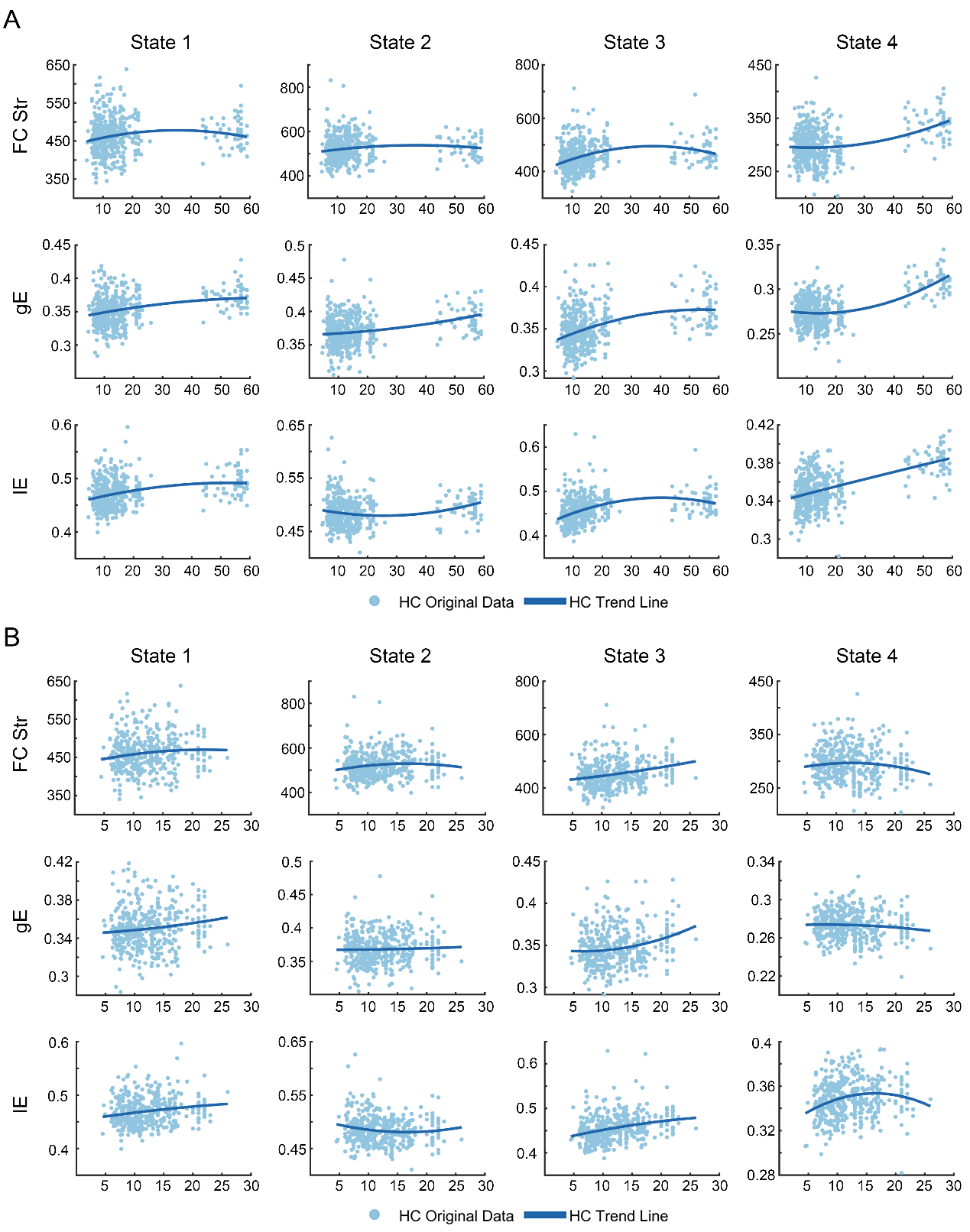

**Supplementary Figure 18. Domain-specific development trajectories of language networks in the external validation cohort.** Figures A–B show the individual data (light blue), with the fitted trajectories superimposed (solid blue). (A) displays the full age range, (B) focuses on the individuals under 30 years old.

**Supplementary tables**

Supplementary Table 1. Site information of the analyzed ABIDE consortium

| Sites | From | Children (5–11) | | Adolescents (12–18) | | Adults (19–40) | |
| --- | --- | --- | --- | --- | --- | --- | --- |
|  |  | ASD (F / M)  *n* = 198  (18 / 88) | HC (F / M)  *n* = 349  (76 / 123) | ASD (F / M)  *n* = 260  (14 / 107) | HC (F / M)  *n* = 293  (27 / 116) | ASD (F / M)  *n* = 166  (9 / 89) | HC (F / M)  *n* = 224  (17 / 104) |
| BNI | ABIDE 2 | None | None | 2 (0 / 2) | 1 (0 / 1) | 24 (0 / 24) | 27 (0 / 27) |
| Caltech | ABIDE 1 | None | None | 1 (0 / 1) | 2 (0 / 2) | 18 (4 / 14) | 16 (4 / 12) |
| EMC | ABIDE 2 | 16 (4 / 12) | 15 (3 / 12) | None | None | None | None |
| GU | ABIDE 2 | 21 (4 / 17) | 36 (19 / 17) | 9 (0 / 9) | 13 (6 / 7) | None | None |
| IU | ABIDE 2 | None | None | 2 (0 / 2) | None | 16 (4 / 12) | 18 (5 / 13) |
| KKI | ABIDE 1&2 | 31 (8 / 23) | 120 (46 / 74) | 7 (1 / 6) | 13 (2 / 11) | None | None |
| LEUVEN | ABIDE 1 | None | None | 12 (1 / 11) | 16 (2 / 14) | 12 (0 / 12) | 13 (0 / 13) |
| MAX | ABIDE 1 | 6 (0 / 6) | 2 (0 / 2) | 2 (0 / 2) | None | 11 (2 / 9) | 27 (4 / 23) |
| NYU | ABIDE 1&2 | 47 (3 / 44) | 60 (10 / 50) | 17 (3 / 14) | 43 (10 / 33) | 12 (2 / 10) | 31 (8 / 23) |
| OHSU | ABIDE 1&2 | 19 (3 / 16) | 54 (23 / 31) | 26 (3 / 23) | 14 (6 / 8) | None | None |
| OILH | ABIDE 2 | None | None | 3 (0 / 3) | 1 (1 / 0) | 16 (2 / 14) | 30 (12/ 18) |
| OLIN | ABIDE 1 | 1 (0 / 1) | 1 (0 / 1) | 9 (1 / 8) | 7 (1 / 6) | 4 (2 / 2) | 4 (1 / 3) |
| PITT | ABIDE 1 | 2 (1 / 1) | 2 (0 / 2) | 13 (3 / 10) | 10 (3 / 7) | 10 (0 / 10) | 7 (1 / 6) |
| SDSU | ABIDE 1&2 | 13 (2 / 11) | 11 (2 / 9) | 24 (3 / 21) | 31 (6 / 25) | None | None |
| TRINITY | ABIDE 1&2 | 4 (0 / 4) | 2 (0 / 2) | 24 (0 / 24) | 25 (0 / 25) | 9 (0 / 9) | 15 (0 / 15) |
| UCLA | ABIDE 1&2 | 21 (3 / 18) | 24 (5 / 19) | 30 (2 / 28) | 30 (6 / 24) | None | None |
| UCD | ABIDE 2 | None | None | 15 (4 / 11) | 13 (4 / 9) | None | None |
| UM | ABIDE 1 | 12 (4 / 8) | 15 (3 / 12) | 34 (5 / 29) | 44 (12 / 32) | None | 3 (1 / 2) |
| USM | ABIDE 1&2 | 1 (0 / 1) | 4 (0 / 4) | 23 (0 / 23) | 20 (0 / 20) | 34 (2 / 32) | 33 (3 / 30) |
| YALE | ABIDE 1 | 4 (1 / 3) | 3 (1 / 2) | 7 (4 / 3) | 10 (4 / 6) | None | None |

BNI, Barrow Neurological Institute; Caltech, California Institute of Technology; EMC, Erasmus University Medical Center Rotterdam; GU, Georgetown University; IU, Indiana University; KKI, Kennedy Krieger Institute; LEUVEN, University of Leven; MAX, Ludwig Maximilians University Munich; NYU, NYU Langone Medical Center; OHSU, Oregon Health and Science University; OLIN, OILH, Institute of Living at Hartford Hospital; PITT, University of Pittsburgh School of Medicine; SDSU, San Diego State University; TRINITY, Trinity Centre for Health Sciences; UCLA, University of California, Los Angeles; UCD, University of California, Davis; UM, University of Michigan; USM, University of Utah School of Medicine; YALE, Yale Child Study Center;

Supplementary Table 2. Anatomical regions，language-related behavioral domains, and paradigm classes of the language network. Coordinates are in the standard Montreal Neurologic Institute Space.

| ID | x | y | z | anatomy | behavioral domains | paradigm classes |
| --- | --- | --- | --- | --- | --- | --- |
| 1 | -4.6 | 15.7 | 53.2 | SFG | Execution. Speech, Phonology, Semantics, Speech | Word Generation (Covert and Overt) |
| 2 | 7.0 | 16.4 | 54.7 | SFG | The homolog of node 1 | - |
| 17 | -41.9 | 13.6 | 36.4 | MFG | Phonology, Semantics | Semantic. Monitor/Discrimination, Word Generation (Covert) |
| 18 | 42.2 | 11.9 | 38.4 | MFG | The homolog of node 17 | - |
| 29 | -45.6 | 13.0 | 23.4 | IFG | Phonology, Semantics, Speech, and Syntax | Phonological. Discrimination, Semantic. Monitor/Discrimination |
| 30 | 45.0 | 15.7 | 25.0 | IFG | The homolog of node 29 | - |
| 31 | -47.6 | 31.6 | 13.6 | IFG | Phonology, Semantics, Speech, and Syntax | Phonological. Discrimination, Semantic. Monitor/Discrimination, Word Generation (Covert and Overt) |
| 32 | 47.8 | 35.1 | 13.2 | IFG | The homolog of node 31 | - |
| 33 | -52.2 | 22.5 | 11.4 | IFG | Semantics, Speech, and Syntax | Reading (Covert), Semantic. Monitor/Discrimination, Word Generation (Covert and Overt) |
| 34 | 54.2 | 23.9 | 11.8 | IFG | The homolog of node 33 | - |
| 35 | -49.1 | 36.3 | -2.8 | IFG | Semantics, Speech, and Syntax | Semantic. Monitor/Discrimination, Word Generation (Covert) |
| 36 | 50.8 | 36.8 | -0.7 | IFG | The homolog of node 35 | - |
| 37 | -39.5 | 22.9 | 3.7 | IFG | Phonology, Semantics, Speech, and Syntax | Semantic. Monitor/Discrimination, Word Generation (Covert) |
| 38 | 42.1 | 22.0 | 3.2 | IFG | The homolog of node 37 | - |
| 39 | -51.3 | 13.2 | 6.0 | IFG | Phonology, Semantics, Speech | Music. Comprehension/Production, Recitation/Repetition. (Covert), Word Generation (Covert) |
| 40 | 53.6 | 14.3 | 11.8 | IFG | The homolog of node 39 | - |
| 53 | -49.5 | -7.1 | 38.8 | PrG | Execution. Speech | Reading (Overt), Recitation/Repetition. (Overt) |
| 54 | 54.9 | -2.0 | 33.3 | PrG | Execution. Speech | Reading (Overt), Recitation/Repetition. (Overt) |
| 63 | -49.1 | 4.7 | 30.5 | PrG | Orthography, Phonology, Semantics, Speech, and Syntax | Phonological. Discrimination, Reading (Covert) |
| 64 | 51.1 | 7.2 | 30.9 | PrG | The homolog of node 63 | - |
| 71 | -53.7 | -32.1 | 12.4 | STG | Execution. Speech, Phonology, and Speech | Music. Comprehension/Production, Passive Listening, Phonological. Discrimination, Reading (Overt), Recitation/Repetition. (Covert and Overt) |
| 72 | 54.5 | -23.7 | 10.6 | STG | Execution. Speech, Phonology | Music. Comprehension/Production, Passive Listening, Phonological. Discrimination, Recitation/Repetition. (Overt) |
| 73 | -50.1 | -10.3 | 1.1 | STG | Execution. Speech, Phonology, Speech | Music. Comprehension/Production, Passive Listening, Phonological. Discrimination, Reading (Overt), Recitation/Repetition. (Overt) |
| 74 | 51.1 | -3.7 | -0.9 | STG | Execution. Speech, | Music. Comprehension/Production, Passive Listening, Recitation/Repetition. (Overt) |
| 75 | -62.8 | -32.8 | 7.4 | STG | Execution. Speech, Phonology, Semantics, Speech | Passive Listening, Phonological. Discrimination, Reading (Overt), Semantic. Monitor/Discrimination |
| 76 | 66.5 | -20.8 | 6.6 | STG | Execution. Speech, Phonology, Speech | Music. Comprehension/Production, Passive Listening, Phonological. Discrimination, Reading (Overt) |
| 77 | -45.1 | 10.5 | -19.4 | STG | The homolog of node 78 | - |
| 78 | 47.1 | 12.3 | -19.7 | STG | Speech | Film Viewing, Passive Listening |
| 79 | -55.1 | -3.2 | -10.1 | STG | Execution. Speech, Phonology, Semantics, Speech | Music. Comprehension/Production, Passive Listening, Phonological. Discrimination, Reading (Overt) |
| 80 | 55.8 | -12.5 | -5.2 | STG | Execution. Speech, Phonology, Semantics, Speech | Music. Comprehension/Production, Passive Listening, Phonological. Discrimination, Reading (Overt), Semantic. Monitor/Discrimination |
| 81 | -65.2 | -30.9 | -11.3 | MTG | Semantics | Semantic. Monitor/Discrimination |
| 82 | 64.5 | -29.2 | -13.2 | MTG | The homolog of node 81 | - |
| 83 | -53.2 | 2.2 | -29.6 | MTG | Language | Semantic. Monitor/Discrimination, Reading (Covert) |
| 84 | 51.1 | 5.7 | -31.8 | MTG | Language | Passive Listening |
| 85 | -58.9 | -57.6 | 4.3 | MTG | Semantics and Syntax | Semantic. Monitor/Discrimination, Word Generation (Overt) |
| 86 | 60.1 | -53.3 | 2.9 | MTG | The homolog of node 85 | Film Viewing |
| 87 | -58.5 | -19.8 | -9.4 | MTG | Phonology, Semantics, Speech, and Syntax | Passive Listening, Phonological. Discrimination, Reading (Covert), Semantic. Monitor/Discrimination |
| 88 | 58.3 | -15.4 | -10.1 | MTG | Semantics and Speech | Passive Listening, Phonological. Discrimination |
| 89 | -45.5 | -26.7 | -26.1 | ITG | Orthography and Semantics | Reading (Covert), Semantic. Monitor/Discrimination |
| 90 | 45.8 | -14.6 | -32.4 | ITG | The homolog of node 89 | - |
| 91 | -50.5 | -57.0 | -14.1 | ITG | Phonology and Semantics | Naming (Overt) |
| 92 | 53.5 | -52.4 | -18.5 | ITG | Phonology and Semantics | - |
| 93 | -43.7 | -2.9 | -41.4 | ITG | Semantics | Semantic. Monitor/Discrimination |
| 94 | 40.5 | -2.9 | -41.4 | ITG | The homolog of node 93 | - |
| 97 | -55.2 | -60.3 | -6.0 | ITG | Semantics and Speech | Film Viewing, Naming (Overt) |
| 98 | 54.2 | -56.9 | -8.6 | ITG | The homolog of node 97 | - |
| 99 | -58.8 | -42.1 | -16.0 | ITG | Orthography and Semantics | Naming (Overt), Reading (Covert), Semantic. Monitor/Discrimination, Word Generation (Overt) |
| 100 | 60.5 | -40.5 | -17.1 | ITG | The homolog of node 99 | - |
| 101 | -54.9 | -30.3 | -27.4 | ITG | Semantics | - |
| 102 | 53.7 | -30.3 | -26.3 | ITG | The homolog of node 101 | - |
| 103 | -32.4 | -16.6 | -32.3 | FuG | Semantics and Speech | Naming (Overt), Semantic. Monitor/Discrimination |
| 104 | 33.1 | -14.6 | -34.1 | FuG | Semantics | Naming (Overt), Semantic. Monitor/Discrimination |
| 105 | -30.6 | -64.4 | -14.1 | FuG | Orthography, Semantics, Speech | Naming (Covert and Overt) |
| 106 | 31.3 | -61.4 | -13.7 | FuG | Language | Naming (Covert and Overt) |
| 107 | -42.3 | -50.9 | -17.3 | FuG | Orthography, Phonology, Semantics, Speech | Naming (Covert and Overt), Phonological. Discrimination, Reading (Covert), and Semantic. Monitor/Discrimination |
| 108 | 42.7 | -49.1 | -18.6 | FuG | Orthography, Semantics | Naming (Covert) |
| 113 | -28.3 | -32.6 | -16.9 | PhG | Semantics | Naming (Overt), Semantic. Monitor/Discrimination |
| 114 | 28.8 | -30.7 | -17.5 | PhG | The homolog of node 113 | Passive Listening, Semantic. Monitor/Discrimination |
| 121 | -54.4 | -39.8 | 4.2 | pSTS | Phonology, Semantics, Speech, Syntax | Passive Listening, Phonological. Discrimination, Reading (Covert), Semantic. Monitor/Discrimination, Word Generation (Covert and Overt) |
| 122 | 52.9 | -36.8 | 3.1 | pSTS | Execution. Speech, Phonology, Semantics, and Speech | Passive Listening, Phonological. Discrimination, Reading (Overt) |
| 123 | -52.4 | -50.3 | 10.8 | pSTS | Orthography, Semantics, Speech, and Syntax | Reading (Covert), Semantic. Monitor/Discrimination |
| 124 | 56.5 | -40.1 | 12.5 | pSTS | The homolog of node 123 | Passive Listening |
| 143 | -46.8 | -64.7 | 25.8 | IPL | Language | Semantic. Monitor/Discrimination, |
| 144 | 53.0 | -54.1 | 24.4 | IPL | The homolog of node 143 | - |
| 155 | -50.4 | -15.8 | 42.1 | PoG | Execution. Speech | Recitation/Repetition. (Overt) |
| 156 | 50.3 | -14.2 | 43.7 | PoG | Execution. Speech | Reading (Overt), Recitation/Repetition. (Overt) |
| 157 | -55.8 | -14.0 | 16.2 | PoG | Execution. Speech | Recitation/Repetition. (Overt) |
| 158 | 55.2 | -10.2 | 15.0 | PoG | Execution. Speech | Recitation/Repetition. (Overt) |

*SFG*, superior frontal gyrus; *MFG*, middle frontal gyrus; *IFG*, inferior frontal gyrus; *OrG*, orbital gyrus; *PrG*, precentral gyrus; *STG*, superior temporal gyrus; *MTG*, middle temporal gyrus; *ITG*, inferior temporal gyrus; *FuG*, fusiform gyrus; *PhG*, hippocampal gyrus; *pSTS*, posterior superior temporal sulcus; *IPL*, inferior parietal lobule. *PoG*, postcentral gyrus. The ID is the number of the parcel in the Brainnetome atlas (Fan et al., 2016). Here we only summarized the language-related behavioral domains and paradigm classes; the full behavioral domains and paradigm classes for each node are available at http://atlas.brainnetome.org/bnatlas.php.

Supplementary Table 3. MRI scan parameters in CCNP cohort.

|  | CKG sample | PEK sample |
| --- | --- | --- |
| TR (ms) | 2500 | 2000 |
| TE (ms) | 30 | 30 |
| FA | 80° | 90° |
| Matrix | 72 × 72 | 64 × 64 |
| Number of slices | 38 | 33 |
| Number of volumes | 184 | 180 |
| Voxel size (mm) | 3 × 3 × 3.33 | 3.5 × 3.5 × 4.2 |

Supplementary Table 4. MRI scan parameters in Southwest University cohort.

|  | 5-12 | 13-18 | 19-60 |
| --- | --- | --- | --- |
| Scanner | Siemens TrioTim | Siemens Prisma_fit | Siemens TrioTim |
| TR (ms) | 2000 | 1000 | 2000 |
| TE (ms) | 30 | 30 | 30 |
| FA | 90° | 73° | 90° |
| Matrix | 64 × 64 | 78 × 78 | 64 × 64 |
| Number of slices | 33 | 56 | 32 |
| Number of volumes | 180 | 480 | 242 |
| Voxel size (mm) | 3.13 × 3.13 × 4.6 | 2.5 × 2.5 × 2.5 | 3.44 × 3.44 × 4 |

Supplementary Table 5. Partial correlation of global topological characteristics with behavior

|  |  |  | VIQ | | ADOS_COMM | | ADOS_SOC | | ADOS_RRB | |
| --- | --- | --- | --- | --- | --- | --- | --- | --- | --- | --- |
|  |  |  | R | *P* | R | *P* | R | *P* | R | *P* |
| Children | State 1 | FC Str | -0.02 | 0.8 | -0.05 | 0.66 | 0.01 | 0.9 | 0.03 | 0.81 |
|  |  | gE | -0.1 | 0.24 | -0.08 | 0.44 | 0.09 | 0.39 | -0.06 | 0.64 |
|  |  | lE | -0.11 | 0.21 | -0.08 | 0.46 | 0.08 | 0.43 | -0.09 | 0.46 |
|  | State 2 | FC Str | 0.16 | 0.06 | -0.12 | 0.29 | -0.07 | 0.51 | -0.09 | 0.48 |
|  |  | gE | -0.13 | 0.13 | -0.1 | 0.36 | 0.08 | 0.44 | -0.02 | 0.85 |
|  |  | lE | **-0.19** | **0.02** | -0.09 | 0.4 | 0.05 | 0.62 | 0.01 | 0.93 |
|  | State 3 | FC Str | -0.07 | 0.44 | -0.07 | 0.53 | 0.07 | 0.53 | 0.05 | 0.66 |
|  |  | gE | -0.01 | 0.9 | -0.13 | 0.23 | 0.03 | 0.81 | -0.14 | 0.24 |
|  |  | lE | -0.05 | 0.56 | -0.13 | 0.23 | 0.01 | 0.92 | -0.12 | 0.31 |
|  | State 4 | FC Str | -0.02 | 0.85 | -0.06 | 0.56 | 0.07 | 0.53 | -0.01 | 0.95 |
|  |  | gE | -0.12 | 0.16 | -0.13 | 0.24 | 0.08 | 0.48 | -0.03 | 0.8 |
|  |  | lE | -0.15 | 0.07 | -0.14 | 0.19 | 0.09 | 0.4 | -0.01 | 0.9 |
| Adolescents | State 1 | FC Str | 0.08 | 0.25 | 0.07 | 0.51 | 0.06 | 0.5 | 0.09 | 0.48 |
|  |  | gE | -0.07 | 0.32 | -0.16 | 0.1 | -0.1 | 0.32 | 0.12 | 0.31 |
|  |  | lE | -0.06 | 0.45 | -0.18 | 0.07 | -0.16 | 0.1 | 0.04 | 0.75 |
|  | State 2 | FC Str | 0.05 | 0.46 | 0.01 | 0.93 | 0.05 | 0.57 | -0.03 | 0.78 |
|  |  | gE | -0.06 | 0.41 | -0.03 | 0.73 | 0.01 | 0.88 | -0.01 | 0.92 |
|  |  | lE | -0.01 | 0.88 | -0.02 | 0.81 | 0.04 | 0.71 | -0.04 | 0.74 |
|  | State 3 | FC Str | 0.03 | 0.67 | 0.1 | 0.29 | 0.14 | 0.15 | 0.11 | 0.36 |
|  |  | gE | 0.05 | 0.51 | -0.14 | 0.14 | -0.03 | 0.78 | 0.08 | 0.51 |
|  |  | lE | 0.04 | 0.6 | -0.07 | 0.47 | 0.04 | 0.69 | 0.1 | 0.4 |
|  | State 4 | FC Str | 0.04 | 0.57 | -0.02 | 0.81 | 0.08 | 0.41 | 0.08 | 0.52 |
|  |  | gE | 0.02 | 0.76 | -0.12 | 0.2 | -0.11 | 0.24 | 0.01 | 0.93 |
|  |  | lE | 0.04 | 0.57 | -0.13 | 0.18 | -0.14 | 0.16 | -0.03 | 0.8 |
| Adults | State 1 | FC Str | 0.04 | 0.69 | **-0.31** | **< 0.01** | **-0.3** | **< 0.01** | -0.1 | 0.43 |
|  |  | gE | -0.14 | 0.17 | 0.16 | 0.13 | 0.11 | 0.3 | -0.01 | 0.94 |
|  |  | lE | **-0.21** | **0.04** | **0.25** | **0.01** | 0.18 | 0.08 | -0.08 | 0.51 |
|  | State 2 | FC Str | -0.04 | 0.67 | **-0.3** | **< 0.01** | -0.17 | 0.09 | -0.21 | 0.08 |
|  |  | gE | **-0.37** | **< 0.01** | **0.25** | **0.01** | **0.3** | **< 0.01** | 0.01 | 0.95 |
|  |  | lE | **-0.44** | **< 0.01** | **0.35** | **< 0.01** | **0.3** | **< 0.01** | 0.05 | 0.71 |
|  | State 3 | FC Str | -0.03 | 0.04 | -0.14 | 0.18 | -0.05 | 0.62 | 0.06 | 0.63 |
|  |  | gE | -**0.22** | **0.03** | 0.12 | 0.22 | **0.25** | **0.01** | -0.19 | 0.13 |
|  |  | lE | -**0.33** | **< 0.01** | **0.32** | **< 0.01** | **0.31** | **< 0.01** | -0.1 | 0.43 |
|  | State 4 | FC Str | **-0.2** | **0.05** | -0.02 | 0.85 | -0.01 | 0.89 | -0.07 | 0.56 |
|  |  | gE | **0.36** | **< 0.01** | **-0.41** | **< 0.01** | **-0.24** | **0.02** | -0.01 | 0.95 |
|  |  | lE | **0.33** | **< 0.01** | **-0.41** | **< 0.01** | **-0.24** | **0.02** | 0.01 | 0.97 |
| All | State 1 | FC Str | 0.02 | 0.62 | -0.07 | 0.23 | -0.06 | 0.33 | 0.01 | 0.91 |
|  |  | gE | -0.07 | 0.15 | -0.04 | 0.52 | 0.03 | 0.62 | 0.02 | 0.75 |
|  |  | lE | -0.08 | 0.08 | < 0.01 | 0.99 | 0.03 | 0.6 | -0.03 | 0.63 |
|  | State 2 | FC Str | 0.06 | 0.24 | -0.11 | 0.06 | -0.05 | 0.37 | -0.11 | 0.11 |
|  |  | gE | **-0.12** | **0.01** | -0.01 | 0.87 | 0.1 | 0.09 | -0.01 | 0.86 |
|  |  | lE | **-0.15** | **< 0.01** | 0.03 | 0.55 | 0.1 | 0.08 | 0.01 | 0.92 |
|  | State 3 | FC Str | -0.02 | 0.67 | 0.01 | 0.93 | 0.07 | 0.25 | 0.05 | 0.49 |
|  |  | gE | 0.01 | 0.9 | -0.1 | 0.1 | 0.04 | 0.53 | -0.04 | 0.61 |
|  |  | lE | -0.04 | 0.42 | -0.01 | 0.85 | 0.08 | 0.15 | -0.03 | 0.66 |
|  | State 4 | FC Str | -0.04 | 0.36 | -0.02 | 0.76 | 0.05 | 0.35 | 0.01 | 0.91 |
|  |  | gE | 0.08 | 0.08 | **-0.22** | **< 0.01** | -0.1 | 0.08 | -0.03 | 0.62 |
|  |  | lE | 0.07 | 0.15 | **-0.22** | **< 0.01** | -0.1 | 0.08 | -0.04 | 0.6 |

Supplementary Table 6. Partial correlation of nodal topological characteristics with behavior (Children)

|  | | VIQ | | | ADOS_COMM | | | ADOS_SOC | | | ADOS_RRB | | |
| --- | --- | --- | --- | --- | --- | --- | --- | --- | --- | --- | --- | --- | --- |
|  |  | Regions | R | *P* | Regions | R | *P* | Regions | R | *P* | Regions | R | *P* |
| Children | State 1 | LIFG | 0.251 | 0.003 | LMTG | 0.216 | 0.047 | LITG | -0.257 | 0.015 | LSFG | -0.313 | 0.008 |
|  |  | RIFG | -0.174 | 0.043 | RITG | -0.268 | 0.013 |  |  |  | LIFG | 0.263  0.252 | 0.028  0.035 |
|  |  | LSTG | -0.169 | 0.049 | LITG | -0.217 | 0.046 |  |  |  |  |  |  |
|  |  | LpSTS | -0.174 | 0.042 | LFuG | 0.333 | 0.002 | LPoG | 0.289 | 0.006 | RITG | 0.244 | 0.042 |
|  |  | LPoG | 0.197 | 0.022 | LPoG | 0.3 | 0.005 |  |  |  | LITG | -0.302 | 0.011 |
|  |  |  |  |  | RPoG | -0.264 | 0.015 |  |  |  | RPhG | -0.257 | 0.032 |
|  | State 2 | RMFG | 0.189 | 0.027 | RIFG | -0.224 | 0.039 | RIFG | -0.266 | 0.012 | LIFG | 0.294 | 0.013 |
|  |  | LSTG | -0.174  -0.2 | 0.043  0.02 |  |  |  |  |  |  | LMTG | 0.328 | 0.006 |
|  |  | RSTG | -0.205 | 0.017 | RITG | 0.261 | 0.016 | RITG | 0.27 | 0.011 | LITG | 0.272  -0.304 | 0.023  0.011 |
|  |  | LMTG | 0.181  -0.186 | 0.035  0.03 |  |  |  |  |  |  |  |  |  |
|  |  | RITG | 0.197  0.201 | 0.021  0.019 | RPhG | 0.237 | 0.029 | RPhG | 0.234 | 0.027 | RITG | -0.239 | 0.046 |
|  |  | LPoG | 0.277 | 0.001 | RIPL | 0.217 | 0.046 | LpSTS | -0.294 | 0.005 | RPhG | 0.251 | 0.036 |
|  |  | RPoG | -0.186 | 0.031 |  |  |  |  |  |  |  |  |  |
|  | State 3 | LMTG | 0.196  -0.17 | 0.022  0.048 | LSTG | -0.228 | 0.036 | RIFG | -0.277 | 0.009 | LIFG | 0.28 | 0.019 |
|  |  |  |  |  |  |  |  |  |  |  | RSTG | -0.251 | 0.036 |
|  |  |  |  |  |  |  |  | LIFG | -0.236 | 0.026 | LITG | -0.242  -0.258 | 0.043  0.031 |
|  |  | RFuG | 0.247 | 0.004 |  |  |  |  |  |  |  |  |  |
|  |  |  |  |  | LPoG | 0.263 | 0.015 | LITG | -0.225 | 0.034 | RFuG | 0.25 | 0.037 |
|  |  | RIPL | -0.239 | 0.005 |  |  |  |  |  |  | LpSTS | 0.415 | <0.001 |
|  |  |  |  |  |  |  |  |  |  |  | RPoG | -0.286 | 0.016 |
|  | State 4 | RIFG | 0.18 | 0.036 | RSTG | 0.225 | 0.039 | RITG | 0.216 | 0.042 | LIFG | 0.262  -0.273 | 0.028  0.022 |
|  |  | LSTG | 0.175 | 0.041 |  |  |  |  |  |  |  |  |  |
|  |  | RSTG | -0.193 | 0.025 | RITG | 0.226 | 0.038 |  |  |  | RIFG | 0.272 | 0.023 |
|  |  | RITG | 0.246 | 0.004 |  |  |  |  |  |  | RPrG | 0.257 | 0.032 |
|  |  | LITG | -0.173 | 0.045 | LPoG | 0.237 | 0.029 |  |  |  | RMTG | 0.312 | 0.009 |
|  |  | RIPL | -0.283 | <0.001 |  |  |  |  |  |  | LITG | -0.26 | 0.03 |
|  |  | LPoG | 0.176 | 0.04 |  |  |  |  |  |  | LPoG | -0.321 | 0.007 |

All *P* values are uncorrected.

Supplementary Table 7. Partial correlation of nodal topological characteristics with behavior (Adolescents)

|  | | VIQ | | | ADOS_COMM | | | ADOS_SOC | | | ADOS_RRB | | |
| --- | --- | --- | --- | --- | --- | --- | --- | --- | --- | --- | --- | --- | --- |
|  |  | Regions | R | *P* | Regions | R | *P* | Regions | R | *P* | Regions | R | *P* |
| Adolescents | State 1 |  | | | LSTG | -0.195 | 0.044 | LPrG | 0.201 | 0.037 | LIFG | -0.252 | 0.037 |
|  |  |  |  |  | LPoG | 0.24 | 0.013 | RpSTS | 0.199 | 0.038 |  |  |  |
|  |  |  |  |  |  |  |  | LPoG | 0.223 | 0.02 |  |  |  |
|  | State 2 | LMTG | 0.191 | 0.008 | RIFG | -0.193 | 0.046 | RMTG | -0.221 | 0.021 | LITG | -0.239 | 0.048 |
|  |  |  |  |  | LPrG | 0.227 | 0.019 |  |  |  |  |  |  |
|  |  |  |  |  | RPrG | 0.208 | 0.032 |  |  |  |  |  |  |
|  |  |  |  |  | LSTG | 0.247 | 0.01 |  |  |  |  |  |  |
|  |  | RITG | 0.213 | 0.003 | LITG | -0.196 | 0.044 | RITG | -0.211 | 0.028 |  |  |  |
|  |  |  |  |  | RITG | 0.219 | 0.023 |  |  |  |  |  |  |
|  |  |  |  |  | RpSTS | 0.335 | <0.001 |  |  |  |  |  |  |
|  | State 3 | RMTG | 0.152 | 0.036 | LMTG | 0.192 | 0.048 | LIFG | 0.226 | 0.018 | LITG | 0.29 | 0.016 |
|  |  |  |  |  | LITG | -0.201 | 0.033 |  |  |  |  |  |  |
|  |  | LPoG | -0.201 | 0.005 | RITG | 0.252 | 0.009 | RPrG | 0.211 | 0.027 |  |  |  |
|  |  |  |  |  | LIPL | 0.254 | 0.008 |  |  |  |  |  |  |
|  |  |  |  |  | RPoG | 0.206 | 0.033 | LIPL | 0.223 | 0.02 |  |  |  |
|  | State 4 | LSTG | -0.153 | 0.034 | LITG | -0.235 | 0.015 | RIFG | 0.191 | 0.047 | RpSTS | 0.261 | 0.03 |
|  |  | RSTG | -0.178 | 0.013 |  |  |  | LIFG | 0.2 | 0.037 |  |  |  |
|  |  | RMTG | 0.173 | 0.017 |  |  |  | LPrG | 0.231 | 0.016 |  |  |  |
|  |  | LFuG | 0.144 | 0.047 | RITG | 0.194 | 0.045 | RFuG | 0.251  0.205 | 0.009  0.032 |  |  |  |

All *P* values are uncorrected.

Supplementary Table 8. Partial correlation of nodal topological characteristics with behavior (Adults)

|  | | VIQ | | | ADOS_COMM | | | ADOS_SOC | | | ADOS_RRB | | |
| --- | --- | --- | --- | --- | --- | --- | --- | --- | --- | --- | --- | --- | --- |
|  |  | Regions | R | *P* | Regions | R | *P* | Regions | R | *P* | Regions | R | *P* |
| Adults | State 1 | LSFG | -0.282 | 0.006 | RITG | 0.216  0.265 | 0.034  0.009 | LITG | 0.227 | 0.025 | LMTG | 0.262  0.321 | 0.031  0.008 |
|  |  | LMTG | -0.253 | 0.014 |  |  |  |  |  |  |  |  |  |
|  |  | LITG | -0.206  -0.219 | 0.046  0.034 |  |  |  |  |  |  | RITG | 0.338 | 0.005 |
|  |  |  |  |  | RpSTS | 0.207 | 0.042 | LPoG | 0.225 | 0.027 |  |  |  |
|  |  | RITG | -0.214 | 0.039 |  |  |  |  |  |  | LITG | 0.448 | <0.001 |
|  |  | LPoG | -0.206 | 0.046 |  |  |  |  |  |  | RFuG | 0.266 | 0.028 |
|  | State 2 | RSFG | -0.242 | 0.019 | RSFG | 0.228 | 0.025 | RSFG | 0.316 | 0.002 | LSTG | 0.476 | <0.001 |
|  |  |  |  |  | RIFG | 0.209 | 0.04 |  |  |  |  |  |  |
|  |  |  |  |  | LPrG | 0.241 | 0.017 | LpSTS | 0.228 | 0.025 |  |  |  |
|  |  | RITG | -0.316 | 0.002 | LITG | 0.247 | 0.015 | RpSTS | 0.316 | 0.002 | RpSTS | 0.584 | <0.001 |
|  |  |  |  |  | RpSTS | 0.264 | 0.009 | LPoG | 0.224 | 0.028 |  |  |  |
|  | State 3 | LMFG | 0.219 | 0.034 | LSTG | 0.205 | 0.044 | LIFG | 0.255 | 0.012 | RMTG | 0.314 | 0.009 |
|  |  |  |  |  | RSTG | 0.245 | 0.016 |  |  |  |  |  |  |
|  |  |  |  |  | RMTG | 0.204 | 0.045 | RPrG | 0.237 | 0.019 |  |  |  |
|  |  | LSTG | -0.232 | 0.024 | LMTG | 0.281 | 0.005 |  |  |  |  |  |  |
|  |  |  |  |  | LITG | 0.283 | 0.005 | LMTG | 0.218 | 0.032 | RFuG | 0.298 | 0.014 |
|  |  |  |  |  | RITG | 0.214 | 0.035 | LITG | 0.33  0.239 | <0.001  0.019 |  |  |  |
|  |  | LPoG | -0.203 | 0.049 | RFuG | 0.241 | 0.017 |  |  |  |  |  |  |
|  |  |  |  |  | LPoG | 0.201 | 0.049 | RFuG | 0.254 | 0.012 |  |  |  |
|  | State 4 | LITG | -0.217 | 0.036 | RMFG | 0.314 | 0.002 | RMFG | 0.243 | 0.016 | RIFG | 0.245 | 0.045 |
|  |  |  |  |  | RIFG | 0.209 | 0.04 |  |  |  | RIFG | 0.263 | 0.031 |
|  |  | LFuG | -0.207 | 0.045 | RSTG | 0.218  0.231 | 0.032  0.023 | LIFG | -0.253 | 0.013 | RSTG | -0.3 | 0.013 |
|  |  |  |  |  | LITG | 0.254  0.227 | 0.012  0.026 | RIFG | 0.202 | 0.047 |  |  |  |
|  |  |  |  |  |  |  |  | RITG | 0.213 | 0.037 |  |  |  |
|  |  | LPoG | -0.229 | 0.026 | 52 | 0.253 | 0.013 | RFuG | 0.22 | 0.03 | LFuG | 0.309 | 0.01 |
|  |  |  |  |  | 60 | 0.239 | 0.019 |  |  |  |  |  |  |
|  |  |  |  |  | 63 | 0.243 | 0.017 |  |  |  |  |  |  |
|  |  |  |  |  | 64 | 0.324 | 0.001 |  |  |  |  |  |  |

All *P* values are uncorrected.

**References**

Achard, S., Salvador, R., Whitcher, B., Suckling, J., & Bullmore, E. (2006). A resilient, low-frequency, small-world human brain functional network with highly connected association cortical hubs. *J Neurosci, 26*(1), 63-72. doi:10.1523/JNEUROSCI.3874-05.2006

Dockes, J., Poldrack, R. A., Primet, R., Gozukan, H., Yarkoni, T., Suchanek, F., . . . Varoquaux, G. (2020). NeuroQuery, comprehensive meta-analysis of human brain mapping. *Elife, 9*. doi:10.7554/eLife.53385

Gong, G., He, Y., Concha, L., Lebel, C., Gross, D. W., Evans, A. C., & Beaulieu, C. (2009). Mapping anatomical connectivity patterns of human cerebral cortex using in vivo diffusion tensor imaging tractography. *Cereb Cortex, 19*(3), 524-536. doi:10.1093/cercor/bhn102

He, Y., Wang, J., Wang, L., Chen, Z. J., Yan, C., Yang, H., . . . Evans, A. C. (2009). Uncovering intrinsic modular organization of spontaneous brain activity in humans. *PLoS One, 4*(4), e5226. doi:10.1371/journal.pone.0005226
